# Multimodal Imaging of the Cellular and Extracellular Microenvironment on the Same Formalin-Fixed Paraffin-Embedded Tissue Section

**DOI:** 10.64898/2026.08.21.746293

**Authors:** Jade K. Macdonald, Thai Pham, Alan J. Simmons, Harsimran Kaur, Jamie L. Allen, Anna J. Smith, Audra M. Judd, Seung Woo Kang, Madeline E. Colley, Melissa A. Farrow, Ken S. Lau, Jeffrey M. Spraggins

**Author notes:** Corresponding Author Jeffrey M Spraggins.

## Abstract

Same-tissue section multimodal imaging is a powerful strategy that spatially profiles tissue histology, cell populations and molecular composition while maximizing tissue economy, preserving spatial molecular relationships, and increasing co-registration capacity. However, performing multiple modalities on the same tissue section can destroy or chemically alter the tissue, compromising downstream data. Here, we systematically assess integration of picrosirius red staining, hematoxylin and eosin staining, and multiplexed immunofluorescence into N-glycan and extracellular matrix peptide matrix-assisted laser/desorption ionization imaging mass spectrometry (IMS) workflows. We evaluate alterations in tissue morphology, stain efficiency, IMS feature intensity as well as IMS feature localization after upstream modality integration. We propose an optimized multimodal sequence that maximizes data quality and follows a very specific order of: autofluorescence microscopy, multiplexed immunofluorescence, picrosirius red staining, N-glycan IMS, hematoxylin and eosin staining, and extracellular matrix peptide IMS. Overall, this work develops an optimized multimodal workflow that comprehensively images tissue morphology, collagen fibers, and cell populations at single-cell resolution as well as multiplexed N-glycan composition and multiplexed extracellular matrix peptides with post-translational modification status from a single tissue section.

## INTRODUCTION

Multimodal tissue imaging has become an increasingly powerful strategy to build a comprehensive molecular and histological picture of complex tissue biology^1–4^. Same-tissue section multimodal workflows integrate structural, molecular, and histological information, providing additional biological context for each modality while optimizing tissue economy, preserving spatial relationships, and improving co-registration capabilities that are oftentimes compromised when using serial sections ^5–7^. This is particularly valuable when studying tissue microenvironments that include the extracellular matrix (ECM), where structural remodeling, protein composition, post-translational modifications and immune cell recruitment are known to change in concert during pathological processes^8,9^.

Several imaging modalities are commonly used to image the collagenous extracellular matrix^10^. Autofluorescence (AF) microscopy provides a rapid, label-free overview of tissue architecture and is frequently used for co-registration of modalities^11^. Chemical stains provide information on tissue morphology at single-cell resolution by binding to cellular or extracellular components based on their chemical properties. Hematoxylin and eosin (H&E) stains cellular and morphological tissue architecture that guides pathologist annotation and diagnoses^12^. This histological framework is essential for co-localizing downstream molecular imaging modality features to defined anatomical and pathological structures. Picrosirius red (PSR) stain is commonly used to image the extracellular matrix^13^. When imaged with brightfield light, PSR stains the extracellular matrix red. When imaged with polarized light, PSR stain images collagen fiber width and organization, a key structural component of the extracellular matrix that has been implicated in disease^14–17^. Multiplexed immunofluorescence (MxIF) enables spatially resolved detection of up to 60 protein markers within a single tissue section, allowing simultaneous identification of immune, epithelial, and stromal cell populations that facilitate characterization of ECM-directed cell recruitment and cell-ECM crosstalk^18^. Imaging mass spectrometry (IMS) provides highly multiplexed, direct detection of molecular analytes^19^. N-glycan IMS workflows enzymatically release and report the spatial distribution of hundreds of N-linked glycans, which localize to histological features as well as change composition based on disease state^20^. ECM-directed IMS workflows rely on enzymatic digestion of collagens and collagen-like proteins, providing spatially resolved detection of collagenase-generated peptides and their post-translational modifications^21^.

While each of these modalities has been performed individually on formalin-fixed paraffin-embedded tissue, combining modalities on a single tissue section introduces technical complexities. Some of these techniques are destructive or chemically modifying. Antibody incubation, chemical stains, and enzymatic digests can alter tissue integrity, epitope availability, or chemical environment in ways that could affect performance of downstream modalities. As a result, the order in which these modalities are performed can determine the overall data quality obtainable from a single tissue section. Here, we develop a multimodal imaging workflow that integrates AF, MxIF, H&E stain, PSR stain, N-glycan IMS and ECM IMS on a single tissue section. We systematically compare integration of each modality within an enzymatic IMS workflow^22^. We propose a multimodal imaging workflow that optimizes data quality while providing a practical framework for maximizing the amount of co-registered molecular, histological and structural data that can be collected from a single tissue section.

## MATERIALS AND METHODS

### Materials

Ammonium bicarbonate (A6141), dimethyl sulfoxide (472301), eosin-y (HT110116), Mayer’s hematoxylin (MHS16), nuclease-free water (W4502), sodium citrate (S461), Trizma base (T1503), xylenes (534056), α-cyano-4-hydroxycinnamic acid (C2020), were purchased from Sigma Aldrich (St. Louis, MO). LC-MS-grade acetonitrile (1.000292500), LC-MS-grade water (1153332500), and ZipTips (ZTC18M096) were purchased from Millipore Sigma (Burlington, MA). Calcium chloride (089866. 18), formic acid (28905), picrosirius red staining kit (NC9039835) and Weigert’s hematoxylin staining kit (NC0856208) were purchased from Fisher Scientific (Waltham, MA).

Histology-grade acetonitrile (A998-4) and trifluoroacetic acid (A116 10x1AMP) were purchased from Fisher Chemical (Waltham, MA). Ethanol (2701) was purchased from Decon Laboratories, Inc (Prussia, PA). Histochoice Clearing Agent (H103) was purchased from VWR (Radnor, PA).

Bluing reagent (CS70230-2) was purchased from Agilent Technologies (Santa Clara, CA). Multiplexed immunofluorescence consumables (antibody conjugation kits, staining kits, reporters, flow cells, 10x Phenocylcer buffer solution (7000001) and 10x phenocylcer buffer additive (240257)) were purchased from Quaterix Corp (Billerica, MA). PNGase F prime was purchased from Bulldog Bio (Portsworth, NH). Collagenase Type 3 was purchased from Worthington Scientific (Lakewood, NJ). Bio iRT peptides (1816351) were purchased from Bruker Daltonics (Billerica, MA).

### Tissue acquisition, ethics approval and tissue processing

Tissue samples were provided de-identified by the National Cancer Institute Cooperative Human Tissue Network. Ethical oversight was provided by the Institutional Review Board #182138 of Vanderbilt University. The study has been carried out in accordance with the Code of Ethics of the World Medical Association (Declaration of Helsinki). A formalin-fixed paraffin-embedded malignant colorectal cancer tissue was sectioned at 6 µm onto indium tin oxide (ITO) coated slides and heated at 55 °C overnight to ensure tissue adherence. Tissues were dewaxed and hydrated with Histochoice (2x), 100% ethanol (2x), 95% ethanol, 70% ethanol, and water (2x) prior to being subjected to each multimodal analysis.

### Autofluorescence microscopy

Initial autofluorescence images were taken with DAPI (λ = 385, 75% brightness, 60% intensity, 6.1 ms exposure time), GFP (λ = 475, 75% brightness, 60% intensity, 50 ms exposure time) and DsRed (λ = 555, 75% brightness, 60% intensity, 100 ms exposure time) filters and a 10x objective on a Zeiss AxioScan.Z1 slide scanner (Carl Zeiss Microscopy GmbH, Oberkochen, Germany).

Autofluorescence images taken after imaging mass spectrometry had altered settings for the DAPI channel (λ = 385, 100% brightness, 30% intensity, 1.0 ms exposure time) due to presence of α-cyano-4-hydroxycinnamic acid matrix.

### Chemical staining

For both staining procedures, tissues were first rehydrated as follows: 100% ethanol (2x), 95% ethanol (1x), 70% ethanol (1x), water (2x). For picrosirius red staining, tissues were stained as follows: Weigert’s hematoxylin (8 m, 1x), water (3x), picrosirius stain solution A (2 m, 1x), water (1x), picrosirius stain kit solution B (60 m, 1x), picrosirius stain kit solution C (2 m, 1x), 70% ethanol (1x), 100% ethanol (3x), xylenes (2x) followed by administration of a coverslip. For hematoxylin and eosin staining, samples were stained after rehydration as follows: Mayer’s hematoxylin (2m), water (3x), Bluing Buffer (1m), water (3x), 70% ethanol (1x), 95% ethanol (1x), eosin (1m), 95% ethanol (20 dips), 95% ethanol (10s), 100% ethanol (2x), xylenes (2x), followed by administration of a coverslip. For all staining comparisons, staining was performed at the same time with the same solutions with the slides in the same slide dipper to ensure any differences were due to the tissue rather than experimental variations.

### Multiplexed immunofluorescence

A Phenocyler Fusion 2.0 system (Quaterix Corp, Billerica, MA) was used for multiplexed immunofluorescence and samples were prepared as recommended per the manufacturer’s guidelines^18^. In short, tissues were de-waxed, subjected to antigen retrieval (110 °C, 20 m), stained with a 35-antibody cocktail (3 hr), fixed and cyclically imaged via 12 sequential cycles of DNA reporter probe (n = 35 antibodies, 3 per cycle) hybridization and wash steps (**Supplemental Table 1**). After data acquisition, tissues were incubated in Histochoice overnight to remove the flow cell.

### N-glycan matrix-assisted laser/desorption ionization imaging mass spectrometry

Tissues were prepared for N-glycan matrix-assisted laser/desorption ionization (MALDI) IMS as follows^20^. Tissues were hydrated with decreasing amount of ethanol (100%, 95%, 70%, 0%) prior to antigen retrieval (10 mM sodium citrate, pH = 6, 95 °C, 20 m). Antigen retrieval functioned to not only reveal epitopes, but also remove picrosirius red stain or fixed MxIF antibodies when applicable. PNGase F Prime (0.2 mg/mL) was administered with an M5 Sprayer (HTX Imaging, Chapel Hill, NC) with the following parameters: 30 °C nozzle temperature, 8 passes, 10 μL/min, 1200 mm/min, 2 mm track pattern, CC pattern. Tissues were incubated in a humidity chamber for 2 hours (37 °C, 100% humidity) to facilitate N-glycan cleavage. CHCA (α-cyano-4-hydroxycinnamic acid, 7 mg/mL, 70% acetonitrile, 0.1% TFA) was administered with an M5 Sprayer. PSR comparison data was acquired on a timsTOF FleX M2 (Bruker Daltonics, Bremen, Germany) in positive ion mode, with 80 μm step size, 300 shots per pixel, 16 μm scan range for a resulting field size of 20 μm, 22 eV ion energy, 20 eV collision energy, 30 μs pre pulse storage, 120 μs transfer time, 700-4000 mass range. MxIF comparison data was acquired on a timsTOF FleX (Bruker Daltonics, Bremen, Germany) with 40 μm step size, 10 eV ion energy, 10 eV collision energy, 22 μs pre pulse storage, 150 μs transfer time and a 500-4000 mass range. After data acquisition, the imaged slides were subjected to autofluorescence microscopy to image laser ablation marks in the CHCA matrix for downstream pixel-level co-registration. N-glycans were removed with decreasing amounts of ethanol as previously described prior to any subsequent modalities^23^.

### Extracellular matrix matrix-assisted laser/desorption ionization imaging mass spectrometry

Tissues were prepared for extracellular matrix MALDI IMS as follows^21^. Tissues were hydrated prior to antigen retrieval (10 mM Tris buffer, pH = 9, 95 °C, 20 m). Collagenase (0.2 mg/mL) was applied with an M5 Sprayer (HTX Imaging, Chapel Hill, NC) with the following parameters: 40 °C nozzle temperature, 8 psi pressure, 25 μL/min flow rate, 1200 velocity, 2 mm track spacing, 8 passes, CC pattern. Tissues were incubated in a humidity chamber for 5 hours (37 °C, 100% humidity). CHCA (α-cyano-4-hydroxycinnamic acid, 7mg/mL, 50% acetonitrile, 1% TFA) was administered with an M5 Sprayer: 72 °C nozzle temperature, 8 psi pressure, 70 μL/min flow rate, 1300 velocity, 3 mm track spacing, 10 passes, CC pattern. PSR comparison data was acquired on a timsTOF FleX M2 (Bruker Daltonics, Bremen, Germany) in positive ion mode, with 80 μm step size, 600 shots per pixel, laser size of 15 μm for a resulting field size of 19 μm, 18 eV ion energy, 25 eV collision energy, 22 μs pre pulse storage, 80 μs transfer time, and a 700-2500 mass range. MxIF comparison data was acquired on a timsTOF FleX (Bruker Daltonics, Bremen, Germany) with 40 μm step size, 12 eV ion energy, 12 eV collision energy, 20 μs pre pulse storage and a 200 μs transfer time. After data acquisition, the imaged slides were subjected to autofluorescence microscopy to image laser ablation marks in the CHCA matrix for downstream co-registration to IMS pixels.

### Imaging Mass Spectrometry Feature Detection and Data Analysis

For N-glycan MALDI IMS, a peak list comprised of all m/z values from an in-house N-glycan database (n = 2642) was imported with a 20-ppm width and features with a maximum peak intensity greater than 15 (n = 239) was used for K-means segmentation analysis by Manhattan distance (k = 5). Mass to charge ratios that overlapped with second isotopes and did not have a detectable peak above background noise within the average mass spectrum were manually removed and the resulting curated N-glycan peaks were used for quantification (n = 160). For extracellular matrix imaging mass spectrometry feature detection, a threshold of 87500 counts was used to select the top 389 peaks with a 20-ppm peak width. Matrix peaks and isotope peaks were manually removed based on mass defect and spatial localization. The resulting peak list (n = 288) was used for downstream K-means segmentation analysis by Manhattan distance (k = 5) and quantification. Calibrated extracellular matrix IMS peaks were identified with a liquid chromatography tandem mass spectrometry reference library by accurate mass within 8 ppm (**Supplemental Table 2**). Maximum peak intensity mean spectrum statistics were exported with peak area interval processing was exported from SCiLS Lab v2026a for quantification. All data analysis and data visualization were performed in SCiLs Lab v2026a (Bruker Daltonics, Billerica, MA).

### Liquid chromatography trapped ion mobility tandem mass spectrometry (LC-TIMS-MS/MS)-based extracellular matrix proteomics

Liquid chromatography trapped ion mobility spectrometry tandem mass spectrometry (LC-TIMS-MS/MS) extracellular matrix proteomics was performed to generate a peptide library for MALDI- IMS peptide identification. CHCA and extracellular matrix peptides were removed with decreasing amounts of ethanol (100%, 95%, 70%, 0%) as previously described^24^. The tissue was subjected to another round of low pH antigen retrieval (sodium citrate, 10 mM, pH = 6), N-glycan removal and high pH antigen retrieval (Tris base, 10 mM, pH = 9). The next layer of extracellular matrix peptides was locally digested, and peptides were extracted with increasing amount of acetonitrile with 0.1% formic acid as previously described^24^. Peptides were isolated with ZipTip per manufacturers guidelines. Peptides were resuspended at a concentration of 200 ng/µL with 1x bio iRT peptides in 0.1% formic acid and 1 µL of sample was injected into a nanoElute connected to a timsTOF Pro2 (Bruker Daltonics, Bremen Germany). Liquid chromatography was performed at a 300 nL/min flow rate with a 15 cm C18 PepSep Column (75 μm inner diameter and 1.9 μm particle size, Bruker Daltonics, Bremen Germany) and a 30-minute gradient from 4% to 30% acetonitrile (0.1% formic acid). Data was acquired in positive ion mode, m/z = 100-1700 mass range, 0.96 cycle time, 8 PASEF ramps per cycle, 100 ms ramp time, 100% duty cycle, 60 μs transfer time, 12 μs pre pulse storage and an extended trapped ion mobility spectrometry polygon as previously described^24^. A calibration filter tuning mix with m/z = 622, 922, 1221, was used to calibrate mass (m/z value) and ion mobility. Quality control of the system was monitored with protein readouts (>3500 proteins) of a 50 ng of K562 standard after every other sample injection. Quality of the column was evaluated via retention time of bio iRT peptides. Data was searched as a nonspecific digest with using Fragpipe 24.0 against a Swiss prot reviewed database downloaded from Uniprot on March 4^th^, 2026 containing 4752 entries with search terms “collagen” OR “extracellular matrix” OR “colorectal” OR “adenoma” OR “GO:0031012” OR “GO:0005615” with previously described parameters^24^. Searched post-translational modifications include methionine oxidation (M + 15.9949), proline hydroxylation (P + 15.9949), lysine oxidative deamination (K + 1.031635) and asparagine and glutamine deamidation (NQ + 0.984016). Peptide spectrum match (PSM) output files were used for MALDI IMS peptide identification. PSMs with a hyperscore greater than 18 and peptide probability greater than 95% were considered for MALDI IMS peptide identification by accurate mass (< 8 ppm).

## RESULTS AND DISCUSSION

Previous work has shown that hematoxylin and eosin (H&E) staining followed by extracellular matrix imaging mass spectrometry (IMS) can be performed after N-glycan IMS on the same tissue section^25^. Other workflows have integrated autofluorescence microscopy as well as antibody-based single cell imaging modalities prior to IMS for complementary high-resolution feature detection^7,11,26^. However, these modalities have not been integrated with both picrosirius red (PSR) stain and multiplexed immunofluorescence (MxIF) for a comprehensive analysis of tissue morphology, collagen fibers, molecular composition and post-translational modification status of the extracellular matrix surrounding identified cell populations from the same tissue section (**Table 1**). To identify the best way to integrate PSR staining and MxIF into the above workflow, a formalin-fixed paraffin-embedded malignant colorectal cancer tissue was used to measure signal intensity and signal localization of each downstream modality (**Supplemental Figure 1**). First, PSR staining, and H&E staining were tested with or without prior N-glycan IMS (**Figure 1**). PSR can be imaged under both brightfield and polarized light. Under brightfield light, PSR stains the extracellular matrix red. Under polarized light, PSR staining enhances the natural birefringence of collagen fibers, producing red, yellow and green light depending on the birefringent-induced phase delay. Thicker, more organized, tightly packed fibers cause larger phase delays (red), whereas thinner, less organized, less tightly packed fibers cause smaller phase delays (green)^27^.

**Figure 1.**
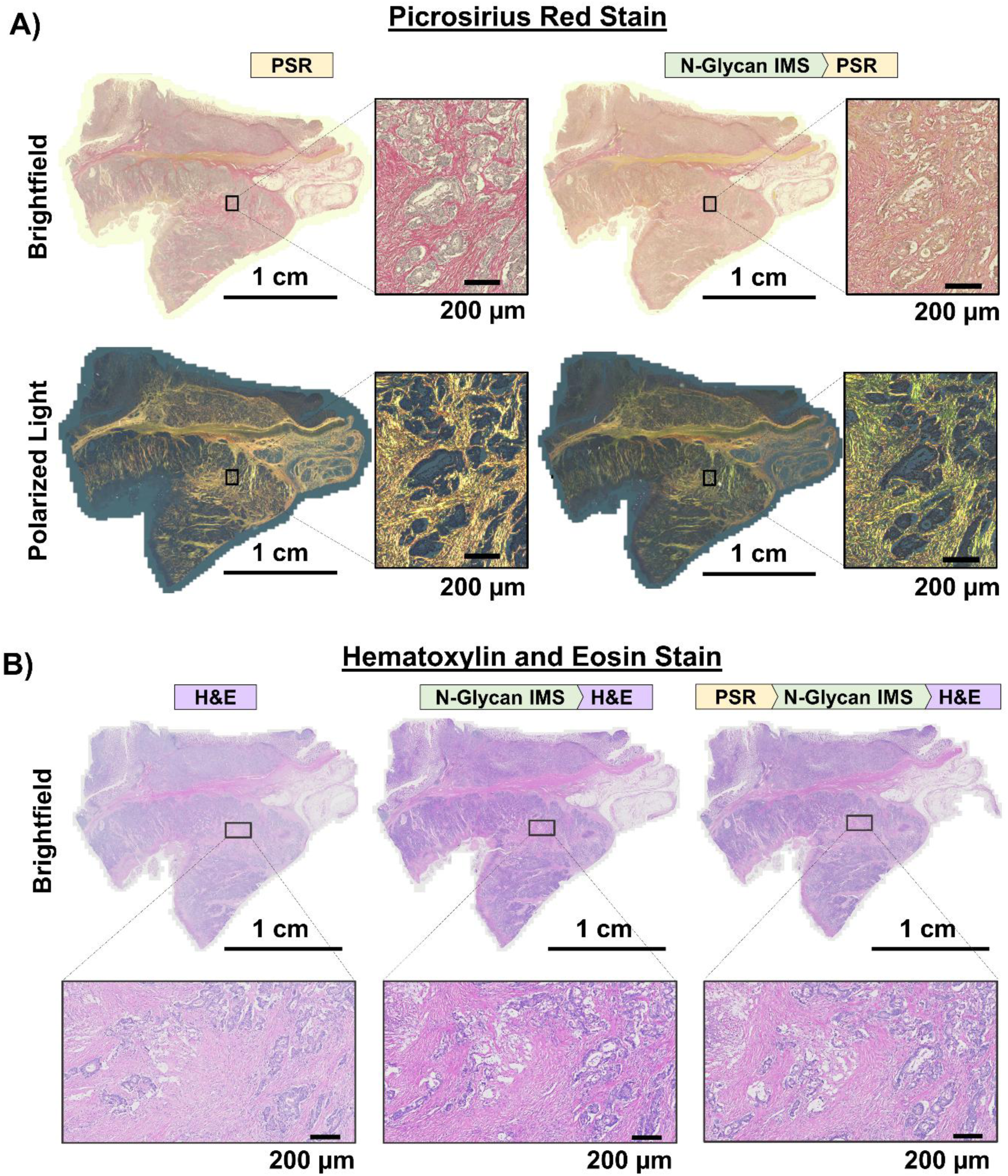
Integration of picrosirius red (PSR) and hematoxylin and eosin (H&E) staining at different stages of the multimodal imaging workflow. A) PSR and Weigert’s hematoxylin staining performed without (left) or with (right) prior N-glycan imaging mass spectrometry (IMS) acquired with brightfield light (top) and polarized light (bottom). B) H&E staining performed without prior modalities (left), performed after N-glycan IMS (middle) and performed after sequential PSR staining and N-glycan IMS (right). Images were acquired using a 20× objective, and identical display settings (0–255 intensity range) were applied to all images.

**Table 1.** Modalities used in this study to image formalin-fixed paraffin-embedded tissue microenvironments and corresponding features detected and spatial resolution acquired.

| **Modality** | **Features Detected** | **Pixel Size** |
| --- | --- | --- |
| **Autofluorescence** | Untargeted tissue morphology<br>Use: pixel-level co-registration of all modalities | 0.651 $\mu\text{m}$<br>10x objective<br>Zeiss AxioScan.Z1 |
| **Hematoxylin and eosin stain** | Individual cellular architecture (nuclei and cytoplasm)<br>Tissue morphology (extracellular matrix and connective tissue)<br>Use: pathological annotation | 0.220 $\mu\text{m}$<br>20x objective<br>Zeiss AxioScan.Z1 |
| **Picrosirius red stain** | Untargeted histology (brightfield)<br>Collagen fiber width and orientation (polarized light)<br>Use: quantification of collagen fibers | 0.220 $\mu\text{m}$<br>20x objective<br>Zeiss AxioScan.Z1 |
| **Multiplexed immunofluorescence** | Targeted protein or cell markers<br>Use: Identification of cell populations or protein composition | 0.508 $\mu\text{m}$<br>20x objective<br>Phenocycler Fusion 2.0 |
| **N-glycan imaging mass spectrometry** | Untargeted N-linked glycans<br>Use: direct detection and relative quantification of N-glycans | 80 $\mu\text{m}$ - PSR comparisons<br>40 $\mu\text{m}$ - MxIF comparisons<br>20 $\mu\text{m}$ – Multimodal Figure<br>timsTOF FleX |
| **Extracellular matrix imaging mass spectrometry** | Untargeted extracellular matrix peptides<br>Use: direct detection and quantification of collagen and extracellular matrix protein domains with post-translational modification status | 80 $\mu\text{m}$ - PSR comparisons<br>40 $\mu\text{m}$ - MxIF comparisons<br>20 $\mu\text{m}$ – Multimodal Figure<br>timsTOF Flex |

PSR staining of the extracellular matrix (red) under brightfield light after N-glycan IMS was muted when compared to PSR staining without prior N-glycan IMS. After N-glycan IMS, PSR-stained collagen fibers under polarized light produced smaller phase delays resulting in a shift toward green fibers compared to PSR stained collagen fibers without prior N-glycan IMS (**Figure 1A**). The shift toward green fibers after N-glycan IMS is likely a result of reduced collagen fiber organization and packing from N-glycan removal rather than reduction of fiber width^27^. N-glycans play an important role in protein folding and collagen fiber formation and stability. Thus, removal of N-glycans likely disrupts collagen fiber organization, decreasing the natural birefringence of collagen fibers, resulting in a shift from red to green fibers under polarized light.

After N-glycan IMS, the nuclei (purple, hematoxylin), cell cytoplasm (pink, eosin) and the extracellular matrix (pink, eosin) of H&E stains were stained darker without changes in tissue morphology. Eosin binds to basic residues in protein in the cytoplasm and extracellular matrix, whereas hematoxylin binds to DNA in the nucleus^12^. Staining was performed on all tissues at the same time with the same solutions to ensure any variation was due to alterations in tissue rather than batch effects. Therefore, the improved staining after N-glycan removal is likely due to increased accessibility to both protein backbones and nucleic acids from reduced bulky carbohydrate chain steric hindrance on nuclear, cellular and extracellular proteins. After sequential PSR staining and N-glycan IMS, there does not appear to be any carry over from the PSR stain to the H&E stain. The stain is darker than without any prior modalities, but slightly lighter than just prior N-glycan IMS alone (**Figure 1B**). The improved H&E staining after N-glycan removal supports previous workflows that integrate N-glycan IMS prior to H&E staining^22^.

To determine if imaging mass spectrometry (IMS) is impacted by prior picrosirius red (PSR) staining, signal intensity and signal localization for both N-glycan IMS and extracellular matrix IMS without (PSR-) or with (PSR+) prior PSR staining was assessed (**Figure 2**). Comparison of overlaid images of six representative N-glycans and quantification of the N-glycans (n = 160) across PSR-and PSR+ showed no apparent increase or decrease in signal (**Figure 2A, Supplemental Figure 2**). To evaluate changes in spatial localization, a segmentation by K-means (n = 5) with Manhattan distance was performed on each tissue section. Even with natural variation between serial sections (18 μm apart), the result showed very similar segments across PSR- and PSR+ experiments. Segments in both PSR- and PSR+ samples localized to non-stromal tumor regions and adjacent submucosa (yellow), normal-adjacent epithelial cells (purple), tumor center (blue), stromal tissue (red) and a sub-segment of stromal tissue at the tumor border (green) (**Figure 2B, Supplemental Figure 1**). To evaluate N-glycan composition of each segment, discriminating features for each segment were ranked by area under the receiver operating characteristic (ROC) curve (**Supplemental Table 3**). **Figure 2C** details the number of top 30 discriminating features shared across each segment. Each of the five PSR- segments shared at least 27 out of 30 top discriminating features by ROC curve with their corresponding PSR+ segment. Similar localization of red and green segments to stromal tissue regions is also reflected in the shared discriminating features (n = 24-25 shared top 30 discriminating features), suggesting that N-glycosylation of stromal tissue in the tumor border is similar to N-glycosylation in stromal tissue found throughout the tissue.

**Figure 2.**
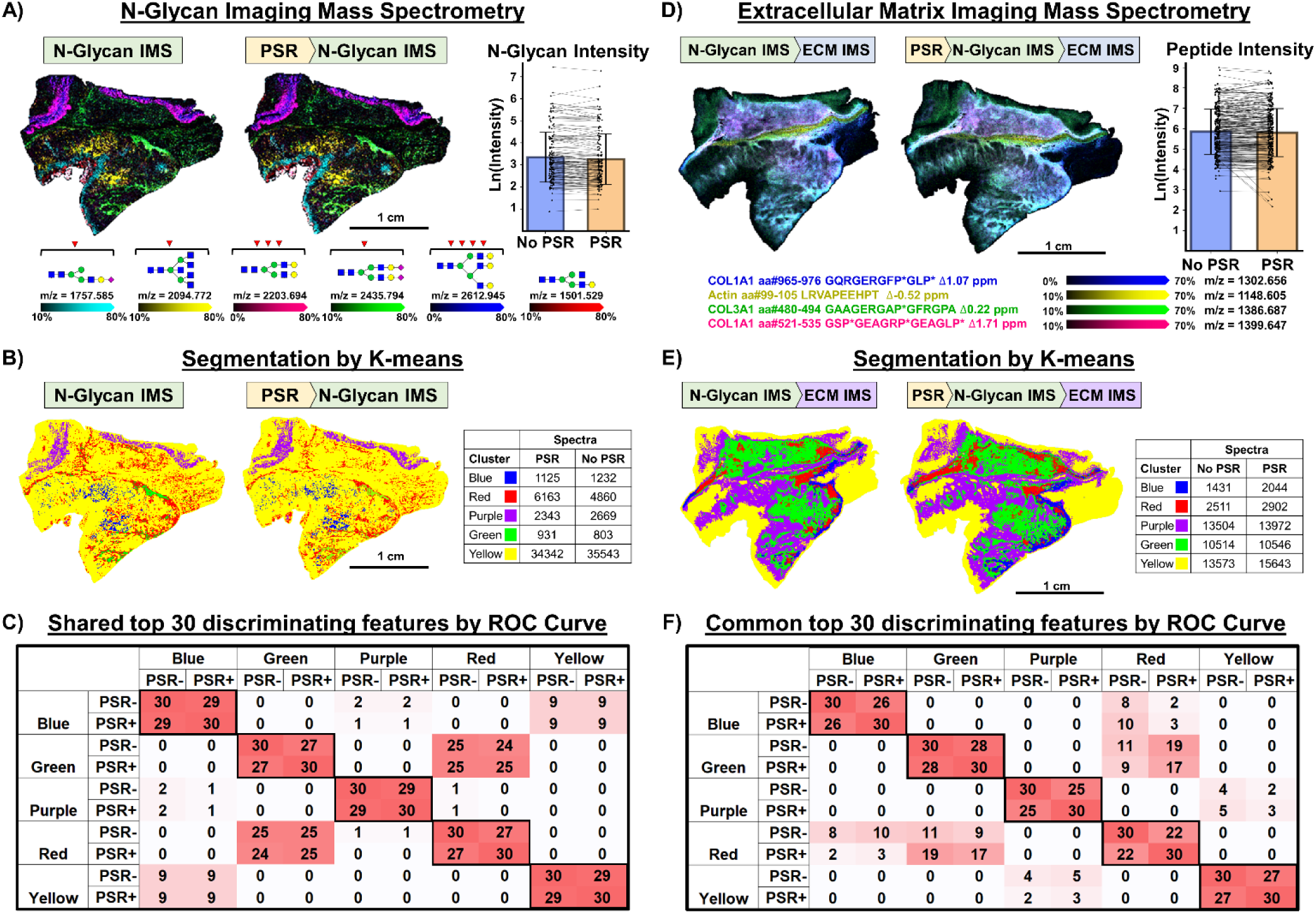
Imaging mass spectrometry (IMS) without or with prior picrosirius red stain (PSR). **A)** Overlaid heatmap of representative N-glycans from N-glycan IMS performed without (left) or with (right) prior PSR stain and corresponding paired quantification of natural log transformed maximum peak intensity of top 160 N-glycans by intensity. **B)** Segmentation by K-means with Manhattan distance using m/z values from N-glycan database (n = 239). **C)** Shared top 30 discriminate N-glycans by receiver operating characteristic (ROC) curve across each segment. **D)** Overlaid heatmap of representative peptides from extracellular matrix IMS performed following N-glycan IMS without (left) or with (right) prior PSR stain and corresponding paired quantification of natural log transformed maximum peak intensity of collagenase-generated peptides (n = 288). **E)** Segmentation by K-means with Manhattan distance using all collagenase-generated peptides (n = 288). **F)** Shared top 10 discriminate peptides by receiver operating characteristic (ROC) curve across each segment. N-glycan IMS sections are three 6 μm sections apart. ECM IMS sections are eighteen 6 μm sections apart. P* indicates hydroxyproline.

Similar to N-glycan IMS intensity, extracellular matrix peptide IMS peptide intensity was not noticeably different across PSR- and PSR+ experiments even with natural variation of serial sections (108 μm apart). Overlaid representative extracellular matrix peptide IMS images of Collagen Type I peptides (blue and pink), Collagen Type III peptide (green), and actin peptide (yellow) show similar peptide intensities and localization across the tissue (**Figure 2D, Supplemental Figure 2**). Quantification of 288 extracellular matrix peptides in PSR- and PSR+ samples does not show a trend toward increasing or decreasing extracellular matrix peptide intensity after PSR staining (**Figure 2D**). Segmentation by K-means (n = 5) with Manhattan distance resulted in segments that localized to tissue border and adjacent submucosa (yellow), tumor center (green), tumor edge (purple), and tumor border (red and blue) in both PSR- and PSR+ samples (**Figure 2E, Supplemental Figure 1**). The shared top 30 discriminating features by ROC curve across each corresponding segment (red = 22 shared features, purple = 25, blue = 26, yellow = 27, green = 28) further suggests that peptide composition was not drastically different across PSR- and PSR+ segments (**Figure 2F, Supplemental Table 4**). These results show that prior PSR staining does not impact the steps in N-glycan and extracellular matrix IMS, including but not limited to feature localization, enzymatic digestion, analyte co-crystallization with CHCA matrix and analyte ionization.

Integrating multiplexed immunofluorescence (MxIF) provides important high spatial resolution information on immune cell populations, tissue-specific cell types as well as other protein markers of interest^18^. This protocol involves simultaneous staining and subsequent fixing of up to 60 antibodies followed by hybridization and removal of fluorescently labeled DNA reporters via cyclic microfluidic administration of reporters (n = 3 per cycle) and washing with DMSO. Tissue subjected to prior multiplexed immunofluorescence (MxIF+) experienced 12 cycles of reporter administration and removal (35 antibodies total, **Supplemental Table 1**). The same experimental design as above was used to determine how N-glycan IMS and extracellular matrix IMS feature intensity and localization is affected by prior MxIF (**Figure 3**). Overlaid representative N-glycan mass spectrometry images and bar graph quantifying all 160 N-glycans shows an increase in intensity of all N-glycans after MxIF (MxIF+) when compared to N-glycan IMS without prior MxIF (MxIF-) (**Figure 3A, Supplemental Figure 3**). Segmentation by K-means (n = 5) with Manhattan distance resulted in segments that localized to tumor and submucosa (yellow), tumor center (blue), stroma (red), and normal-adjacent mucosa (purple) in both MxIF- and MxIF+ experiments (**Figure 3B, Supplemental Figure 1**). The last green segment localized to a subsection of normal-adjacent mucosa in the MxIF- tissue and a subsection of tumor-associated stroma in the MxIF+ tissue. The four segments with similar localization shared most of their top 30 discriminating features (blue = 22 shared top discriminate features, purple = 26, red = 22, yellow = 23) (**Figure 3C, Supplemental Table 3**). The green segment did not share any top discriminate features across MxIF- and MxIF+ experiments. Instead, the green MxIF- segment co-localized to a region within the purple MxIF- segment (22 shared top discriminate features) innormal-adjacent mucosa, whereas the green MxIF+ segment co-localized to a region within the red MxIF+ segment (21 MxIF- and 27 MxIF+ shared discriminate features) in the tumor region (**Supplemental Figure 4**). Differential green sub-segments sharing top discriminating features of pre-existing shared segments suggest that there are no uniquely distinct compositional N-glycan signatures that are emerging or disappearing after MxIF despite different segmentation.

**Figure 3.**
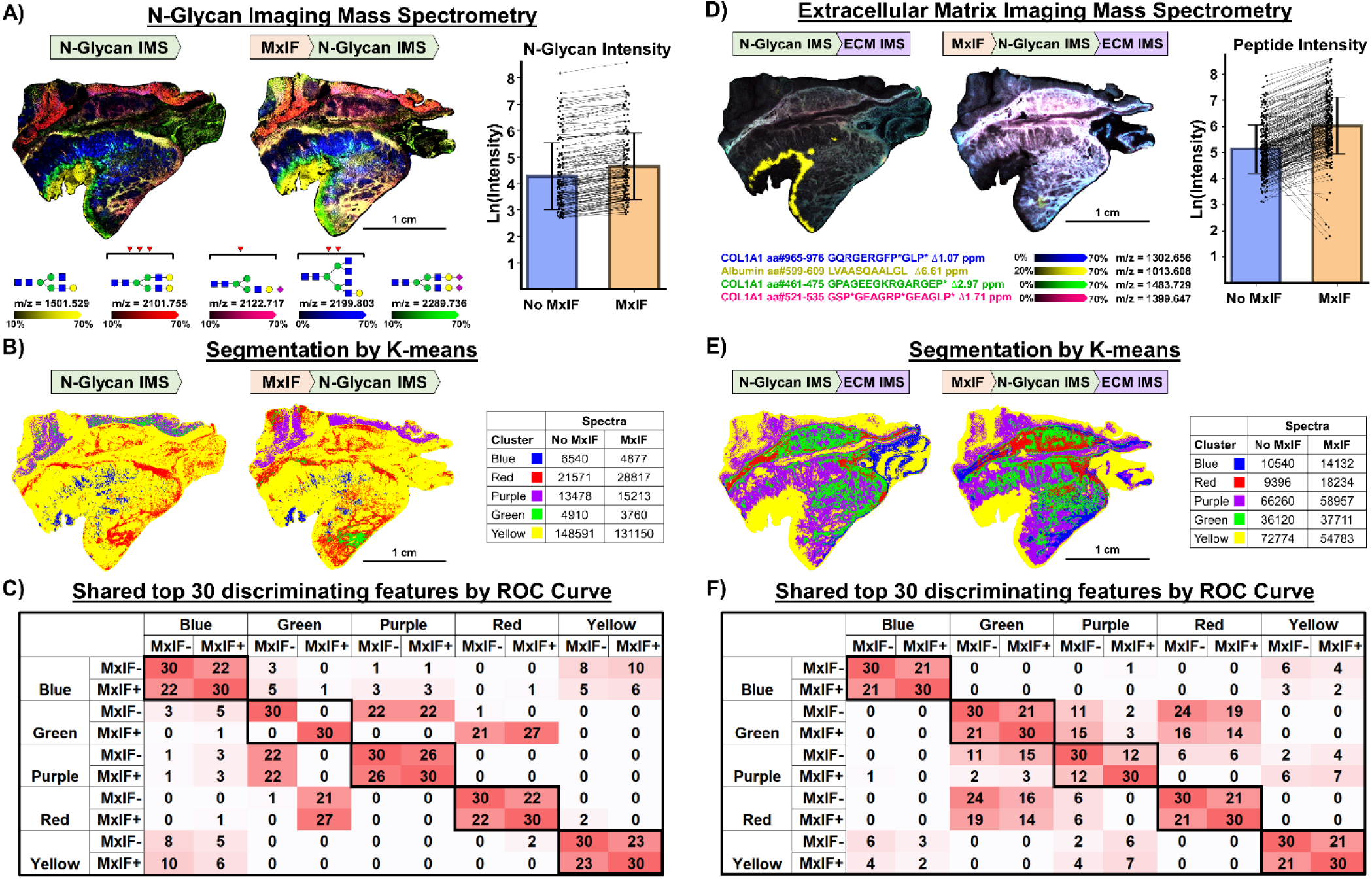
Imaging mass spectrometry (IMS) without or with prior multiplexed immunofluorescence (MxIF). **A)** Overlaid heatmap of representative N-glycans from N-glycan IMS performed without (left) or with (right) prior PSR stain and corresponding paired quantification of natural log transformed maximum peak intensity of top 160 N-glycans by intensity. **B)** Segmentation by K-means with Manhattan distance using m/z values from N-glycan database (n = 239). **C)** Shared top 30 discriminate N-glycans by receiver operating characteristic (ROC) curve across each segment from K-means segmentation. **D)** Overlaid heatmap of representative peptides from extracellular matrix IMS performed after N-glycan IMS without (left) or with (right) prior PSR stain and corresponding paired quantification of natural log transformed maximum peak intensity of collagenase-generated peptides (n = 288). **E)** Segmentation by K-means with Manhattan distance using all collagenase-generated peptides (n = 288). **F)** Shared top 10 discriminate peptides by receiver operating characteristic (ROC) curve across each segment. Sections are two 6 μm sections apart. P* indicates hydroxyproline.

Similar to N-glycan IMS, extracellular matrix IMS peptide intensity overall increased after MxIF. Representative extracellular matrix IMS images of peptides from Collagen Type I (green, blue and pink) show an increase in intensity after MxIF. A representative IMS image of an albumin peptide shows a dramatic loss in signal (yellow) in the center of the bottom tumor (**Figure 3A, Supplemental Figure 3**). Quantification of all 288 extracellular matrix peptides show a trend of increased maximum peak intensity after MxIF with only nine peptides decreasing in maximum peak intensity after MxIF. The five peptides (m/z = 789.331, 1013.606, 1622.946, 1847.066, and 1864.091) with the highest percent change (67%, 96%, 94%, 94%, and 95%, respectively) co- localized (0.37, 1.00, 0.84, 0.87, and 0.86, respectively by Pearson’s Correlation Coefficient) to the representative albumin peak (yellow), suggesting loss of signal is specific to this histological region rather than the workflow itself (**Supplemental Table 5**). Despite increases in ECM peptide intensity, segmentation by K-means with Manhattan distance resulted in segments with similar localization to tissue border (yellow), tumor border and submucosa (blue), non-stromal tumor and normal-adjacent mucosa (purple), and tumor stroma (green and red) in both MxIF- and MxIF+ (**Figure 3E, Supplemental Figure 1**). Furthermore, blue, green, red, and yellow segments had similar peptide intensity profiles, sharing 21 top 30 discriminating features across MxIF- and MxIF+ tissue sections. The purple segment, which localized to histological regions that are not rich in extracellular matrix proteins, only shared 12 shared top discriminating features (**Figure 3F, Supplemental Table 4**). Since K-means segments are defined by extracellular matrix peptide intensity profiles, top discriminating features of segments that localize to histological features that are not rich in extracellular matrix proteins (yellow and purple) are most likely to be affected by increases in peptide intensity as these discriminating features are defined by a lack of extracellular matrix peptide intensity. Overall, IMS signal intensity increased without significant impact on feature localization. MxIF involves additional antigen retrieval and fixation steps as well as repetitive washings with high DMSO solvent to remove DNA probes after each cycle^18^. The overall increase in both N-glycan and collagen peptide signal is likely due to decreased ion suppression from extensive washing and removal of small molecules during wash cycles of MxIF that are not removed during traditional dewaxing and antigen retrieval steps.

In conclusion, N-glycan IMS prior to chemical staining did not impact overall tissue morphology but altered picrosirius red staining and improved hematoxylin and eosin staining. Furthermore, PSR staining prior to N-glycan IMS did not affect N-glycan and extracellular matrix peptide signal intensity nor feature localization. Based on these results, a sequential workflow of PSR staining, N-glycan IMS, H&E and extracellular matrix IMS can be performed on the same tissue section for comprehensive imaging of the extracellular matrix. To include cell population information, multiplexed immunofluorescence can be performed prior to PSR staining resulting in increases in intensity of all measured N-glycans (n = 160) and an increase in intensity of 96.9% of all measured extracellular matrix peptides (n = 279/288) without changes in feature localization. The proposed workflow integrating all modalities on the same tissue section is as follows: autofluorescence microscopy (initial co-registration), multiplexed immunofluorescence (high-resolution cell population and protein localization), picrosirius red stain (collagen fiber width and orientation),

N-glycan IMS (N-glycans), autofluorescence microscopy (with matrix-containing laser ablation marks for pixel-level co-registration), H&E stain (single-cell spatial resolution tissue morphology), extracellular matrix IMS (post-translationally modified extracellular matrix peptides) and a final autofluorescence microscopy (with matrix-containing laser ablation marks for pixel-level co-registration) (**Figure 4**). Overall, this integrated multimodal workflow produces data on cell populations, collagen fibers and tissue histology at single-cell spatial resolution as well as multiplexed N-glycan and post-translationally modified extracellular matrix protein domain data for a spatially preserved 2-dimensional landscape of the cellular and extracellular tissue microenvironment. Performing these modalities on the same tissue section allows for retention of spatial relationships, improving co-registration of across modalities and providing a framework to investigate spatial gradients of cell-extracellular matrix crosstalk in formalin-fixed paraffin-embedded tissues.

**Figure 4:**
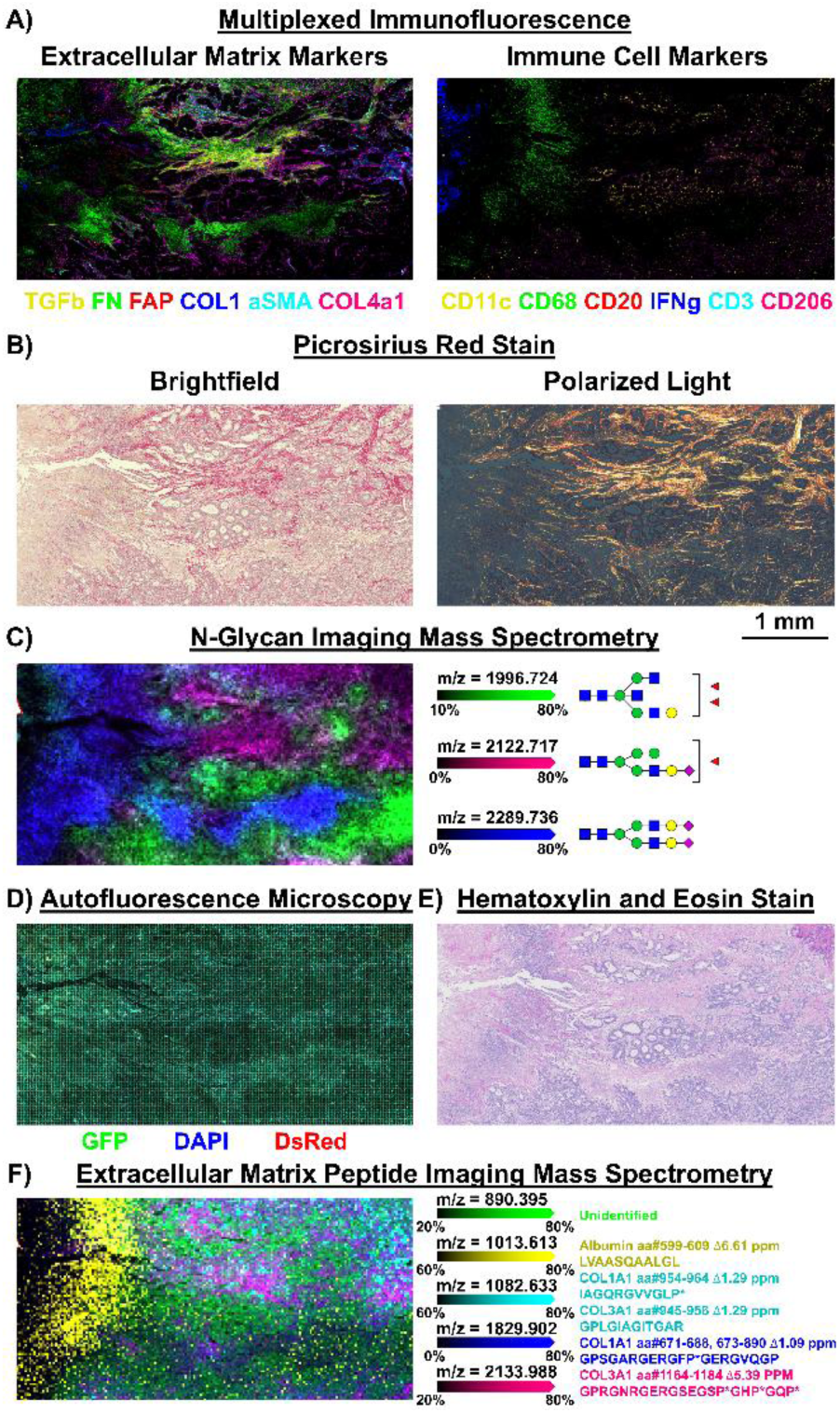
Optimized same-tissue multimodal workflow. Images from **A)** multiplexed immunofluorescence with representative extracellular matrix markers (left) and immune cell markers (right); **B)** picrosirius red staining under brightfield light (left) and polarized light (right); **C)** N-glycan imaging mass spectrometry; **D)** autofluorescence microscopy showing laser ablation of α-cyano-4-hydroxycinnamic acid following N-glycan imaging mass spectrometry; **E)** hematoxylin and eosin stain and **F)** extracellular matrix imaging mass spectrometry. All modalities were performed on the same tissue section and images shown are from the same region of interest.

## ASSOCIATED CONTENT

### Supporting Information

The following files are available free of charge.

Figure S1: Pathologically annotated hematoxylin and eosin stain of study tissue and study overview. (PDF)

Figure S2: Individual IMS ion images of representative features from picrosirius red stain comparison. (PDF)

Figure S3: Individual IMS ion images of representative features from multiplexed immunofluorescence comparison. (PDF)

Figure S4: Green N-glycan segments and corresponding individual ion images of top 10 discriminating features from multiplexed immunofluorescence comparison. (PDF)

Table S1: Multiplexed immunofluorescence antibody panel. (PDF)

Table S2: Matched LC-TIMS-MS/MS peptide library. (PDF)

Table S3: All discriminating N-glycan features of each K-means segment by ROC curve. (PDF)

Table S4: All discriminating ECM peptide features of each K-means segment by ROC curve. (PDF)

Table S5: Peptides with decreasing maximum peptide intensity after MxIF. (PDF)

Mass spectrometry data is available as a MassIVE Dataset with ID #MSV000102886. Instructions on accessing the data can be found <u>here</u>.

## AUTHOR INFORMATION

### Author Contributions

The manuscript was written through contributions of all authors. All authors have given approval to the final version of the manuscript.

## ABBREVIATIONS

AF: autofluorescence
CHCA: α-cyano-4-hydroxycinnamic acid
DAPI: 4’,6-diamidino-2-phenylindole
DMSO: dimethyl sulfoxide
DNA: deoxyribonucleic acid
DsRed: *Discosoma* red; ECM, extracellular matrix
GFP: green fluorescent protein
H&E: hematoxylin and eosin
IMS: imaging mass spectrometry
LC-MS: liquid chromatography mass spectrometry
LC-MS/MS: liquid chromatography tandem mass spectrometry
ITO: indium tin oxide
MALDI: matrix-assisted laser/desorption ionization
MxIF: multiplexed immunofluorescence
N-glycan: asparagine-linked glycan
PNGase F: Peptide:N-glycosidase F
Ppm: parts per million
PSM: peptide spectrum match
PSR: picrosirius red
ROC: receiver operating characteristic
SHG: second harmonic generation
TIMS: trapped ion mass spectrometry.

---

TOC Figure: This study develops a same-tissue multimodal spatial workflow by systematically assessing integration of picrosirius red staining, hematoxylin and eosin staining, multiplexed immunofluorescence, N-glycan imaging mass spectrometry and extracellular matrix imaging mass spectrometry.

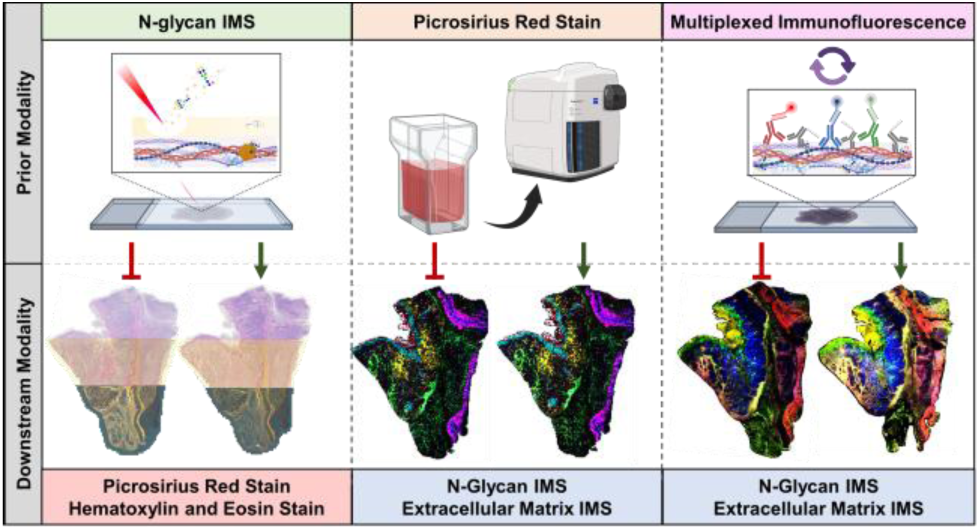

