## Supplemental Figure 1 for "Multimodal Imaging of the Cellular and Extracellular Microenvironment on the Same Formalin-Fixed Paraffin-Embedded Tissue Section"

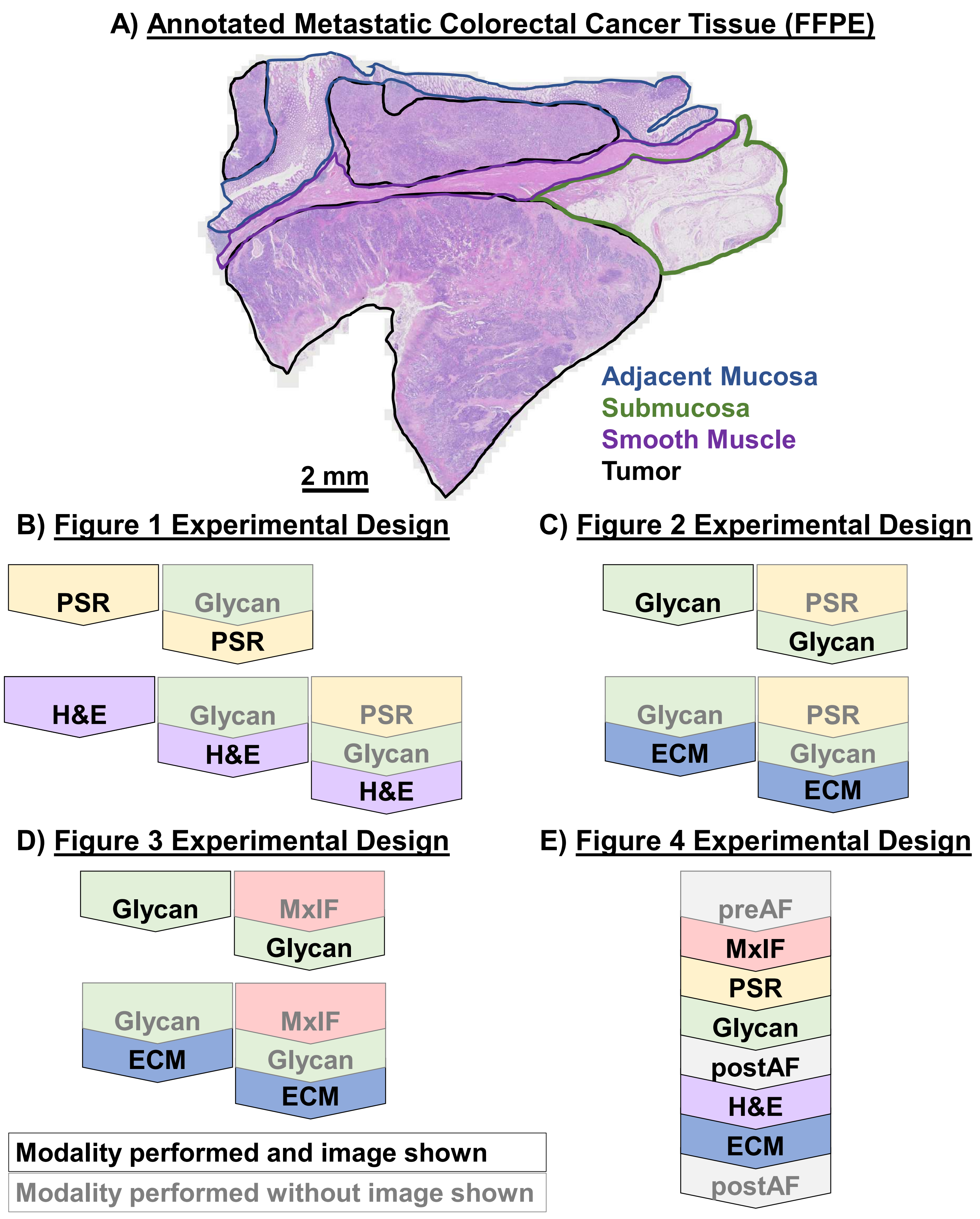


**Supplemental Figure 1: Study overview. A)** Pathologically-annotated hematoxylin and eosin stain of the formalin-fixed paraffin-embedded malignant colorectal cancer tissue used in this study. Experimental design overviews of **B)** figure 1, **C)** figure 2, **D)** figure 3, and **E)** figure 4. Modalities that are presented in each figure are indicated with black text and a black outline. Relevant modalities that were performed but not shown in figures are indicates with grey text and a grey outline. Autofluorescent microscopy (AF), multiplexed immunofluorescence (MxIF), picrosirius red stain (PSR), N-glycan imaging mass spectrometry (Glycan), hematoxylin and eosin stain (H&E), extracellular matrix imaging mass spectrometry (ECM).
