## Supplemental Figure 2 for "Multimodal Imaging of the Cellular and Extracellular Microenvironment on the Same Formalin-Fixed Paraffin-Embedded Tissue Section"

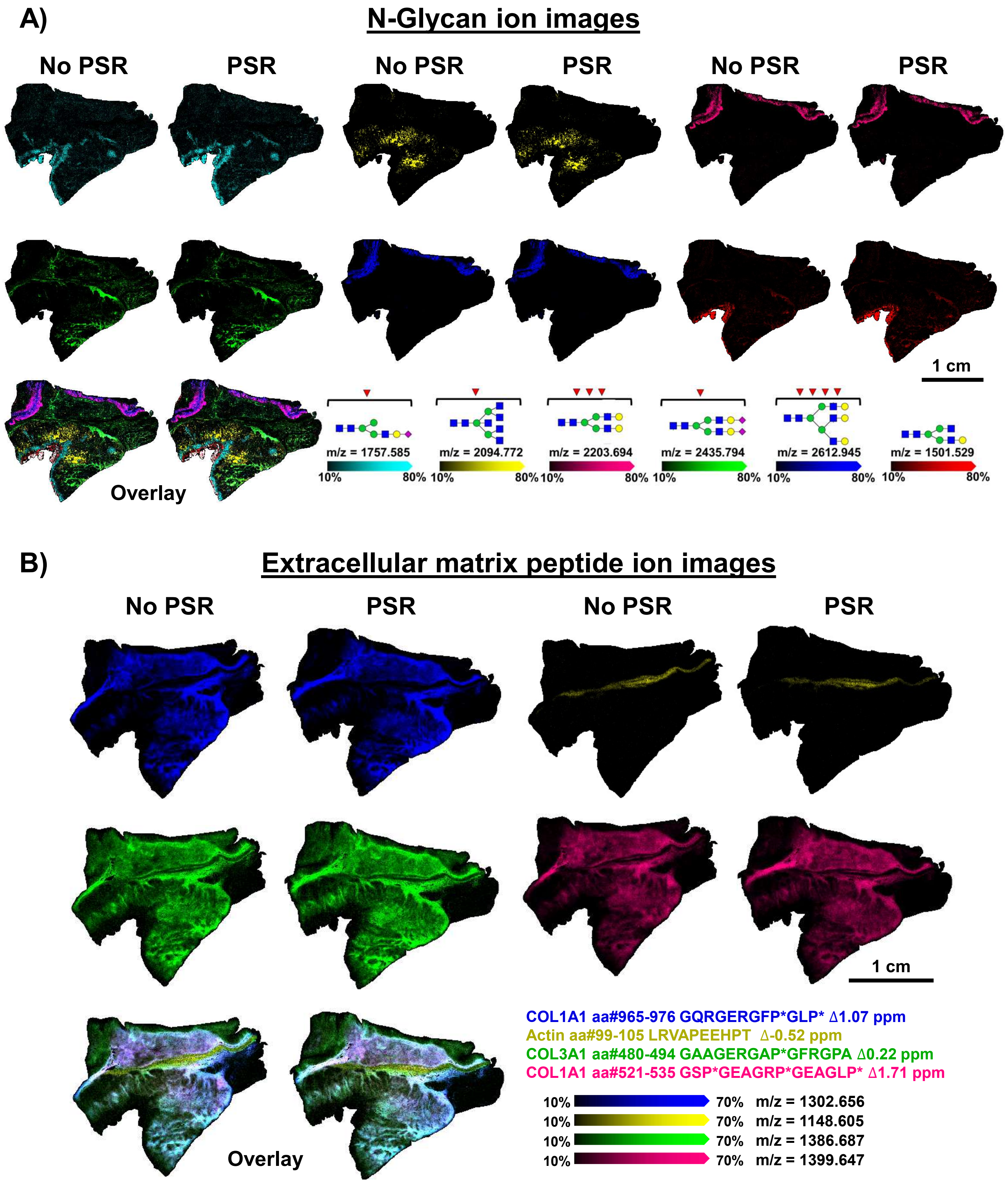


Supplemental Figure 2: Overlay and individual ion images of A) N-glycans and B) extracellular matrix peptides from picrosirius red staining experiments.
