## Supplemental Figure 3 for "Multimodal Imaging of the Cellular and Extracellular Microenvironment on the Same Formalin-Fixed Paraffin-Embedded Tissue Section"

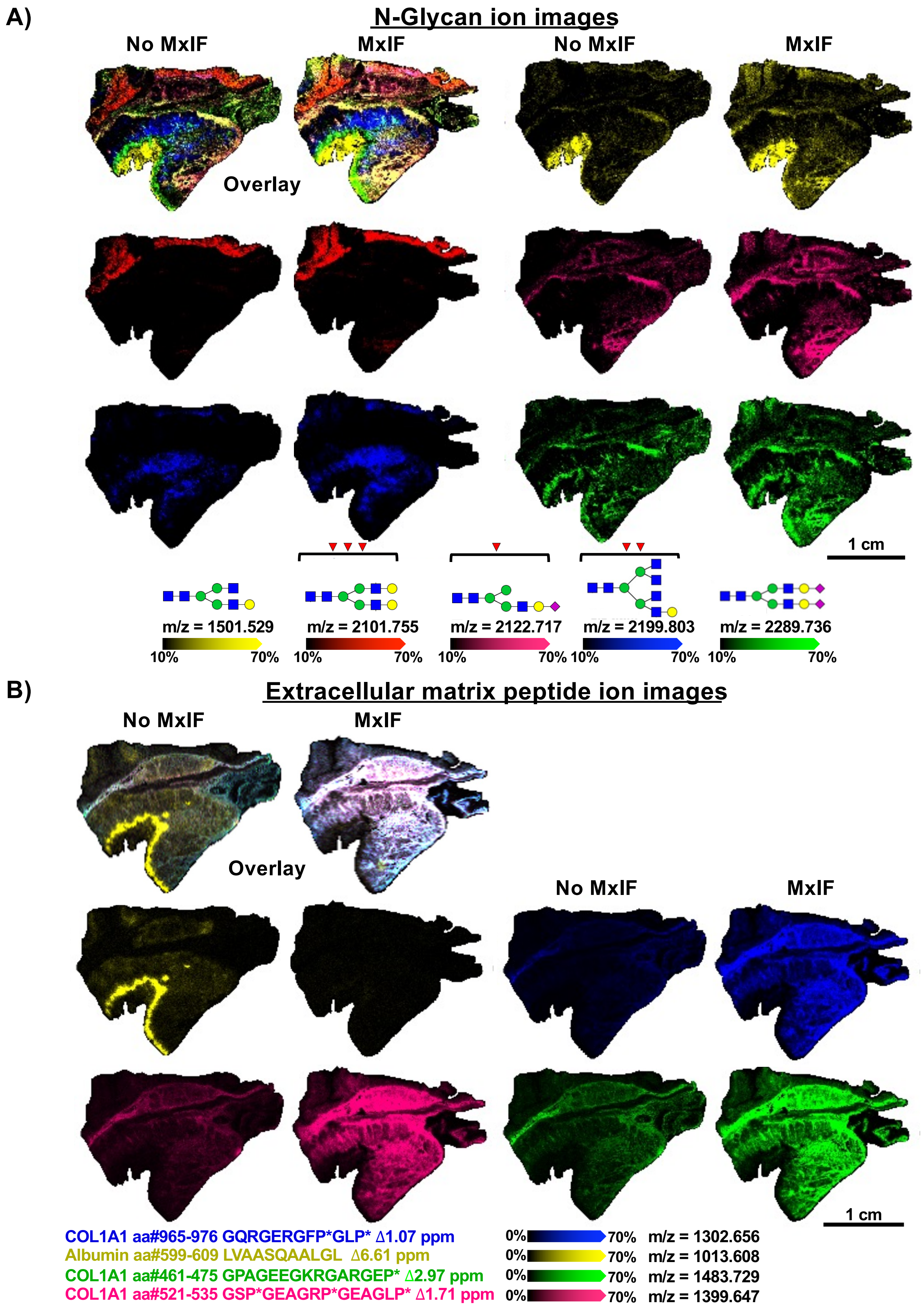


**Supplemental Figure 3: Overlay and individual ion images of A) N-glycans and B) extracellular matrix peptides from multiplexed immunofluorescence experiments.**
