## Supplemental Figure 4 for "Multimodal Imaging of the Cellular and Extracellular Microenvironment on the Same Formalin-Fixed Paraffin-Embedded Tissue Section"

### A) Green K-means Segment – MxIF-

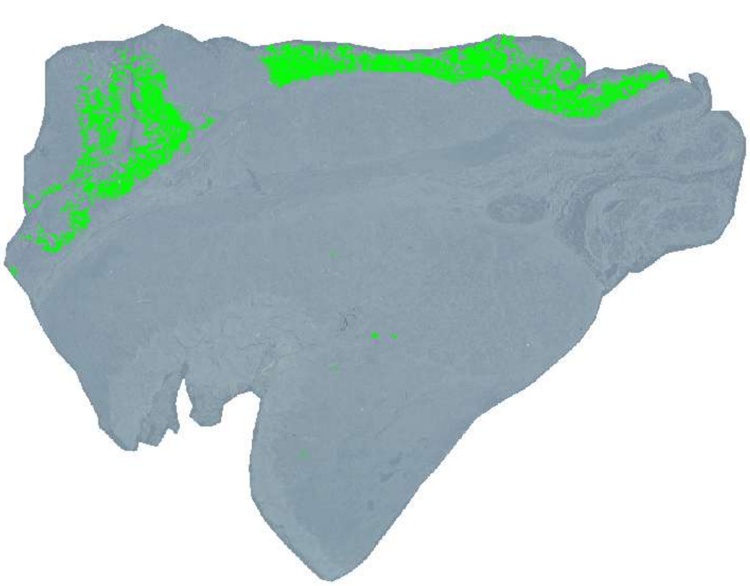

### C) Green K-means Segment – MxIF+

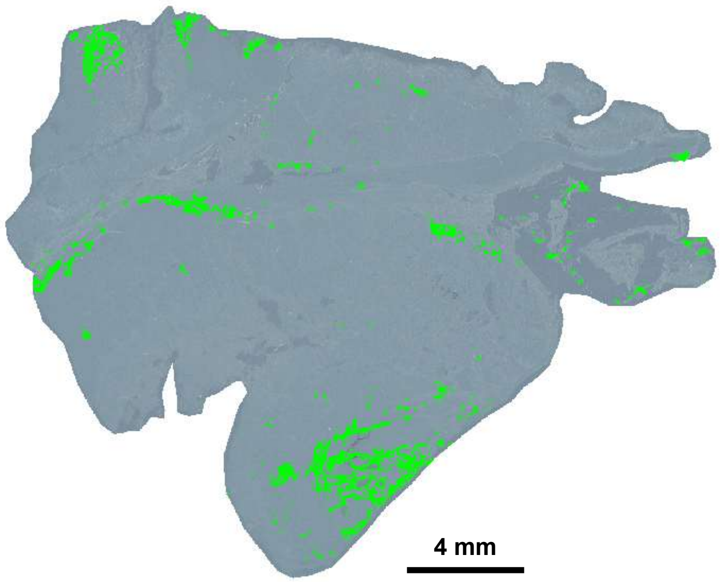


### B) Top 10 Discriminating Features of Green K-means Segment (MxIF-)

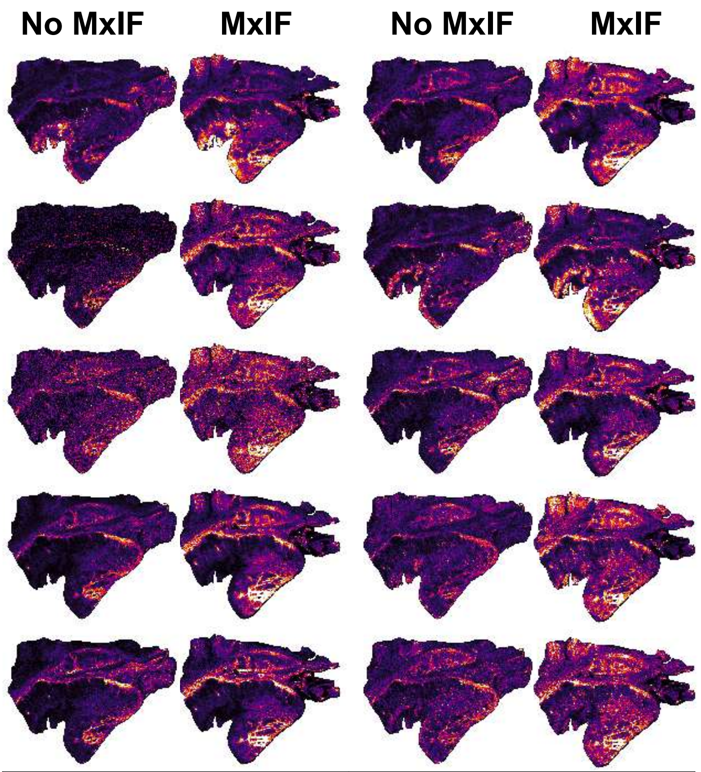


| | |
|---|---|
| m/z = 1428.512 | m/z = 1590.565 |
| m/z = 1631.592 | m/z = 1793.644 |
| m/z = 1850.666 | m/z = 1996.724 |
| m/z = 2101.755 | m/z = 2158.777 |
| m/z = 2304.835 | m/z = 2612.945 |

### D) Top 10 Discriminating Features of Green K-means Segment (MxIF+)

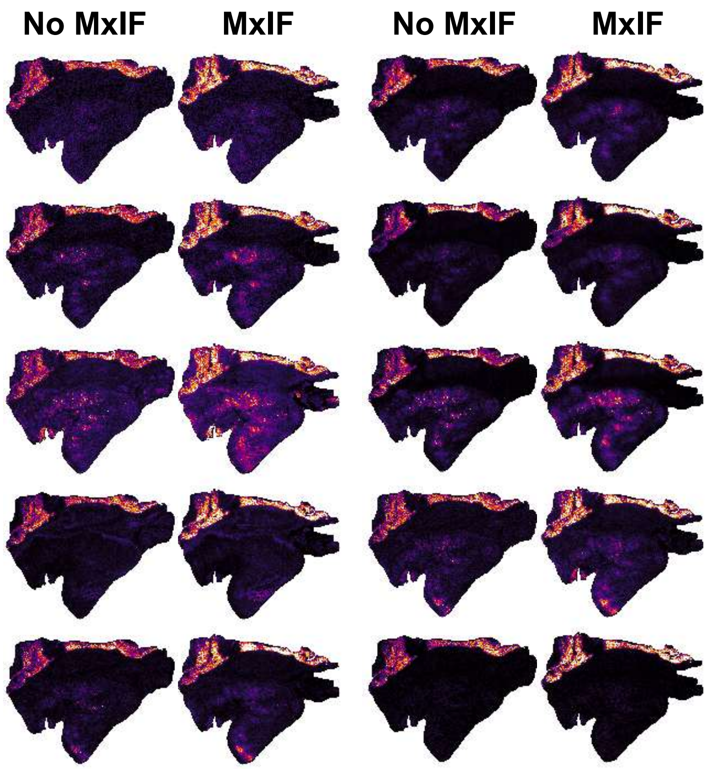


| | |
|---|---|
| m/z = 1663.581 | m/z = 1809.639 |
| m/z = 1837.541 | m/z = 1976.659 |
| m/z = 2048.599 | m/z = 2100.735 |
| m/z = 2122.717 | m/z = 2174.772 |
| m/z = 2435.794 | m/z = 2487.849 |

**Supplemental Figure 4: Green K-means segment from N-glycan imaging mass spectrometry image without (MxIF-) and with (MxIF+) prior multiplexed immunofluorescence. A)** Green K-means segment and **B)** corresponding top 10 discriminating features by receiver operating characteristic from MxIF- localize to normal adjacent tissue. **C)** Green K-means segment and **D)** corresponding top 10 discriminating features by receiver operating characteristic from MxIF+ localize to stromal tissue within tumor regions.
