## Supplemental Table 1 for "Multimodal Imaging of the Cellular and Extracellular Microenvironment on the Same Formalin-Fixed Paraffin-Embedded Tissue Section"

**Table 1: Multiplexed immunofluorescence antibody panel.** List of antibodies and corresponding channel, barcode and catalog number. Cycles indicate the order in which reporters were hybridized and imaged. Catolog numbers for each antibody are color coded based on supplier company.

| **Channel** | **ATTO550** | | | **CY5** | | | **AF750** | | |
|---|---|---|---|---|---|---|---|---|---|
| **Cycle** | **Antibody** | **Barcode** | **Cat No** | **Antibody** | **Barcode** | **Cat No** | **Antibody** | **Barcode** | **Cat No** |
| 1 | CD8 | BX020 | MA5-13473 | CD206 | BX120 | 321102 | IFNg | BX037 | ab231301 |
| 2 | TGFBI | BX017 | ab249517 | CD68 | BX036 | ab233172 | FoxP3 | BX022 | ab96048 |
| 3 | MMP9 | BX076 | ab204850 | EpCAM | BX091 | AKYP0119 | CD20 | BX028 | ab236434 |
| 4 | CD11b | BX050 | ab20997 | CD4 | BX021 | ab288725 | Ki67 | BX043 | ab279657 |
| 5 | MUC5AC | BX023 | 60100SF | Emilin1 | BX468 | ab243324 | CD3 | BX010 | ab245731 |
| 6 | COL4A3 | BX035 | 7076 | DCN | BX135 | ab175404 | PDGFRA | BX073 | ab271835 |
| 7 | MUC2 | BX070 | ab272706 | Fibronectin | BX033 | ab268022 | CD45 | BX001 | ab269347 |
| 8 | CD133 | BX500 | ab284397 | B-Catenin | BX003 | ab196204 | GLUT1 | BX019 | ab237900 |
| 9 | PD-L1 | BX078 | ab226766 | Collagen I | BX054 | ab215969 | FAP | BX013 | 81192 |
| 10 | CD27 | BX026 | ab256583 | Vimentin | BX042 | ab193555 | PD-1 | BX025 | 63815SF |
| 11 | aSMA | BX029 | ab240654 | VEGFR2 | BX030 | ab243912 | CD11c | BX016 | 14-9761-82 |
| 12 | COL4A1 | BX014 | ab226485 | | | | e-Cad | BX031 | F748 |

| **Supplier Company** |
|---|
| Abcam (Cambridge, UK) |
| ThermoFisher Scientific (Waltham, MA) |
| Cell Signaling Technology (Danvers, MA) |
| R&D Systems (Minneapolis, MN) |
| Quaterix (Billerica, MA) |
| Chondrex Inc, Woodinville, WA) |
| BioLegend (San Diego, CA) |
