## Supplemental Table 2 for "Multimodal Imaging of the Cellular and Extracellular Microenvironment on the Same Formalin-Fixed Paraffin-Embedded Tissue Section"

**Supplemental Table 2 (1/4): Matched peptide library.** Peptides from the liquid chromatography tandem mass spectrometry peptide library whose calculated protonated peptide mass is within 8 ppm of a calibrated matrix-assisted laser/desorption ionization mass to charge ratio. The "Modified Peptide" column includes peptide sequence with proline hydroxylation sites indicated with an asterisk. All detectable modification sites including proline hydroxylation, methionine oxidation, arginine deamidation and glutamine deamidation are listed under "Assigned Modifications" column. Peptide location within the protein is indicated with "Start" and "End" columns. Peptides with "FALSE" under the "Is Unique" column map to an additional protein and the most probable protein is listed under "Protein Description."

| Calculated Peptide Mass (+H) | Modified Peptide | Assigned Modifications | Peptide Length | Start | End | Protein ID | Gene | Protein Description | Is Unique |
|---|---|---|---|---|---|---|---|---|---|
| 758.4519 | ITGARGLA | | 8 | 952 | 959 | P02461 | COL3A1 | Collagen alpha-1(III) chain | TRUE |
| 758.4519 | IAGITGAR | | 8 | 949 | 956 | P02461 | COL3A1 | Collagen alpha-1(III) chain | TRUE |
| 781.4315 | GPVGARGPA | | 9 | 1079 | 1087 | P02452 | COL1A1 | Collagen alpha-1(I) chain | TRUE |
| 781.4315 | GPAGAVGPR | | 9 | 988 | 996 | P08123 | COL1A2 | Collagen alpha-2(I) chain | TRUE |
| 785.39 | GPAGSRGAP* | 9P(15.9949) | 9 | 1077 | 1085 | P02461 | COL3A1 | Collagen alpha-1(III) chain | TRUE |
| 789.3373 | GEQGPSGAS | | 9 | 1127 | 1135 | P02452 | COL1A1 | Collagen alpha-1(I) chain | TRUE |
| 797.4264 | GLRGGAGPP* | 9P(15.9949) | 9 | 693 | 701 | P02461 | COL3A1 | Collagen alpha-1(III) chain | TRUE |
| 797.4264 | GLRGGAGP*P | 8P(15.9949) | 9 | 693 | 701 | P02461 | COL3A1 | Collagen alpha-1(III) chain | TRUE |
| 804.4284 | LMNLGGLA | 2M(15.9949) | 8 | 129 | 136 | P37802 | TAGLN2 | Transgelin-2 | TRUE |
| 807.3731 | LTGDATTE | | 8 | 987 | 994 | Q99715 | COL12A1 | Collagen alpha-1(XII) chain | TRUE |
| 809.4628 | GPVGPVGAR | | 9 | 1076 | 1084 | P02452 | COL1A1 | Collagen alpha-1(I) chain | TRUE |
| 812.3937 | GYPGPAGPP | | 9 | 171 | 179 | P02461 | COL3A1 | Collagen alpha-1(III) chain | TRUE |
| 812.3971 | GPMGPPGLA | 3M(15.9949) | 9 | 998 | 1006 | P02452 | COL1A1 | Collagen alpha-1(I) chain | TRUE |
| 812.3971 | GPAGIMGPP* | 9P(15.9949) | 9 | 448 | 456 | P12107 | COL11A1 | Collagen alpha-1(XI) chain | TRUE |
| 822.3853 | GPQGHAGAQ | | 9 | 372 | 380 | P02461 | COL3A1 | Collagen alpha-1(III) chain | TRUE |
| 827.4005 | GPAGERGAP* | 9P(15.9949) | 9 | 504 | 512 | P02461 | COL3A1 | Collagen alpha-1(III) chain | TRUE |
| 827.4005 | GPRGAAGEP* | 9P(15.9949) | 9 | 516 | 524 | P02461 | COL3A1 | Collagen alpha-1(III) chain | TRUE |
| 827.4006 | PGGNLGAQN | | 9 | 73 | 81 | Q99497 | PARK7 | Parkinson disease protein 7 | TRUE |
| 827.437 | GPIGSRGPS | | 9 | 604 | 612 | P08123 | COL1A2 | Collagen alpha-2(I) chain | TRUE |
| 833.3934 | GADGRAGVM | | 9 | 409 | 417 | P08123 | COL1A2 | Collagen alpha-2(I) chain | TRUE |
| 833.3941 | FAPGQWAG | | 8 | 82 | 89 | P30622 | CLIP1 | CAP-Gly domain-containing linker protein 1 | TRUE |
| 835.368 | LAEASSEE | | 8 | 326 | 333 | Q14151 | SAFB2 | Scaffold attachment factor B2 | TRUE |
| 841.4162 | GPRGETGPA | | 9 | 908 | 916 | P02452 | COL1A1 | Collagen alpha-1(I) chain | TRUE |
| 843.3955 | GRDGVP*GGP* | 6P(15.9949),9P(15.9949) | 9 | 525 | 533 | P02461 | COL3A1 | Collagen alpha-1(III) chain | FALSE |
| 843.3955 | GPAGERGSP* | 9P(15.9949) | 9 | 506 | 514 | P02452 | COL1A1 | Collagen alpha-1(I) chain | TRUE |
| 843.4207 | VGVDGPNVS | | 9 | 162 | 170 | P55011 | SLC12A2 | Solute carrier family 12 member 2 | TRUE |
| 843.4207 | EANLGPGVS | | 9 | 561 | 569 | P07942 | LAMB1 | Laminin subunit beta-1 | TRUE |
| 843.4319 | GERGAAGLP* | 9P(15.9949) | 9 | 731 | 739 | P02452 | COL1A1 | Collagen alpha-1(I) chain | FALSE |
| 843.4319 | GRDGVAGVP* | 9P(15.9949) | 9 | 504 | 512 | P02462 | COL4A1 | Collagen alpha-1(IV) chain | TRUE |
| 849.3836 | ITDIENGS | 6N(0.9840) | 8 | 277 | 284 | Q9BXN1 | ASPN | Asporin | TRUE |
| 849.3883 | GADGRAGVM | 9M(15.9949) | 9 | 409 | 417 | P08123 | COL1A2 | Collagen alpha-2(I) chain | TRUE |
| 855.3955 | AGAAGP*AGNPG | 6P(15.9949) | 11 | 379 | 389 | P02452 | COL1A1 | Collagen alpha-1(I) chain | TRUE |
| 858.3991 | GPP*GFTGPP* | 3P(15.9949),9P(15.9949) | 9 | 197 | 205 | P02462 | COL4A1 | Collagen alpha-1(IV) chain | TRUE |
| 858.4025 | PAAGAPLMD | 8M(15.9949) | 9 | 72 | 80 | Q9NQC3 | RTN4 | Reticulon-4 | TRUE |
| 861.42 | LSPLDSET | | 8 | 1274 | 1281 | P11047 | LAMC1 | Laminin subunit gamma-1 | TRUE |
| 861.4213 | GFP*GARGPS | 3P(15.9949) | 9 | 410 | 418 | P02452 | COL1A1 | Collagen alpha-1(I) chain | TRUE |
| 861.4213 | GAP*GFP*GAR | 3P(15.9949),6P(15.9949) | 9 | 407 | 415 | P02452 | COL1A1 | Collagen alpha-1(I) chain | TRUE |
| 863.3893 | GPP*GVSGGGY | 3P(15.9949) | 10 | 1099 | 1108 | P08123 | COL1A2 | Collagen alpha-2(I) chain | TRUE |
| 863.3927 | VAMNPTNT | 3M(15.9949) | 8 | 59 | 66 | P11142 | HSPA8 | Heat shock cognate 71 kDa protein | FALSE |
| 865.3727 | DPFNPFE | | 7 | 395 | 401 | P53634 | CTSC | Dipeptidyl peptidase 1 | TRUE |
| 869.4475 | GPP*GLRGSP* | 3P(15.9949),9P(15.9949) | 9 | 394 | 402 | P08123 | COL1A2 | Collagen alpha-2(I) chain | TRUE |
| 869.4475 | GPIGSRGPE | | 9 | 249 | 257 | P05997 | COL5A2 | Collagen alpha-2(V) chain | TRUE |
| 872.4108 | KGSP*GADGPA | 4P(15.9949) | 10 | 934 | 943 | P02452 | COL1A1 | Collagen alpha-1(I) chain | TRUE |
| 872.4189 | FDFSFLP | | 7 | 1196 | 1202 | P02452 | COL1A1 | Collagen alpha-1(I) chain | TRUE |
| 872.4546 | LMP*GSVGPV | 3P(15.9949) | 9 | 210 | 218 | P05997 | COL5A2 | Collagen alpha-2(V) chain | TRUE |
| 872.4546 | INILGMDP | | 8 | 160 | 167 | P02748 | C9 | Complement component C9 | TRUE |
| 872.4546 | NILGMDPL | | 8 | 161 | 168 | P02748 | C9 | Complement component C9 | TRUE |
| 872.4584 | GQTGARGLP* | 9P(15.9949) | 9 | 211 | 219 | P08123 | COL1A2 | Collagen alpha-2(I) chain | TRUE |
| 875.3814 | EMPADLPS | 2M(15.9949) | 8 | 321 | 328 | P02768 | ALB | Albumin | TRUE |
| 875.3853 | GESGREGAP* | 9P(15.9949) | 9 | 1010 | 1018 | P02452 | COL1A1 | Collagen alpha-1(I) chain | TRUE |
| 875.3893 | P*GGP*GAAGFP* | 10P(15.9949),1P(15.9949),4P(15.9949) | 10 | 866 | 875 | P02461 | COL3A1 | Collagen alpha-1(III) chain | TRUE |
| 884.4148 | GP*LPP*GWE | 2P(15.9949),5P(15.9949) | 8 | 405 | 412 | O00308 | WWP2 | NEDD4-like E3 ubiquitin-protein ligase WWP2 | FALSE |
| 884.4182 | LYLQMNS | 5M(15.9949) | 7 | 98 | 104 | P01764 | IGHV3-23 | Immunoglobulin heavy variable 3-23 | FALSE |
| 884.422 | GPRGSEGPQ | | 9 | 359 | 367 | P02452 | COL1A1 | Collagen alpha-1(I) chain | TRUE |
| 884.422 | GPRGDTGPAG | | 10 | 1472 | 1481 | P25940 | COL5A3 | Collagen alpha-3(V) chain | TRUE |
| 884.4261 | GLHGFP*GAP* | 6P(15.9949),9P(15.9949) | 9 | 839 | 847 | P08572 | COL4A2 | Collagen alpha-2(IV) chain | TRUE |
| 884.4261 | GPQGFQGPP | | 9 | 197 | 205 | P02452 | COL1A1 | Collagen alpha-1(I) chain | TRUE |
| 887.3782 | LDSDGSFF | | 8 | 281 | 288 | P01857 | IGHG1 | Immunoglobulin heavy constant gamma 1 | FALSE |
| 887.3782 | FESGDFVS | | 8 | 242 | 249 | P22314 | UBA1 | Ubiquitin-like modifier-activating enzyme 1 | TRUE |
| 887.4469 | GLAGVGETPS | | 10 | 14 | 23 | P55011 | SLC12A2 | Solute carrier family 12 member 2 | TRUE |
| 887.4469 | GKDGLNGLP* | 6N(0.9840),9P(15.9949) | 9 | 1151 | 1159 | P02452 | COL1A1 | Collagen alpha-1(I) chain | TRUE |
| 891.4247 | FGFVTFSS | | 8 | 64 | 71 | P22626 | HNRNPA2B1 | Heterogeneous nuclear ribonucleoproteins A2/B1 | TRUE |
| 891.4319 | GARGFP*GTP* | 6P(15.9949),9P(15.9949) | 9 | 163 | 171 | P08123 | COL1A2 | Collagen alpha-2(I) chain | FALSE |
| 894.3952 | FSGPQNASS | | 9 | 519 | 527 | P20810 | CAST | Calpastatin | TRUE |
| 894.3992 | GPPSAGFDF | | 9 | 1190 | 1198 | P02452 | COL1A1 | Collagen alpha-1(I) chain | TRUE |
| 894.439 | FADIQMGI | | 8 | 343 | 350 | O75695 | RP2 | Protein XRP2 | TRUE |
| 897.4424 | GRP*GEVGP*P | 3P(15.9949),8P(15.9949) | 9 | 917 | 925 | P02452 | COL1A1 | Collagen alpha-1(I) chain | TRUE |
| 897.4425 | INNTAVGHA | 2N(0.9840) | 9 | 2557 | 2565 | P12111 | COL6A3 | Collagen alpha-3(VI) chain | TRUE |
| 897.4788 | GPRGEVGLP* | 9P(15.9949) | 9 | 280 | 288 | P08123 | COL1A2 | Collagen alpha-2(I) chain | TRUE |
| 897.4788 | GPDGLRGIP* | 9P(15.9949) | 9 | 1435 | 1443 | P20908 | COL5A1 | Collagen alpha-1(V) chain | FALSE |
| 900.4057 | P*GVNGAP*GEA | 1P(15.9949),7P(15.9949) | 10 | 915 | 924 | P08123 | COL1A2 | Collagen alpha-2(I) chain | TRUE |
| 900.417 | GEREGGAPQ | | 9 | 181 | 189 | Q5TF21 | MTCL3 | Microtubule cross-linking factor 3 | TRUE |
| 900.417 | GAPGERGEQ | | 9 | 2760 | 2768 | Q02388 | COL7A1 | Collagen alpha-1(VII) chain | TRUE |
| 900.417 | GERGEQGPA | | 9 | 629 | 637 | P02452 | COL1A1 | Collagen alpha-1(I) chain | TRUE |
| 900.417 | QGQQGGAGPT | | 10 | 227 | 236 | P05997 | COL5A2 | Collagen alpha-2(V) chain | TRUE |
| 908.4359 | IQIGNSFE | 5N(0.9840) | 8 | 163 | 170 | Q8WXQ8 | CPA5 | Carboxypeptidase A5 | TRUE |
| 908.4359 | LLQDGEFS | | 8 | 78 | 85 | P07737 | PFN1 | Profilin-1 | TRUE |
| 908.4359 | FEADISNL | | 8 | 149 | 156 | P08236 | GUSB | Beta-glucuronidase | TRUE |
| 908.4359 | IQLNGEFS | 4N(0.9840) | 8 | 328 | 335 | Q8TDX6 | CSGALNACT1 | Chondroitin sulfate N-acetylgalactosaminyltransferase 1 | TRUE |
| 908.436 | LQPASEYT | | 8 | 1056 | 1063 | P02751 | FN1 | Fibronectin | TRUE |
| 909.3737 | FQYDHEA | | 7 | 56 | 62 | Q15293 | RCN1 | Reticulocalbin-1 | FALSE |
| 912.4169 | AQGP*GGGGAGAP | 4P(15.9949) | 12 | 213 | 224 | P50552 | VASP | Vasodilator-stimulated phosphoprotein | TRUE |
| 912.417 | GQDGRP*GP*P | 6P(15.9949),8P(15.9949) | 9 | 560 | 568 | P02452 | COL1A1 | Collagen alpha-1(I) chain | TRUE |
| 912.417 | GQDGRP*GPP* | 6P(15.9949),9P(15.9949) | 9 | 560 | 568 | P02452 | COL1A1 | Collagen alpha-1(I) chain | TRUE |
| 912.4533 | GPQGVRGEP* | 9P(15.9949) | 9 | 365 | 373 | P02452 | COL1A1 | Collagen alpha-1(I) chain | TRUE |
| 912.4533 | P*GAAGRTGP*P | 1P(15.9949),9P(15.9949) | 10 | 789 | 798 | P08123 | COL1A2 | Collagen alpha-2(I) chain | TRUE |
| 912.4533 | DLNRQGP*P | 7P(15.9949) | 8 | 391 | 398 | Q9UBN4 | TRPC4 | Short transient receptor potential channel 4 | TRUE |
| 912.4533 | GERGVQGP*P | 8P(15.9949) | 9 | 683 | 691 | P02452 | COL1A1 | Collagen alpha-1(I) chain | TRUE |
| 912.4533 | GERGVQGPP* | 9P(15.9949) | 9 | 683 | 691 | P02452 | COL1A1 | Collagen alpha-1(I) chain | TRUE |
| 913.401 | GPRGEDGPE | | 9 | 835 | 843 | P20908 | COL5A1 | Collagen alpha-1(V) chain | FALSE |
| 914.4803 | LRGGGQAANA | | 10 | 1016 | 1025 | Q9Y6R7 | FCGBP | IgGFc-binding protein | TRUE |
| 919.4268 | FAGDDAPRA | | 9 | 23 | 31 | P62736 | ACTA2 | Actin, aortic smooth muscle | FALSE |
| 922.3974 | GQP*GVMGFP* | 2Q(0.9840),3P(15.9949),9P(15.9949) | 9 | 573 | 581 | P02461 | COL3A1 | Collagen alpha-1(III) chain | FALSE |
| 927.3764 | LDFEQEM | 7M(15.9949) | 7 | 221 | 227 | P63261 | ACTG1 | Actin, cytoplasmic 2 | FALSE |
| 927.3842 | P*GAGGGYYP*G | 1P(15.9949),9P(15.9949) | 10 | 7 | 16 | P30626 | SRI | Sorcin | TRUE |
| 938.4829 | VVESTGVFT | | 9 | 95 | 103 | P04406 | GAPDH | Glyceraldehyde-3-phosphate dehydrogenase | TRUE |
| 941.4435 | GPRGSEGPQG | | 10 | 359 | 368 | P02452 | COL1A1 | Collagen alpha-1(I) chain | TRUE |
| 943.4996 | ALNDHFVK | | 8 | 302 | 309 | P04406 | GAPDH | Glyceraldehyde-3-phosphate dehydrogenase | TRUE |
| 943.5095 | LQGVDLLAD | | 9 | 41 | 49 | P10809 | HSPD1 | 60 kDa heat shock protein, mitochondrial | TRUE |
| 943.5095 | IAGQVLDIN | 9N(0.9840) | 9 | 75 | 83 | P07910 | HNRNPC | Heterogeneous nuclear ribonucleoproteins C1/C2 | TRUE |
| 944.4908 | GSRGERGLP* | 9P(15.9949) | 9 | 877 | 885 | P08123 | COL1A2 | Collagen alpha-2(I) chain | TRUE |

**Supplemental Table 2 (2/4): Matched peptide library.** Peptides from the liquid chromatography tandem mass spectrometry peptide library whose calculated protonated peptide mass is within 8 ppm of a calibrated matrix-assisted laser/desorption ionization mass to charge ratio. The "Modified Peptide" column includes peptide sequence with proline hydroxylation sites indicated with an asterisk. All detectable modification sites including proline hydroxylation, methionine oxidation, arginine deamidation and glutamine deamidation are listed under "Assigned Modifications" column. Peptide location within the protein is indicated with "Start" and "End" columns. Peptides with "FALSE" under the "Is Unique" column map to an additional protein and the most probable protein is listed under "Protein Description."

| Calculated Peptide Mass (+H) | Modified Peptide | Assigned Modifications | Peptide Length | Start | End | Protein ID | Gene | Protein Description | Is Unique |
|---|---|---|---|---|---|---|---|---|---|
| 944.4935 | LGEVPTIES | | 9 | 1331 | 1339 | P46940 | IQGAP1 | Ras GTPase-activating-like protein IQGAP1 | TRUE |
| 954.5002 | NP*RP*VGLGP* | 2P(15.9949),4P(15.9949),9P(15.9949) | 9 | 199 | 207 | P0CG12 | DERPC | Decreased expression in renal and prostate cancer protein | TRUE |
| 954.5003 | GPRGEVGLP*G | 9P(15.9949) | 10 | 280 | 289 | P08123 | COL1A2 | Collagen alpha-2(I) chain | TRUE |
| 954.5043 | LVPGGPGFGPG | | 11 | 322 | 332 | P15502 | ELN | Elastin | TRUE |
| 957.471 | PVEEAPKGM | | 9 | 161 | 169 | P52565 | ARHGDIA | Rho GDP-dissociation inhibitor 1 | FALSE |
| 962.4802 | GERGAP*GFR | 6P(15.9949) | 9 | 483 | 491 | P02461 | COL3A1 | Collagen alpha-1(III) chain | TRUE |
| 962.4802 | GARGERGFP* | 9P(15.9949) | 9 | 674 | 682 | P02452 | COL1A1 | Collagen alpha-1(I) chain | TRUE |
| 963.424 | LENCPPGFS | | 9 | 2079 | 2087 | Q9H251 | CDH23 | Cadherin-23 | TRUE |
| 966.4084 | LMVGDEASE | 2M(15.9949) | 9 | 55 | 63 | P61160 | ACTR2 | Actin-related protein 2 | TRUE |
| 966.4778 | LSDDEKFL | | 8 | 7 | 14 | P35749 | MYH11 | Myosin-11 | TRUE |
| 970.5091 | LIAPVAEEE | | 9 | 8 | 16 | P07195 | LDHB | L-lactate dehydrogenase B chain | TRUE |
| 981.4312 | GPPSAGFDFS | | 10 | 1190 | 1199 | P02452 | COL1A1 | Collagen alpha-1(I) chain | TRUE |
| 984.5221 | GPVGAAGATGAR | | 12 | 331 | 342 | P08123 | COL1A2 | Collagen alpha-2(I) chain | TRUE |
| 1001.5122 | GSRGERGLP*G | 9P(15.9949) | 10 | 877 | 886 | P08123 | COL1A2 | Collagen alpha-2(I) chain | TRUE |
| 1017.4636 | EGAP*GKWQQ | 4P(15.9949),9Q(0.9840) | 9 | 477 | 485 | P05060 | CHGB | Secretogranin-1 | TRUE |
| 1017.4708 | GARGNDGATGAA | | 12 | 320 | 331 | P02452 | COL1A1 | Collagen alpha-1(I) chain | TRUE |
| 1017.4735 | LADDVDLEQ | | 9 | 390 | 398 | P55072 | VCP | Transitional endoplasmic reticulum ATPase | TRUE |
| 1019.5016 | GQRGERGFP* | 9P(15.9949) | 9 | 965 | 973 | P02452 | COL1A1 | Collagen alpha-1(I) chain | FALSE |
| 1032.4704 | GATGDRGEAGAA | | 12 | 688 | 699 | P08123 | COL1A2 | Collagen alpha-2(I) chain | TRUE |
| 1032.4745 | PAQDFGQAGAA | | 11 | 1016 | 1026 | O95428 | PAPLN | Papilin | TRUE |
| 1032.4778 | AGRP*GEP*GLM | 10M(15.9949),4P(15.9949),7P(15.9949) | 10 | 438 | 447 | P08123 | COL1A2 | Collagen alpha-2(I) chain | TRUE |
| 1034.5741 | GPVGPVGARGPA | | 12 | 1076 | 1087 | P02452 | COL1A1 | Collagen alpha-1(I) chain | TRUE |
| 1034.5741 | GPVGPAGAVGPR | | 12 | 985 | 996 | P08123 | COL1A2 | Collagen alpha-2(I) chain | TRUE |
| 1034.5741 | GPAGPVGPVGAR | | 12 | 1073 | 1084 | P02452 | COL1A1 | Collagen alpha-1(I) chain | TRUE |
| 1039.4577 | IGVADADYSE | | 10 | 1167 | 1176 | Q05707 | COL14A1 | Collagen alpha-1(XIV) chain | TRUE |
| 1039.4579 | FSDDVQVET | | 9 | 284 | 292 | Q92783 | STAM | Signal transducing adapter molecule 1 | TRUE |
| 1040.4754 | GPAGSRGDGGP*P | 11P(15.9949) | 12 | 772 | 783 | P08123 | COL1A2 | Collagen alpha-2(I) chain | TRUE |
| 1040.4754 | GPAGSRGDGGPP* | 12P(15.9949) | 12 | 772 | 783 | P08123 | COL1A2 | Collagen alpha-2(I) chain | TRUE |
| 1041.5363 | LASNPAPFPAG | | 11 | 79 | 89 | P0CG12 | DERPC | Decreased expression in renal and prostate cancer protein | TRUE |
| 1041.5436 | GPVGAAGATGARG | | 13 | 331 | 343 | P08123 | COL1A2 | Collagen alpha-2(I) chain | TRUE |
| 1043.4751 | GPSGESRPGSP* | 11P(15.9949) | 11 | 1719 | 1729 | Q05707 | COL14A1 | Collagen alpha-1(XIV) chain | TRUE |
| 1043.4864 | GPAGARGNDGAT | | 12 | 317 | 328 | P02452 | COL1A1 | Collagen alpha-1(I) chain | TRUE |
| 1043.4903 | GAPGAP*GHPGP*P | 11P(15.9949),6P(15.9949) | 12 | 1041 | 1052 | P02461 | COL3A1 | Collagen alpha-1(III) chain | TRUE |
| 1043.4903 | GAPGAP*GHPGPP* | 12P(15.9949),6P(15.9949) | 12 | 1041 | 1052 | P02461 | COL3A1 | Collagen alpha-1(III) chain | TRUE |
| 1043.4903 | GAPGAPGHP*GP*P | 11P(15.9949),9P(15.9949) | 12 | 1041 | 1052 | P02461 | COL3A1 | Collagen alpha-1(III) chain | TRUE |
| 1054.4575 | LEGEFTEET | | 9 | 400 | 408 | P20908 | COL5A1 | Collagen alpha-1(V) chain | TRUE |
| 1058.5224 | GARGSDGSVGPV | | 12 | 232 | 243 | P08123 | COL1A2 | Collagen alpha-2(I) chain | TRUE |
| 1058.5265 | GPTGAVGFAGPQ | | 12 | 840 | 851 | P05997 | COL5A2 | Collagen alpha-2(V) chain | TRUE |
| 1058.5265 | LADSFHLQQ | | 9 | 569 | 577 | Q13813 | SPTAN1 | Spectrin alpha chain, non-erythrocytic 1 | TRUE |
| 1068.5067 | GPAGERGSP*GPA | 9P(15.9949) | 12 | 506 | 517 | P02452 | COL1A1 | Collagen alpha-1(I) chain | TRUE |
| 1068.5067 | GPP*GSRGSP*GAP | 3P(15.9949),9P(15.9949) | 12 | 1242 | 1253 | Q01955 | COL4A3 | Collagen alpha-3(IV) chain | TRUE |
| 1068.5067 | GPAGSRGAP*GPQ | 12Q(0.9840),9P(15.9949) | 12 | 1077 | 1088 | P02461 | COL3A1 | Collagen alpha-1(III) chain | TRUE |
| 1068.5067 | GPAGSP*GERGPA | 6P(15.9949) | 12 | 1084 | 1095 | P20908 | COL5A1 | Collagen alpha-1(V) chain | TRUE |
| 1068.5067 | GPAGERGSPGP*A | 11P(15.9949) | 12 | 506 | 517 | P02452 | COL1A1 | Collagen alpha-1(I) chain | TRUE |
| 1068.5068 | PAGERGEQGPA | | 11 | 627 | 637 | P02452 | COL1A1 | Collagen alpha-1(I) chain | TRUE |
| 1076.5482 | GATGFP*GAAGRV | 6P(15.9949) | 12 | 872 | 883 | P02452 | COL1A1 | Collagen alpha-1(I) chain | FALSE |
| 1082.6316 | IAGQRGVVGLP* | 11P(15.9949) | 11 | 954 | 964 | P02452 | COL1A1 | Collagen alpha-1(I) chain | TRUE |
| 1082.6316 | PLGIAGITGARG | | 12 | 946 | 957 | P02461 | COL3A1 | Collagen alpha-1(III) chain | TRUE |
| 1082.6316 | GPLGIAGITGAR | | 12 | 945 | 956 | P02461 | COL3A1 | Collagen alpha-1(III) chain | TRUE |
| 1096.4653 | GPDGNNGAQGP*P | 11P(15.9949) | 12 | 523 | 534 | P08123 | COL1A2 | Collagen alpha-2(I) chain | TRUE |
| 1096.4653 | GPDGNNGAQGPP* | 12P(15.9949) | 12 | 523 | 534 | P08123 | COL1A2 | Collagen alpha-2(I) chain | TRUE |
| 1098.5135 | VLSGGTTMYPG | 8M(15.9949) | 11 | 300 | 310 | P62736 | ACTA2 | Actin, aortic smooth muscle | FALSE |
| 1098.5141 | IAFYSYTFS | | 9 | 336 | 344 | Q9UJ70 | NAGK | N-acetyl-D-glucosamine kinase | TRUE |
| 1098.5173 | GPQGESRTGP*P | 10P(15.9949) | 11 | 3004 | 3014 | Q99715 | COL12A1 | Collagen alpha-1(XII) chain | TRUE |
| 1098.5173 | GPQGESRTGPP* | 11P(15.9949) | 11 | 3004 | 3014 | Q99715 | COL12A1 | Collagen alpha-1(XII) chain | TRUE |
| 1098.5901 | GPVGRTGEVGAV | | 12 | 826 | 837 | P08123 | COL1A2 | Collagen alpha-2(I) chain | TRUE |
| 1098.5901 | GARGLP*GTAGLP* | 12P(15.9949),6P(15.9949) | 12 | 251 | 262 | P02452 | COL1A1 | Collagen alpha-1(I) chain | TRUE |
| 1102.5626 | LSADEVELGLG | | 11 | 209 | 219 | Q3LXA3 | TKFC | Triokinase/FMN cyclase | TRUE |
| 1102.5638 | GIAGAP*GFP*GAR | 6P(15.9949),9P(15.9949) | 12 | 404 | 415 | P02452 | COL1A1 | Collagen alpha-1(I) chain | TRUE |
| 1106.5112 | GFAGEKGPSGEA | | 12 | 841 | 852 | P08123 | COL1A2 | Collagen alpha-2(I) chain | TRUE |
| 1106.5224 | GSRGFP*GADGVA | 6P(15.9949) | 12 | 491 | 502 | P02452 | COL1A1 | Collagen alpha-1(I) chain | TRUE |
| 1106.5226 | FSGYLLYPM | 9M(15.9949) | 9 | 736 | 744 | P27658 | COL8A1 | Collagen alpha-1(VIII) chain | TRUE |
| 1106.5226 | FSGYLLYP*M | 8P(15.9949) | 9 | 736 | 744 | P27658 | COL8A1 | Collagen alpha-1(VIII) chain | TRUE |
| 1113.5097 | LYAYEPADTA | | 10 | 149 | 158 | O95994 | AGR2 | Anterior gradient protein 2 homolog | TRUE |
| 1113.5169 | GPSGEP*GKQGPS | 6P(15.9949) | 12 | 977 | 988 | P02452 | COL1A1 | Collagen alpha-1(I) chain | TRUE |
| 1113.5169 | GPSGEP*GKQGAP* | 12P(15.9949),6P(15.9949) | 12 | 999 | 1010 | P02458 | COL2A1 | Collagen alpha-1(II) chain | TRUE |
| 1113.521 | GPP*GIP*GFDGAP | 3P(15.9949),6P(15.9949) | 12 | 1393 | 1404 | P02462 | COL4A1 | Collagen alpha-1(IV) chain | TRUE |
| 1113.521 | GP*PGIP*GFDGAP | 2P(15.9949),6P(15.9949) | 12 | 1393 | 1404 | P02462 | COL4A1 | Collagen alpha-1(IV) chain | TRUE |
| 1115.5367 | LWEIENNPT | | 9 | 86 | 94 | P51858 | HDGF | Hepatoma-derived growth factor | TRUE |
| 1115.5439 | GAKGDRGETGPA | | 12 | 1031 | 1042 | P02452 | COL1A1 | Collagen alpha-1(I) chain | TRUE |
| 1120.5707 | MLPANYVEAI | | 10 | 8516 | 8525 | P20929 | NEB | Nebulin | TRUE |
| 1120.5744 | GPRGLP*GPP*GAP* | 12P(15.9949),6P(15.9949),9P(15.9949) | 12 | 185 | 196 | P02452 | COL1A1 | Collagen alpha-1(I) chain | TRUE |
| 1120.5844 | LTANITKSGNT | 4N(0.9840) | 11 | 202 | 212 | P01877 | IGHA2 | Immunoglobulin heavy constant alpha 2 | FALSE |
| 1122.5459 | LTGTINSSMQA | | 11 | 495 | 505 | Q9Y490 | TLN1 | Talin-1 | TRUE |
| 1122.5465 | ILGQWFEET | | 9 | 590 | 598 | Q5T4S7 | UBR4 | E3 ubiquitin-protein ligase UBR4 | TRUE |
| 1122.5537 | GPAGRP*GEVGP*P | 11P(15.9949),6P(15.9949) | 12 | 914 | 925 | P02452 | COL1A1 | Collagen alpha-1(I) chain | TRUE |
| 1122.5537 | GPAGRP*GEVGPP* | 12P(15.9949),6P(15.9949) | 12 | 914 | 925 | P02452 | COL1A1 | Collagen alpha-1(I) chain | TRUE |
| 1125.5282 | GPAGERGSP*GPAG | 9P(15.9949) | 13 | 506 | 518 | P02452 | COL1A1 | Collagen alpha-1(I) chain | TRUE |
| 1125.5283 | GPAGERGEQGPA | | 12 | 626 | 637 | P02452 | COL1A1 | Collagen alpha-1(I) chain | TRUE |
| 1125.5283 | GPAGERGEQGAP | | 12 | 648 | 659 | P02458 | COL2A1 | Collagen alpha-1(II) chain | TRUE |
| 1125.5283 | GPAGPAGERGEQ | | 12 | 623 | 634 | P02452 | COL1A1 | Collagen alpha-1(I) chain | FALSE |
| 1128.5278 | GSP*GERGEVGPA | 3P(15.9949) | 12 | 709 | 720 | P08123 | COL1A2 | Collagen alpha-2(I) chain | TRUE |
| 1128.532 | GPP*GLP*GFAGNP* | 12P(15.9949),3P(15.9949),6P(15.9949) | 12 | 136 | 147 | P02462 | COL4A1 | Collagen alpha-1(IV) chain | TRUE |
| 1131.5276 | AGEKGPSGEAGTA | | 13 | 843 | 855 | P08123 | COL1A2 | Collagen alpha-2(I) chain | TRUE |
| 1131.5316 | LADDHDVLSF | | 10 | 253 | 262 | P49257 | LMAN1 | Protein ERGIC-53 | FALSE |
| 1131.5388 | GPAGKSGDRGET | | 12 | 1058 | 1069 | P02452 | COL1A1 | Collagen alpha-1(I) chain | TRUE |
| 1131.5428 | GDDAPRAVFPS | | 11 | 25 | 35 | P62736 | ACTA2 | Actin, aortic smooth muscle | FALSE |
| 1133.5261 | DDHFLFDKP | | 9 | 189 | 197 | P12277 | CKB | Creatine kinase B-type | FALSE |
| 1137.5282 | GPAGQDGRP*GP*P | 11P(15.9949),9P(15.9949) | 12 | 557 | 568 | P02452 | COL1A1 | Collagen alpha-1(I) chain | TRUE |
| 1139.5915 | GPRGRTGDAGPV | | 12 | 1166 | 1177 | P02452 | COL1A1 | Collagen alpha-1(I) chain | TRUE |
| 1139.6531 | GIAGQRGVVGLP* | 12P(15.9949) | 12 | 953 | 964 | P02452 | COL1A1 | Collagen alpha-1(I) chain | TRUE |
| 1139.6531 | GPLGIAGITGARG | | 13 | 945 | 957 | P02461 | COL3A1 | Collagen alpha-1(III) chain | TRUE |
| 1139.6531 | IAGQRGVVGLP*G | 11P(15.9949) | 12 | 954 | 965 | P02452 | COL1A1 | Collagen alpha-1(I) chain | TRUE |
| 1142.5071 | GPP*GESGREGAP* | 12P(15.9949),3P(15.9949) | 12 | 1007 | 1018 | P02452 | COL1A1 | Collagen alpha-1(I) chain | TRUE |
| 1142.5071 | GP*PGESGREGAP* | 12P(15.9949),2P(15.9949) | 12 | 1007 | 1018 | P02452 | COL1A1 | Collagen alpha-1(I) chain | TRUE |
| 1147.5047 | GSQGAP*GLQGMP* | 11M(15.9949),12P(15.9949),6P(15.9949) | 12 | 719 | 730 | P02452 | COL1A1 | Collagen alpha-1(I) chain | TRUE |
| 1148.6058 | LRVAPEEHPT | | 10 | 96 | 105 | P62736 | ACTA2 | Actin, aortic smooth muscle | FALSE |
| 1154.6163 | GPRGEVGLP*GLS | 9P(15.9949) | 12 | 280 | 291 | P08123 | COL1A2 | Collagen alpha-2(I) chain | TRUE |
| 1157.5108 | VNENLENYY | | 9 | 272 | 280 | P51884 | LUM | Lumican | TRUE |
| 1157.518 | GSQGESGRPGP*P* | 11P(15.9949),12P(15.9949) | 12 | 555 | 566 | P02461 | COL3A1 | Collagen alpha-1(III) chain | TRUE |
| 1157.518 | GSQGESGRP*GP*P | 11P(15.9949),9P(15.9949) | 12 | 555 | 566 | P02461 | COL3A1 | Collagen alpha-1(III) chain | TRUE |

**Supplemental Table 2 (3/4): Matched peptide library.** Peptides from the liquid chromatography tandem mass spectrometry peptide library whose calculated protonated peptide mass is within 8 ppm of a calibrated matrix-assisted laser/desorption ionization mass to charge ratio. The "Modified Peptide" column includes peptide sequence with proline hydroxylation sites indicated with an asterisk. All detectable modification sites including proline hydroxylation, methionine oxidation, arginine deamidation and glutamine deamidation are listed under "Assigned Modifications" column. Peptide location within the protein is indicated with "Start" and "End" columns. Peptides with "FALSE" under the "Is Unique" column map to an additional protein and the most probable protein is listed under "Protein Description."

| Calculated Peptide Mass (+H) | Modified Peptide | Assigned Modifications | Peptide Length | Start | End | Protein ID | Gene | Protein Description | Is Unique |
|---|---|---|---|---|---|---|---|---|---|
| 1157.518 | GRDGNPGSDGLP* | 12P(15.9949) | 12 | 1011 | 1022 | P02461 | COL3A1 | Collagen alpha-1(III) chain | TRUE |
| 1157.518 | GSQGESGRP*GPP* | 12P(15.9949),9P(15.9949) | 12 | 555 | 566 | P02461 | COL3A1 | Collagen alpha-1(III) chain | TRUE |
| 1157.5221 | GPQGFQGPAGEP* | 12P(15.9949) | 12 | 109 | 120 | P08123 | COL1A2 | Collagen alpha-2(I) chain | TRUE |
| 1161.5758 | GAAGERGAP*GFR | 9P(15.9949) | 12 | 480 | 491 | P02461 | COL3A1 | Collagen alpha-1(III) chain | TRUE |
| 1161.5786 | LEPGQEYNVL | | 10 | 1043 | 1052 | P24821 | TNC | Tenascin | TRUE |
| 1163.5625 | GARGQAGVMGFP* | 12P(15.9949) | 12 | 572 | 583 | P02452 | COL1A1 | Collagen alpha-1(I) chain | TRUE |
| 1171.5449 | GARGAP*GDRGEP* | 12P(15.9949),6P(15.9949) | 12 | 794 | 805 | P02452 | COL1A1 | Collagen alpha-1(I) chain | TRUE |
| 1172.5177 | GSP*GERGETGP*P | 11P(15.9949),3P(15.9949) | 12 | 795 | 806 | P02461 | COL3A1 | Collagen alpha-1(III) chain | TRUE |
| 1172.5177 | GSP*GERGETGPP* | 12P(15.9949),3P(15.9949) | 12 | 795 | 806 | P02461 | COL3A1 | Collagen alpha-1(III) chain | TRUE |
| 1173.5129 | GRDGNP*GSDGLP* | 12P(15.9949),6P(15.9949) | 12 | 1011 | 1022 | P02461 | COL3A1 | Collagen alpha-1(III) chain | TRUE |
| 1179.5573 | GARGQAGVMGFP* | 12P(15.9949),9M(15.9949) | 12 | 572 | 583 | P02452 | COL1A1 | Collagen alpha-1(I) chain | TRUE |
| 1185.5493 | GSEGPQGVRGEP* | 12P(15.9949) | 12 | 362 | 373 | P02452 | COL1A1 | Collagen alpha-1(I) chain | TRUE |
| 1185.5493 | GPTGQVGDP*GER | 9P(15.9949) | 12 | 487 | 498 | P20908 | COL5A1 | Collagen alpha-1(V) chain | TRUE |
| 1185.5494 | GQPGAKGEQGEAG | | 13 | 843 | 855 | P02458 | COL2A1 | Collagen alpha-1(II) chain | TRUE |
| 1188.5531 | LAAQYEHDLE | | 10 | 193 | 202 | P62826 | RAN | GTP-binding nuclear protein Ran | TRUE |
| 1188.5603 | GADGSP*GKDGVRG | 6P(15.9949) | 13 | 752 | 764 | P02452 | COL1A1 | Collagen alpha-1(I) chain | TRUE |
| 1203.575 | GARGEP*GNIGFP* | 12P(15.9949),6P(15.9949) | 12 | 484 | 495 | P08123 | COL1A2 | Collagen alpha-2(I) chain | TRUE |
| 1203.5864 | GPSGARGERGFP* | 12P(15.9949) | 12 | 671 | 682 | P02452 | COL1A1 | Collagen alpha-1(I) chain | TRUE |
| 1205.5656 | GERGGP*GSRGFP* | 12P(15.9949),6P(15.9949) | 12 | 485 | 496 | P02452 | COL1A1 | Collagen alpha-1(I) chain | TRUE |
| 1211.6125 | GPRGERGEAGIP* | 12P(15.9949) | 12 | 444 | 455 | P02461 | COL3A1 | Collagen alpha-1(III) chain | TRUE |
| 1211.6126 | GAP*GERGRP*GLP* | 12P(15.9949),3P(15.9949),9P(15.9949) | 12 | 303 | 314 | P02461 | COL3A1 | Collagen alpha-1(III) chain | TRUE |
| 1211.6126 | GLP*GERGRP*GAP* | 12P(15.9949),3P(15.9949),9P(15.9949) | 12 | 305 | 316 | P02452 | COL1A1 | Collagen alpha-1(I) chain | TRUE |
| 1211.6167 | TLPHPNLHGPE | | 11 | 2090 | 2100 | P02751 | FN1 | Fibronectin | TRUE |
| 1226.6123 | GPAGEVGKP*GERG | 9P(15.9949) | 13 | 562 | 574 | P08123 | COL1A2 | Collagen alpha-2(I) chain | TRUE |
| 1242.582 | GPAGARGNDGATGAA | | 15 | 317 | 331 | P02452 | COL1A1 | Collagen alpha-1(I) chain | TRUE |
| 1257.5818 | GPAGATGDRGEAGAA | | 15 | 685 | 699 | P08123 | COL1A2 | Collagen alpha-2(I) chain | TRUE |
| 1271.5498 | GSP*GEQGPSGASGPA | 3P(15.9949) | 15 | 1124 | 1138 | P02452 | COL1A1 | Collagen alpha-1(I) chain | TRUE |
| 1280.5428 | GP*P*GP*PSAGFDFS | 2P(15.9949),3P(15.9949),5P(15.9949) | 13 | 1187 | 1199 | P02452 | COL1A1 | Collagen alpha-1(I) chain | TRUE |
| 1280.5428 | GP*P*GPP*SAGFDFS | 2P(15.9949),3P(15.9949),6P(15.9949) | 13 | 1187 | 1199 | P02452 | COL1A1 | Collagen alpha-1(I) chain | TRUE |
| 1280.6816 | GPRGLP*GERGRP* | 12P(15.9949),6P(15.9949) | 12 | 302 | 313 | P02452 | COL1A1 | Collagen alpha-1(I) chain | TRUE |
| 1280.6816 | GRPGRP*GERGLP* | 12P(15.9949),6P(15.9949) | 12 | 234 | 245 | P02461 | COL3A1 | Collagen alpha-1(III) chain | TRUE |
| 1283.6338 | GPAGARGSDGSVGPV | | 15 | 229 | 243 | P08123 | COL1A2 | Collagen alpha-2(I) chain | TRUE |
| 1283.6338 | GARGSDGSVGPVGPA | | 15 | 232 | 246 | P08123 | COL1A2 | Collagen alpha-2(I) chain | TRUE |
| 1283.6338 | GPAGARGESGLAGAP* | 15P(15.9949) | 15 | 989 | 1003 | P39060 | COL18A1 | Collagen alpha-1(XVIII) chain | TRUE |
| 1286.6599 | GQRGERGFPGLP* | 12P(15.9949) | 12 | 965 | 976 | P02452 | COL1A1 | Collagen alpha-1(I) chain | FALSE |
| 1286.6599 | GQRGERGFP*GLP | 9P(15.9949) | 12 | 965 | 976 | P02452 | COL1A1 | Collagen alpha-1(I) chain | FALSE |
| 1302.6548 | GQRGERGFP*GLP* | 12P(15.9949),9P(15.9949) | 12 | 965 | 976 | P02452 | COL1A1 | Collagen alpha-1(I) chain | FALSE |
| 1306.6062 | ASEFFRSGKYD | | 11 | 248 | 258 | P06733 | ENO1 | Alpha-enolase | TRUE |
| 1323.7743 | GPLGIAGITGARGLA | | 15 | 945 | 959 | P02461 | COL3A1 | Collagen alpha-1(III) chain | TRUE |
| 1324.6353 | LAVTMDDFRWA | | 11 | 445 | 455 | P55072 | VCP | Transitional endoplasmic reticulum ATPase | TRUE |
| 1328.6188 | GAP*GGKGDAGAP*GER | 12P(15.9949),3P(15.9949) | 15 | 669 | 683 | P02461 | COL3A1 | Collagen alpha-1(III) chain | TRUE |
| 1336.6603 | GGPGADGVPGKDGPR | | 15 | 747 | 761 | P02461 | COL3A1 | Collagen alpha-1(III) chain | TRUE |
| 1340.6075 | GP*AGKDGEAGAQGP*P | 14P(15.9949),2P(15.9949) | 15 | 608 | 622 | P02452 | COL1A1 | Collagen alpha-1(I) chain | TRUE |
| 1343.6588 | GREP*GP*PIYAAPS | 4P(15.9949),6P(15.9949) | 13 | 209 | 221 | O43281 | EFS | Embryonal Fyn-associated substrate | TRUE |
| 1343.67 | GATGFP*GAAGRVGP*P | 14P(15.9949),6P(15.9949) | 15 | 872 | 886 | P02452 | COL1A1 | Collagen alpha-1(I) chain | FALSE |
| 1343.67 | GIAGAP*GFP*GARGPS | 6P(15.9949),9P(15.9949) | 15 | 404 | 418 | P02452 | COL1A1 | Collagen alpha-1(I) chain | TRUE |
| 1343.67 | GATGFP*GAAGRVGPP* | 15P(15.9949),6P(15.9949) | 15 | 872 | 886 | P02452 | COL1A1 | Collagen alpha-1(I) chain | FALSE |
| 1343.67 | GIAGAP*GFPGARGP*S | 14P(15.9949),6P(15.9949) | 15 | 404 | 418 | P02452 | COL1A1 | Collagen alpha-1(I) chain | TRUE |
| 1343.6701 | GAP*GIAGAP*GFP*GAR | 12P(15.9949),3P(15.9949),9P(15.9949) | 15 | 401 | 415 | P02452 | COL1A1 | Collagen alpha-1(I) chain | TRUE |
| 1343.6701 | GATGFPGSAGRVGP*P | 14P(15.9949) | 15 | 906 | 920 | P05997 | COL5A2 | Collagen alpha-2(V) chain | TRUE |
| 1350.6396 | GPAGPAGERGEQGPA | | 15 | 623 | 637 | P02452 | COL1A1 | Collagen alpha-1(I) chain | TRUE |
| 1352.6551 | GGPGADGVP*GKDGPR | 9P(15.9949) | 15 | 747 | 761 | P02461 | COL3A1 | Collagen alpha-1(III) chain | TRUE |
| 1352.6551 | GGP*GADGVPGKDGPR | 3P(15.9949) | 15 | 747 | 761 | P02461 | COL3A1 | Collagen alpha-1(III) chain | TRUE |
| 1359.665 | GATGFP*GAAGRVGP*P* | 14P(15.9949),15P(15.9949),6P(15.9949) | 15 | 872 | 886 | P02452 | COL1A1 | Collagen alpha-1(I) chain | FALSE |
| 1359.665 | GIAGAP*GFP*GARGP*S | 14P(15.9949),6P(15.9949),9P(15.9949) | 15 | 404 | 418 | P02452 | COL1A1 | Collagen alpha-1(I) chain | TRUE |
| 1359.665 | GATGFP*GSAGRVGP*P | 14P(15.9949),6P(15.9949) | 15 | 906 | 920 | P05997 | COL5A2 | Collagen alpha-2(V) chain | TRUE |
| 1359.6651 | LSYTATDRHPQA | | 12 | 1268 | 1279 | P49327 | FASN | Fatty acid synthase | TRUE |
| 1361.6667 | GARGERGFP*GERG | 9P(15.9949) | 13 | 674 | 686 | P02452 | COL1A1 | Collagen alpha-1(I) chain | TRUE |
| 1365.6472 | GP*PGQFPFDFLQ | 2P(15.9949) | 12 | 1307 | 1318 | P39060 | COL18A1 | Collagen alpha-1(XVIII) chain | TRUE |
| 1365.6472 | GPP*GQFPFDFLQ | 3P(15.9949) | 12 | 1307 | 1318 | P39060 | COL18A1 | Collagen alpha-1(XVIII) chain | TRUE |
| 1384.6814 | GAAGEP*GKAGERGVP* | 15P(15.9949),6P(15.9949) | 15 | 587 | 601 | P02452 | COL1A1 | Collagen alpha-1(I) chain | TRUE |
| 1384.6815 | GPVGPAGKSGDRGET | | 15 | 1055 | 1069 | P02452 | COL1A1 | Collagen alpha-1(I) chain | TRUE |
| 1384.6926 | GLRGEIGNP*GRDGA | 9P(15.9949) | 14 | 661 | 674 | P08123 | COL1A2 | Collagen alpha-2(I) chain | TRUE |
| 1384.6928 | FRMTPFSAIGLE | 3M(15.9949) | 12 | 414 | 425 | P27824 | CANX | Calnexin | TRUE |
| 1386.6871 | GAAGERGAP*GFRGPA | 9P(15.9949) | 15 | 480 | 494 | P02461 | COL3A1 | Collagen alpha-1(III) chain | TRUE |
| 1386.6971 | GADGSPGKDGVRGLT | | 15 | 752 | 766 | P02452 | COL1A1 | Collagen alpha-1(I) chain | TRUE |
| 1399.6448 | GSP*GEAGRP*GEAGLP* | 15P(15.9949),3P(15.9949),9P(15.9949) | 15 | 521 | 535 | P02452 | COL1A1 | Collagen alpha-1(I) chain | TRUE |
| 1399.6561 | LFQPSFIGMESAG | 9M(15.9949) | 13 | 263 | 275 | P62736 | ACTA2 | Actin, aortic smooth muscle | FALSE |
| 1410.6719 | GPRGERGPP*GESGAA | 9P(15.9949) | 15 | 586 | 600 | P08123 | COL1A2 | Collagen alpha-2(I) chain | TRUE |
| 1410.6719 | GPRGERGP*PGESGAA | 8P(15.9949) | 15 | 586 | 600 | P08123 | COL1A2 | Collagen alpha-2(I) chain | TRUE |
| 1421.7859 | GPQGIAGQRGVVGLP* | 15P(15.9949) | 15 | 950 | 964 | P02452 | COL1A1 | Collagen alpha-1(I) chain | TRUE |
| 1426.6667 | GPTGARGAP*GDRGEP* | 15P(15.9949),9P(15.9949) | 15 | 791 | 805 | P02452 | COL1A1 | Collagen alpha-1(I) chain | TRUE |
| 1426.6667 | TGARGAP*GDRGEPGP* | 15P(15.9949),7P(15.9949) | 15 | 793 | 807 | P02452 | COL1A1 | Collagen alpha-1(I) chain | TRUE |
| 1426.6667 | TGARGAPGDRGEP*GP* | 13P(15.9949),15P(15.9949) | 15 | 793 | 807 | P02452 | COL1A1 | Collagen alpha-1(I) chain | TRUE |
| 1438.7032 | GPRGSP*GERGEVGPA | 6P(15.9949) | 15 | 706 | 720 | P08123 | COL1A2 | Collagen alpha-2(I) chain | TRUE |
| 1438.7032 | GP*RGSPGERGEVGPA | 2P(15.9949) | 15 | 706 | 720 | P08123 | COL1A2 | Collagen alpha-2(I) chain | TRUE |
| 1441.5937 | GEAGRDGNP*GNDGP*P | 14P(15.9949),9P(15.9949) | 15 | 922 | 936 | P08123 | COL1A2 | Collagen alpha-2(I) chain | TRUE |
| 1446.6793 | GPP*GARGQAGVMGFP* | 12M(15.9949),15P(15.9949),3P(15.9949) | 15 | 569 | 583 | P02452 | COL1A1 | Collagen alpha-1(I) chain | TRUE |
| 1446.6793 | GP*PGARGQAGVMGFP* | 12M(15.9949),15P(15.9949),2P(15.9949) | 15 | 569 | 583 | P02452 | COL1A1 | Collagen alpha-1(I) chain | TRUE |
| 1446.6793 | P*QGARGQPGVMGFP* | 11M(15.9949),14P(15.9949),1P(15.9949) | 14 | 592 | 605 | P02458 | COL2A1 | Collagen alpha-1(II) chain | TRUE |
| 1446.682 | LDYGFVTPLTSMS | 12M(15.9949) | 13 | 598 | 610 | P19827 | ITIH1 | Inter-alpha-trypsin inhibitor heavy chain H1 | TRUE |
| 1446.6899 | AFFDTASTGKTFPG | | 14 | 518 | 531 | P02671 | FGA | Fibrinogen alpha chain | TRUE |
| 1458.7004 | GLQGMP*GERGAAGLP* | 15P(15.9949),5M(15.9949),6P(15.9949) | 15 | 725 | 739 | P02452 | COL1A1 | Collagen alpha-1(I) chain | TRUE |
| 1460.6798 | GLQGMP*GERGGLGSP* | 15P(15.9949),5M(15.9949),6P(15.9949) | 15 | 723 | 737 | P02461 | COL3A1 | Collagen alpha-1(III) chain | TRUE |
| 1472.6876 | GPP*GERGGP*GSRGFP* | 15P(15.9949),3P(15.9949),9P(15.9949) | 15 | 482 | 496 | P02452 | COL1A1 | Collagen alpha-1(I) chain | TRUE |
| 1480.743 | FVNDIFERIAGEA | | 13 | 66 | 78 | P58876 | H2BC5 | Histone H2B type 1-D | FALSE |
| 1480.7501 | GPRGERGEAGIP*GVP* | 12P(15.9949),15P(15.9949) | 15 | 444 | 458 | P02461 | COL3A1 | Collagen alpha-1(III) chain | TRUE |
| 1480.7501 | RGPEGSRGLP*GVEGP | 10P(15.9949) | 15 | 714 | 728 | P20849 | COL9A1 | Collagen alpha-1(IX) chain | TRUE |
| 1483.7246 | GPAGEEGKRGARGEP* | 15P(15.9949) | 15 | 461 | 475 | P02452 | COL1A1 | Collagen alpha-1(I) chain | FALSE |
| 1488.6682 | GRDGVP*GGPGMRGMP | 11M(15.9949),14M(15.9949),6P(15.9949) | 15 | 525 | 539 | P02461 | COL3A1 | Collagen alpha-1(III) chain | TRUE |
| 1488.6682 | GRDGVP*GGP*GMRGMP | 14M(15.9949),6P(15.9949),9P(15.9949) | 15 | 525 | 539 | P02461 | COL3A1 | Collagen alpha-1(III) chain | TRUE |
| 1495.7247 | GPRGSEGPQGVRGEP* | 15P(15.9949) | 15 | 359 | 373 | P02452 | COL1A1 | Collagen alpha-1(I) chain | TRUE |
| 1502.7235 | IFSGFLLFPDMEA | 11M(15.9949) | 13 | 241 | 253 | P02746 | C1QB | Complement C1q subcomponent subunit B | TRUE |
| 1502.7307 | GLQGVIGFPGMQGPE | 11M(15.9949) | 15 | 55 | 69 | P02462 | COL4A1 | Collagen alpha-1(IV) chain | TRUE |
| 1508.7451 | GPAGARGSDGSVGPVGPA | | 18 | 229 | 246 | P08123 | COL1A2 | Collagen alpha-2(I) chain | TRUE |
| 1521.675 | GPRGEQGFMGNTGPT | 9M(15.9949) | 15 | 1347 | 1361 | P08572 | COL4A2 | Collagen alpha-2(IV) chain | TRUE |
| 1521.7879 | GPRGAP*GERGRP*GLP* | 12P(15.9949),15P(15.9949),6P(15.9949) | 15 | 300 | 314 | P02461 | COL3A1 | Collagen alpha-1(III) chain | TRUE |
| 1521.7879 | GPRGLP*GERGRP*GAP* | 12P(15.9949),15P(15.9949),6P(15.9949) | 15 | 302 | 316 | P02452 | COL1A1 | Collagen alpha-1(I) chain | TRUE |
| 1521.7879 | P*GKNGDKGHAGLAGAR | 1P(15.9949) | 16 | 504 | 519 | P08123 | COL1A2 | Collagen alpha-2(I) chain | TRUE |
| 1528.7013 | LKPSEAPELDEDEG | | 14 | 67 | 80 | P33241 | LSP1 | Lymphocyte-specific protein 1 | TRUE |
| 1540.7937 | GLRGEIGNP*GRDGAR | 9P(15.9949) | 15 | 661 | 675 | P08123 | COL1A2 | Collagen alpha-2(I) chain | TRUE |

**Supplemental Table 2 (4/4): Matched peptide library.** Peptides from the liquid chromatography tandem mass spectrometry peptide library whose calculated protonated peptide mass is within 8 ppm of a calibrated matrix-assisted laser/desorption ionization mass to charge ratio. The "Modified Peptide" column includes peptide sequence with proline hydroxylation sites indicated with an asterisk. All detectable modification sites including proline hydroxylation, methionine oxidation, arginine deamidation and glutamine deamidation are listed under "Assigned Modifications" column. Peptide location within the protein is indicated with "Start" and "End" columns. Peptides with "FALSE" under the "Is Unique" column map to an additional protein and the most probable protein is listed under "Protein Description."

| Calculated Peptide Mass (+H) | Modified Peptide | Assigned Modifications | Peptide Length | Start | End | Protein ID | Gene | Protein Description | Is Unique |
|---|---|---|---|---|---|---|---|---|---|
| 1540.8038 | LGTIAKSGTKAFMEA | 13M(15.9949) | 15 | 102 | 116 | P08238 | HSP90AB1 | Heat shock protein HSP 90-beta | FALSE |
| 1588.7925 | ITEANEKTREAQQA | | 14 | 1398 | 1411 | P11047 | LAMC1 | Laminin subunit gamma-1 | TRUE |
| 1588.7937 | GARGERGFP*GERGVQ | 9P(15.9949) | 15 | 674 | 688 | P02452 | COL1A1 | Collagen alpha-1(I) chain | TRUE |
| 1609.7928 | GPVGPAGKSGDRGETGPA | | 18 | 1055 | 1072 | P02452 | COL1A1 | Collagen alpha-1(I) chain | TRUE |
| 1609.7967 | GP*PGFPGAVGAKGEAGPQ | 2P(15.9949) | 18 | 341 | 358 | P02452 | COL1A1 | Collagen alpha-1(I) chain | TRUE |
| 1609.8041 | GPRGDQGPVGRTGEVGA | | 17 | 820 | 836 | P08123 | COL1A2 | Collagen alpha-2(I) chain | TRUE |
| 1610.792 | GP*PGATGFPGSAGRVGP*P | 17P(15.9949),2P(15.9949) | 18 | 903 | 920 | P05997 | COL5A2 | Collagen alpha-2(V) chain | TRUE |
| 1610.7921 | GP*PGATGFP*GAAGRVGP*P | 17P(15.9949),2P(15.9949),9P(15.9949) | 18 | 869 | 886 | P02452 | COL1A1 | Collagen alpha-1(I) chain | FALSE |
| 1622.8132 | LLEADVAAHQDRIDG | | 15 | 722 | 736 | Q13813 | SPTAN1 | Spectrin alpha chain, non-erythrocytic 1 | TRUE |
| 1622.8141 | MAGRIYISGMAPRPS | 10M(15.9949) | 15 | 350 | 364 | P04004 | VTN | Vitronectin | TRUE |
| 1622.8244 | GPAGKEGGKGPRGETGPA | | 18 | 899 | 916 | P02452 | COL1A1 | Collagen alpha-1(I) chain | TRUE |
| 1629.7323 | GAAGARGNDGARGSDGQP* | 18P(15.9949) | 18 | 315 | 332 | P02461 | COL3A1 | Collagen alpha-1(III) chain | TRUE |
| 1639.8073 | PQGEQIQINFTHVE | | 14 | 741 | 754 | O60494 | CUBN | Cubilin | TRUE |
| 1639.8073 | FQPEANPSHLTLNTA | | 15 | 1362 | 1376 | Q8WX93 | PALLD | Palladin | TRUE |
| 1639.8226 | FSDYPPLGRFAVRD | | 14 | 415 | 428 | P68104 | EEF1A1 | Elongation factor 1-alpha 1 | FALSE |
| 1656.7571 | GPSGEEGKRGPNGEAGSA | | 18 | 373 | 390 | P08123 | COL1A2 | Collagen alpha-2(I) chain | TRUE |
| 1670.7727 | GPSGEP*GKQGPSGASGER | 6P(15.9949) | 18 | 977 | 994 | P02452 | COL1A1 | Collagen alpha-1(I) chain | TRUE |
| 1670.7727 | QP*QSQAAAAGAAAGGQQAA | 17Q(0.9840),2P(15.9949) | 19 | 535 | 553 | Q8NFD5 | ARID1B | AT-rich interactive domain-containing protein 1B | TRUE |
| 1675.7782 | GPAGRDGAPGNPGERGPP* | 11N(0.9840),18P(15.9949) | 18 | 1075 | 1092 | Q8NFW1 | COL22A1 | Collagen alpha-1(XXII) chain | TRUE |
| 1681.81 | LKDMAIATGGAVFGEEG | 4M(15.9949) | 17 | 313 | 329 | P10809 | HSPD1 | 60 kDa heat shock protein, mitochondrial | TRUE |
| 1681.8138 | GPKGSP*GEAGRP*GEAGLP* | 12P(15.9949),18P(15.9949),6P(15.9949) | 18 | 518 | 535 | P02452 | COL1A1 | Collagen alpha-1(I) chain | TRUE |
| 1681.8139 | SAVDNGSLANTPHLRE | 5N(0.9840) | 16 | 258 | 273 | P07585 | DCN | Decorin | TRUE |
| 1692.8034 | LGSSEADQDGLASTVRS | | 17 | 458 | 474 | Q07065 | CKAP4 | Cytoskeleton-associated protein 4 | TRUE |
| 1692.8047 | GPAGVRGPNGDAGRP*GEP* | 15P(15.9949),18P(15.9949) | 18 | 427 | 444 | P08123 | COL1A2 | Collagen alpha-2(I) chain | TRUE |
| 1703.7904 | LGPTRPSGEAETTQME | | 16 | 2809 | 2824 | Q7Z6Z7 | HUWE1 | E3 ubiquitin-protein ligase HUWE1 | TRUE |
| 1708.8724 | GPRGDQGPVGRTGEVGAV | | 18 | 820 | 837 | P08123 | COL1A2 | Collagen alpha-2(I) chain | TRUE |
| 1731.7639 | GRDGSP*GGKGDRGENGSP* | 18P(15.9949),6P(15.9949) | 18 | 1023 | 1040 | P02461 | COL3A1 | Collagen alpha-1(III) chain | TRUE |
| 1767.9206 | GQTGARGLP*GERGRVGAP* | 18P(15.9949),9P(15.9949) | 18 | 211 | 228 | P08123 | COL1A2 | Collagen alpha-2(I) chain | TRUE |
| 1772.7945 | GPP*GERGGP*GSRGFP*GADG | 15P(15.9949),3P(15.9949),9P(15.9949) | 19 | 482 | 500 | P02452 | COL1A1 | Collagen alpha-1(I) chain | TRUE |
| 1781.8999 | GLRGEIGNP*GRDGARGAP* | 18P(15.9949),9P(15.9949) | 18 | 661 | 678 | P08123 | COL1A2 | Collagen alpha-2(I) chain | TRUE |
| 1822.8261 | LIDLEDETIDAEVMNS | 14M(15.9949) | 16 | 429 | 444 | P55072 | VCP | Transitional endoplasmic reticulum ATPase | TRUE |
| 1829.9 | GPSGARGERGFP*GERGVQ | 12P(15.9949) | 18 | 671 | 688 | P02452 | COL1A1 | Collagen alpha-1(I) chain | TRUE |
| 1829.9 | SGARGERGFP*GERGVQGP | 10P(15.9949) | 18 | 673 | 690 | P02452 | COL1A1 | Collagen alpha-1(I) chain | TRUE |
| 1864.0902 | LLSPGSVDPLTRLVLVNA | | 18 | 149 | 166 | P35237 | SERPINB6 | Serpin B6 | TRUE |
| 1924.901 | IITNWDDMEKIWHHS | | 15 | 77 | 91 | P62736 | ACTA2 | Actin, aortic smooth muscle | FALSE |
| 1924.901 | IVTNWDDMEKIWHHT | | 15 | 75 | 89 | P63261 | ACTG1 | Actin, cytoplasmic 2 | FALSE |
| 1942.9 | GPP*GERGGP*GSRGFP*GADGVA | 15P(15.9949),3P(15.9949),9P(15.9949) | 21 | 482 | 502 | P02452 | COL1A1 | Collagen alpha-1(I) chain | TRUE |
| 1942.9 | GP*PGERGGP*GSRGFP*GADGVA | 15P(15.9949),2P(15.9949),9P(15.9949) | 21 | 482 | 502 | P02452 | COL1A1 | Collagen alpha-1(I) chain | TRUE |
| 1942.9 | GP*P*GERGGPGSRGFP*GADGVA | 15P(15.9949),2P(15.9949),3P(15.9949) | 21 | 482 | 502 | P02452 | COL1A1 | Collagen alpha-1(I) chain | TRUE |
| 1942.914 | GEP*GRFGVNSSDVPGP*AGLP | 16P(15.9949),3P(15.9949),9N(0.9840) | 20 | 918 | 937 | P39060 | COL18A1 | Collagen alpha-1(XVIII) chain | TRUE |
| 1975.9942 | GPRGDQGPVGRTGEVGAVGP*P | 20P(15.9949) | 21 | 820 | 840 | P08123 | COL1A2 | Collagen alpha-2(I) chain | TRUE |
| 1975.9942 | GPRGDQGPVGRTGEVGAVGPP* | 21P(15.9949) | 21 | 820 | 840 | P08123 | COL1A2 | Collagen alpha-2(I) chain | TRUE |
| 1976.0044 | LILAAAQMPAYQELVEEA | 8M(15.9949) | 18 | 48 | 65 | P37837 | TALDO1 | Transaldolase | TRUE |
| 1986.872 | P*QGSSGP*QGHMGPQGPPGPQG | 1P(15.9949),7P(15.9949) | 21 | 695 | 715 | Q9C0J8 | WDR33 | pre-mRNA 3' end processing protein WDR33 | TRUE |
| 1986.8871 | LMGENRTMTIHNGMFFS | 12N(0.9840) | 17 | 390 | 406 | P02675 | FGB | Fibrinogen beta chain | TRUE |
| 2010.9542 | VFTTEEVPAQQYLEIDE | | 17 | 618 | 634 | Q05707 | COL14A1 | Collagen alpha-1(XIV) chain | TRUE |
| 2010.9586 | GALGEP*GKQGSRGDP*GDAGPR | 15P(15.9949),6P(15.9949) | 21 | 478 | 498 | P12110 | COL6A2 | Collagen alpha-2(VI) chain | TRUE |
| 2079.98 | GPP*GKNGDDGEAGKPGRP*GER | 18P(15.9949),3P(15.9949) | 21 | 224 | 244 | P02452 | COL1A1 | Collagen alpha-1(I) chain | TRUE |
| 2079.98 | GKNGDDGEAGKPGRP*GERGP*P | 15P(15.9949),20P(15.9949) | 21 | 227 | 247 | P02452 | COL1A1 | Collagen alpha-1(I) chain | TRUE |
| 2079.98 | GP*PGKNGDDGEAGKPGRP*GER | 18P(15.9949),2P(15.9949) | 21 | 224 | 244 | P02452 | COL1A1 | Collagen alpha-1(I) chain | TRUE |
| 2079.9836 | LHADPDLGVLCPTGCQLQEA | | 20 | 96 | 115 | P02675 | FGB | Fibrinogen beta chain | TRUE |
| 2093.041 | GPAGEVGKP*GERGLHGEFGLP* | 21P(15.9949),9P(15.9949) | 21 | 562 | 582 | P08123 | COL1A2 | Collagen alpha-2(I) chain | TRUE |
| 2093.041 | PAGEVGKP*GERGLHGEFGLP*G | 20P(15.9949),8P(15.9949) | 21 | 563 | 583 | P08123 | COL1A2 | Collagen alpha-2(I) chain | TRUE |
| 2093.062 | GAPGGLAGDLVGEP*GAKGDRGLP* | 14P(15.9949),23P(15.9949) | 23 | 2313 | 2335 | Q02388 | COL7A1 | Collagen alpha-1(VII) chain | TRUE |
| 2095.0273 | GPAGKDGESGRP*GRP*GERGLP* | 12P(15.9949),15P(15.9949),21P(15.9949) | 21 | 225 | 245 | P02461 | COL3A1 | Collagen alpha-1(III) chain | TRUE |
| 2133.9765 | GPRGNRGERGSEGSP*GHP*GQP* | 15P(15.9949),18P(15.9949),21P(15.9949) | 21 | 1164 | 1184 | P02461 | COL3A1 | Collagen alpha-1(III) chain | TRUE |
| 2214.9968 | GAEGSP*GRDGSP*GAKGDRGETGPA | 12P(15.9949),6P(15.9949) | 24 | 1019 | 1042 | P02452 | COL1A1 | Collagen alpha-1(I) chain | TRUE |
| 2459.0553 | VIESNWNEIVDSFDDMNLSES | 16M(15.9949) | 21 | 22 | 42 | P60842 | EIF4A1 | Eukaryotic initiation factor 4A-I | TRUE |
