## Supplemental Table 3 for "Multimodal Imaging of the Cellular and Extracellular Microenvironment on the Same Formalin-Fixed Paraffin-Embedded Tissue Section"

**Supplemental Table 3 (1/3): Discriminating N-glycan features of each k-means segment by area under the receiver operating characteristic curve.** Segments are color coded as blue (B), green (G), purple (P), red (R),or yellow (Y) and indicated as N-glycan IMS performed without (-) or with (+) prior multiplexed immunofluorescence (MXIF) or picrosirius red staining (PSR). Values highlighted in green indicate top 10 discriminate features for each segment. Values highlighted in yellow indicate top 30 discriminate features for each segment. K-means segmentation analyses using Manhattan distance (k = 5) and receiver operating characteristic curves were performed in SCiLs v2026a Pro.

| m/z | MxIF- (B) | MxIF+ (B) | MxIF- (G) | MxIF+ (G) | MxIF- (P) | MxIF+ (P) | MxIF- (R) | MxIF+ (R) | MxIF- (Y) | MxIF+ (Y) | m/z | PSR- (B) | PSR+ (B) | PSR- (G) | PSR+ (G) | PSR- (P) | PSR+ (P) | PSR- (R) | PSR+ (R) | PSR- (Y) | PSR+ (Y) |
|---|---|---|---|---|---|---|---|---|---|---|---|---|---|---|---|---|---|---|---|---|---|
| 933.317 | 0.858 | 0.867 | 0.839 | 0.878 | 0.653 | 0.866 | 0.664 | 0.680 | 0.247 | 0.164 | 933.317 | 0.537 | 0.560 | 0.571 | 0.607 | 0.562 | 0.543 | 0.523 | 0.556 | 0.455 | 0.433 |
| 1095.370 | 0.801 | 0.813 | 0.878 | 0.861 | 0.734 | 0.920 | 0.653 | 0.675 | 0.229 | 0.156 | 1095.370 | 0.627 | 0.621 | 0.702 | 0.752 | 0.694 | 0.723 | 0.604 | 0.659 | 0.338 | 0.288 |
| 1136.396 | 0.864 | 0.884 | 0.882 | 0.921 | 0.681 | 0.838 | 0.762 | 0.704 | 0.177 | 0.153 | 1136.396 | 0.759 | 0.755 | 0.833 | 0.893 | 0.768 | 0.764 | 0.661 | 0.745 | 0.247 | 0.189 |
| 1257.423 | 0.882 | 0.847 | 0.935 | 0.902 | 0.825 | 0.973 | 0.712 | 0.720 | 0.143 | 0.099 | 1257.423 | 0.775 | 0.775 | 0.866 | 0.901 | 0.786 | 0.754 | 0.693 | 0.761 | 0.216 | 0.180 |
| 1282.454 | 0.944 | 0.956 | 0.960 | 0.868 | 0.818 | 0.928 | 0.652 | 0.641 | 0.164 | 0.156 | 1282.454 | 0.896 | 0.877 | 0.847 | 0.897 | 0.914 | 0.895 | 0.647 | 0.733 | 0.197 | 0.133 |
| 1298.449 | 0.763 | 0.822 | 0.824 | 0.970 | 0.604 | 0.799 | 0.867 | 0.798 | 0.173 | 0.109 | 1298.449 | 0.614 | 0.626 | 0.949 | 0.970 | 0.679 | 0.656 | 0.797 | 0.861 | 0.189 | 0.171 |
| 1339.476 | 0.913 | 0.913 | 0.932 | 0.777 | 0.791 | 0.901 | 0.579 | 0.585 | 0.222 | 0.217 | 1339.476 | 0.902 | 0.890 | 0.646 | 0.713 | 0.843 | 0.824 | 0.547 | 0.615 | 0.304 | 0.242 |
| 1419.476 | 0.785 | 0.784 | 0.907 | 0.895 | 0.763 | 0.917 | 0.724 | 0.735 | 0.180 | 0.119 | 1419.476 | 0.610 | 0.617 | 0.834 | 0.859 | 0.712 | 0.745 | 0.711 | 0.760 | 0.250 | 0.213 |
| 1428.512 | 0.761 | 0.800 | 0.989 | 0.640 | 0.966 | 0.995 | 0.500 | 0.541 | 0.222 | 0.238 | 1428.512 | 0.707 | 0.704 | 0.550 | 0.592 | 0.978 | 0.973 | 0.484 | 0.521 | 0.345 | 0.288 |
| 1444.507 | 0.830 | 0.863 | 0.964 | 0.968 | 0.828 | 0.939 | 0.832 | 0.802 | 0.082 | 0.050 | 1444.507 | 0.696 | 0.666 | 0.980 | 0.987 | 0.873 | 0.841 | 0.846 | 0.894 | 0.089 | 0.082 |
| 1460.502 | 0.778 | 0.811 | 0.918 | 0.951 | 0.731 | 0.911 | 0.801 | 0.763 | 0.149 | 0.094 | 1460.502 | 0.623 | 0.607 | 0.883 | 0.908 | 0.793 | 0.772 | 0.711 | 0.774 | 0.221 | 0.192 |
| 1485.534 | 0.941 | 0.939 | 0.974 | 0.871 | 0.873 | 0.957 | 0.691 | 0.649 | 0.122 | 0.142 | 1485.534 | 0.943 | 0.934 | 0.816 | 0.870 | 0.927 | 0.911 | 0.636 | 0.717 | 0.197 | 0.130 |
| 1501.529 | 0.772 | 0.819 | 0.856 | 0.944 | 0.668 | 0.769 | 0.818 | 0.769 | 0.172 | 0.143 | 1501.529 | 0.685 | 0.675 | 0.916 | 0.950 | 0.721 | 0.692 | 0.750 | 0.829 | 0.203 | 0.172 |
| 1524.464 | 0.562 | 0.625 | 0.589 | 0.942 | 0.588 | 0.571 | 0.466 | 0.804 | 0.464 | 0.218 | 1524.464 | 0.529 | 0.553 | 0.889 | 0.917 | 0.822 | 0.771 | 0.708 | 0.753 | 0.227 | 0.212 |
| 1540.459 | 0.490 | 0.579 | 0.482 | 0.930 | 0.462 | 0.573 | 0.844 | 0.775 | 0.331 | 0.243 | 1540.459 | 0.417 | 0.437 | 0.978 | 0.987 | 0.451 | 0.444 | 0.887 | 0.905 | 0.216 | 0.245 |
| 1542.555 | 0.940 | 0.928 | 0.972 | 0.570 | 0.892 | 0.974 | 0.400 | 0.486 | 0.273 | 0.272 | 1542.555 | 0.963 | 0.958 | 0.500 | 0.534 | 0.901 | 0.877 | 0.439 | 0.500 | 0.367 | 0.298 |
| 1546.446 | 0.733 | 0.741 | 0.912 | 0.811 | 0.803 | 0.909 | 0.501 | 0.623 | 0.295 | 0.208 | 1546.446 | 0.675 | 0.680 | 0.641 | 0.688 | 0.854 | 0.842 | 0.530 | 0.573 | 0.343 | 0.296 |
| 1581.528 | 0.855 | 0.834 | 0.789 | 0.905 | 0.636 | 0.833 | 0.670 | 0.736 | 0.257 | 0.142 | 1581.528 | 0.718 | 0.715 | 0.773 | 0.807 | 0.555 | 0.588 | 0.676 | 0.739 | 0.309 | 0.267 |
| 1589.545 | 0.719 | 0.794 | 0.967 | 0.735 | 0.893 | 0.979 | 0.588 | 0.593 | 0.211 | 0.202 | 1589.545 | 0.741 | 0.734 | 0.630 | 0.667 | 0.937 | 0.917 | 0.536 | 0.584 | 0.309 | 0.258 |
| 1590.565 | 0.855 | 0.860 | 0.992 | 0.527 | 0.970 | 0.996 | 0.386 | 0.486 | 0.265 | 0.277 | 1590.565 | 0.857 | 0.855 | 0.477 | 0.510 | 0.996 | 0.995 | 0.431 | 0.483 | 0.363 | 0.287 |
| 1606.560 | 0.808 | 0.809 | 0.982 | 0.799 | 0.952 | 0.991 | 0.561 | 0.614 | 0.187 | 0.176 | 1606.560 | 0.729 | 0.706 | 0.739 | 0.783 | 0.966 | 0.960 | 0.575 | 0.641 | 0.264 | 0.203 |
| 1611.527 | 0.723 | 0.792 | 0.496 | 0.937 | 0.413 | 0.618 | 0.813 | 0.785 | 0.322 | 0.193 | 1611.527 | 0.592 | 0.608 | 0.828 | 0.852 | 0.585 | 0.544 | 0.695 | 0.740 | 0.299 | 0.293 |
| 1622.555 | 0.648 | 0.700 | 0.767 | 0.974 | 0.614 | 0.804 | 0.830 | 0.814 | 0.219 | 0.112 | 1622.555 | 0.487 | 0.504 | 0.907 | 0.931 | 0.623 | 0.629 | 0.759 | 0.800 | 0.251 | 0.239 |
| 1631.592 | 0.830 | 0.831 | 0.988 | 0.639 | 0.965 | 0.994 | 0.469 | 0.547 | 0.227 | 0.231 | 1631.592 | 0.882 | 0.877 | 0.554 | 0.618 | 0.982 | 0.977 | 0.480 | 0.533 | 0.323 | 0.250 |
| 1647.587 | 0.797 | 0.837 | 0.988 | 0.942 | 0.889 | 0.963 | 0.858 | 0.797 | 0.049 | 0.050 | 1647.587 | 0.785 | 0.760 | 0.952 | 0.971 | 0.952 | 0.930 | 0.831 | 0.882 | 0.068 | 0.047 |
| 1663.581 | 0.710 | 0.805 | 0.461 | 0.976 | 0.353 | 0.516 | 0.918 | 0.859 | 0.293 | 0.177 | 1663.581 | 0.555 | 0.571 | 0.977 | 0.985 | 0.322 | 0.327 | 0.890 | 0.922 | 0.231 | 0.253 |
| 1688.613 | 0.940 | 0.933 | 0.983 | 0.621 | 0.919 | 0.975 | 0.423 | 0.498 | 0.249 | 0.258 | 1688.613 | 0.973 | 0.966 | 0.554 | 0.597 | 0.950 | 0.946 | 0.443 | 0.507 | 0.343 | 0.263 |
| 1692.504 | 0.812 | 0.792 | 0.952 | 0.532 | 0.884 | 0.943 | 0.393 | 0.489 | 0.305 | 0.303 | 1692.504 | 0.845 | 0.838 | 0.499 | 0.541 | 0.910 | 0.895 | 0.455 | 0.500 | 0.370 | 0.311 |
| 1704.608 | 0.868 | 0.879 | 0.940 | 0.776 | 0.828 | 0.934 | 0.547 | 0.595 | 0.233 | 0.202 | 1704.608 | 0.886 | 0.862 | 0.630 | 0.700 | 0.821 | 0.812 | 0.537 | 0.602 | 0.320 | 0.259 |
| 1735.603 | 0.663 | 0.721 | 0.963 | 0.790 | 0.861 | 0.973 | 0.650 | 0.636 | 0.199 | 0.180 | 1735.603 | 0.576 | 0.582 | 0.764 | 0.787 | 0.921 | 0.900 | 0.602 | 0.648 | 0.277 | 0.239 |
| 1745.635 | 0.800 | 0.707 | 0.480 | 0.347 | 0.455 | 0.315 | 0.400 | 0.409 | 0.520 | 0.616 | 1745.635 | 0.959 | 0.925 | 0.449 | 0.424 | 0.450 | 0.464 | 0.434 | 0.440 | 0.501 | 0.487 |
| 1749.525 | 0.770 | 0.775 | 0.953 | 0.607 | 0.842 | 0.947 | 0.483 | 0.546 | 0.279 | 0.259 | 1749.525 | 0.768 | 0.763 | 0.545 | 0.582 | 0.892 | 0.880 | 0.500 | 0.538 | 0.351 | 0.301 |
| 1751.597 | 0.637 | 0.704 | 0.807 | 0.823 | 0.713 | 0.879 | 0.630 | 0.641 | 0.288 | 0.211 | 1751.597 | 0.631 | 0.657 | 0.704 | 0.744 | 0.813 | 0.785 | 0.578 | 0.632 | 0.321 | 0.279 |
| 1752.618 | 0.709 | 0.744 | 0.978 | 0.624 | 0.945 | 0.991 | 0.486 | 0.538 | 0.248 | 0.250 | 1752.618 | 0.668 | 0.651 | 0.539 | 0.575 | 0.952 | 0.942 | 0.484 | 0.526 | 0.359 | 0.306 |
| 1757.585 | 0.769 | 0.811 | 0.637 | 0.907 | 0.477 | 0.694 | 0.748 | 0.737 | 0.307 | 0.195 | 1757.585 | 0.568 | 0.577 | 0.848 | 0.859 | 0.612 | 0.573 | 0.694 | 0.734 | 0.293 | 0.291 |
| 1768.613 | 0.747 | 0.766 | 0.978 | 0.664 | 0.961 | 0.991 | 0.463 | 0.546 | 0.248 | 0.238 | 1768.613 | 0.667 | 0.665 | 0.574 | 0.631 | 0.969 | 0.963 | 0.496 | 0.541 | 0.342 | 0.282 |
| 1773.579 | 0.663 | 0.720 | 0.465 | 0.945 | 0.414 | 0.606 | 0.796 | 0.781 | 0.346 | 0.208 | 1773.579 | 0.584 | 0.608 | 0.838 | 0.860 | 0.640 | 0.600 | 0.669 | 0.717 | 0.301 | 0.287 |
| 1777.649 | 0.615 | 0.628 | 0.970 | 0.711 | 0.891 | 0.967 | 0.634 | 0.606 | 0.205 | 0.221 | 1777.649 | 0.637 | 0.605 | 0.682 | 0.720 | 0.863 | 0.877 | 0.581 | 0.605 | 0.308 | 0.274 |
| 1792.624 | 0.646 | 0.698 | 0.917 | 0.975 | 0.757 | 0.890 | 0.849 | 0.811 | 0.138 | 0.082 | 1792.624 | 0.695 | 0.690 | 0.967 | 0.975 | 0.853 | 0.824 | 0.797 | 0.838 | 0.128 | 0.118 |
| 1793.644 | 0.851 | 0.842 | 0.992 | 0.647 | 0.968 | 0.995 | 0.446 | 0.529 | 0.234 | 0.240 | 1793.644 | 0.901 | 0.894 | 0.614 | 0.654 | 0.994 | 0.994 | 0.449 | 0.522 | 0.331 | 0.244 |
| 1809.639 | 0.445 | 0.535 | 0.681 | 0.996 | 0.507 | 0.704 | 0.968 | 0.931 | 0.229 | 0.092 | 1809.639 | 0.352 | 0.386 | 0.998 | 0.999 | 0.514 | 0.504 | 0.963 | 0.957 | 0.155 | 0.202 |
| 1814.606 | 0.699 | 0.768 | 0.583 | 0.889 | 0.518 | 0.619 | 0.693 | 0.708 | 0.343 | 0.249 | 1814.606 | 0.570 | 0.595 | 0.864 | 0.900 | 0.579 | 0.550 | 0.697 | 0.743 | 0.298 | 0.287 |
| 1815.559 | 0.535 | 0.608 | 0.550 | 0.937 | 0.496 | 0.422 | 0.593 | 0.802 | 0.438 | 0.277 | 1815.559 | 0.487 | 0.526 | 0.915 | 0.943 | 0.560 | 0.533 | 0.750 | 0.790 | 0.274 | 0.271 |
| 1831.554 | 0.554 | 0.645 | 0.392 | 0.921 | 0.361 | 0.445 | 0.849 | 0.760 | 0.365 | 0.292 | 1831.554 | 0.481 | 0.485 | 0.984 | 0.992 | 0.238 | 0.251 | 0.898 | 0.917 | 0.258 | 0.294 |
| 1832.575 | 0.542 | 0.589 | 0.570 | 0.836 | 0.524 | 0.657 | 0.812 | 0.694 | 0.305 | 0.272 | 1832.575 | 0.462 | 0.472 | 0.854 | 0.825 | 0.597 | 0.588 | 0.798 | 0.769 | 0.243 | 0.286 |
| 1834.671 | 0.845 | 0.836 | 0.988 | 0.514 | 0.951 | 0.990 | 0.406 | 0.487 | 0.263 | 0.283 | 1834.671 | 0.922 | 0.917 | 0.470 | 0.478 | 0.973 | 0.971 | 0.441 | 0.477 | 0.355 | 0.293 |
| 1837.541 | 0.587 | 0.599 | 0.556 | 0.990 | 0.497 | 0.522 | 0.687 | 0.900 | 0.377 | 0.173 | 1837.541 | 0.438 | 0.502 | 0.964 | 0.983 | 0.584 | 0.545 | 0.864 | 0.874 | 0.193 | 0.219 |
| 1850.666 | 0.914 | 0.909 | 0.989 | 0.795 | 0.940 | 0.985 | 0.587 | 0.641 | 0.157 | 0.148 | 1850.666 | 0.947 | 0.939 | 0.743 | 0.798 | 0.966 | 0.955 | 0.585 | 0.665 | 0.228 | 0.153 |
| 1853.536 | 0.522 | 0.597 | 0.420 | 0.951 | 0.366 | 0.474 | 0.889 | 0.818 | 0.344 | 0.248 | 1853.536 | 0.430 | 0.446 | 0.992 | 0.995 | 0.300 | 0.302 | 0.925 | 0.933 | 0.229 | 0.274 |
| 1854.557 | 0.521 | 0.562 | 0.663 | 0.889 | 0.576 | 0.738 | 0.847 | 0.746 | 0.259 | 0.207 | 1854.557 | 0.450 | 0.452 | 0.924 | 0.923 | 0.528 | 0.528 | 0.845 | 0.821 | 0.224 | 0.269 |
| 1859.523 | 0.522 | 0.557 | 0.560 | 0.953 | 0.497 | 0.539 | 0.729 | 0.799 | 0.365 | 0.240 | 1859.523 | 0.466 | 0.505 | 0.873 | 0.901 | 0.625 | 0.574 | 0.703 | 0.738 | 0.295 | 0.296 |
| 1866.661 | 0.705 | 0.770 | 0.803 | 0.868 | 0.682 | 0.798 | 0.670 | 0.679 | 0.266 | 0.203 | 1866.661 | 0.634 | 0.652 | 0.783 | 0.830 | 0.699 | 0.689 | 0.651 | 0.712 | 0.296 | 0.256 |
| 1889.596 | 0.574 | 0.624 | 0.631 | 0.884 | 0.591 | 0.711 | 0.604 | 0.697 | 0.380 | 0.241 | 1889.596 | 0.587 | 0.567 | 0.741 | 0.772 | 0.804 | 0.770 | 0.604 | 0.648 | 0.309 | 0.286 |
| 1891.692 | 0.913 | 0.883 | 0.550 | 0.511 | 0.401 | 0.557 | 0.431 | 0.490 | 0.492 | 0.436 | 1891.692 | 0.991 | 0.983 | 0.475 | 0.494 | 0.505 | 0.494 | 0.455 | 0.492 | 0.465 | 0.430 |
| 1895.583 | 0.815 | 0.788 | 0.982 | 0.472 | 0.934 | 0.980 | 0.366 | 0.454 | 0.298 | 0.318 | 1895.583 | 0.886 | 0.881 | 0.430 | 0.477 | 0.969 | 0.968 | 0.417 | 0.455 | 0.381 | 0.312 |
| 1905.634 | 0.779 | 0.768 | 0.932 | 0.804 | 0.848 | 0.936 | 0.574 | 0.650 | 0.229 | 0.177 | 1905.634 | 0.671 | 0.647 | 0.671 | 0.706 | 0.769 | 0.821 | 0.602 | 0.666 | 0.316 | 0.253 |
| 1907.687 | 0.657 | 0.459 | 0.795 | 0.434 | 0.738 | 0.436 | 0.468 | 0.459 | 0.365 | 0.562 | 1907.687 | 0.773 | 0.710 | 0.452 | 0.416 | 0.481 | 0.578 | 0.462 | 0.446 | 0.498 | 0.480 |
| 1911.578 | 0.548 | 0.574 | 0.920 | 0.804 | 0.813 | 0.936 | 0.669 | 0.697 | 0.232 | 0.172 | 1911.578 | 0.562 | 0.560 | 0.734 | 0.752 | 0.858 | 0.861 | 0.632 | 0.656 | 0.280 | 0.254 |
| 1913.650 | 0.529 | 0.712 | 0.582 | 0.834 | 0.573 | 0.557 | 0.503 | 0.663 | 0.456 | 0.314 | 1913.650 | 0.572 | 0.616 | 0.828 | 0.869 | 0.557 | 0.531 | 0.667 | 0.728 | 0.328 | 0.301 |
| 1914.671 | 0.819 | 0.827 | 0.879 | 0.801 | 0.751 | 0.872 | 0.592 | 0.620 | 0.253 | 0.214 | 1914.671 | 0.741 | 0.731 | 0.747 | 0.805 | 0.755 | 0.739 | 0.585 | 0.643 | 0.313 | 0.270 |
| 1938.682 | 0.679 | 0.721 | 0.980 | 0.837 | 0.906 | 0.983 | 0.700 | 0.683 | 0.151 | 0.142 | 1938.682 | 0.709 | 0.700 | 0.812 | 0.838 | 0.964 | 0.955 | 0.640 | 0.692 | 0.217 | 0.170 |
| 1939.702 | 0.729 | 0.766 | 0.988 | 0.729 | 0.961 | 0.995 | 0.611 | 0.612 | 0.170 | 0.188 | 1939.702 | 0.763 | 0.745 | 0.648 | 0.687 | 0.989 | 0.986 | 0.543 | 0.587 | 0.285 | 0.230 |
| 1948.714 | 0.889 | 0.843 | 0.528 | 0.524 | 0.457 | 0.456 | 0.437 | 0.492 | 0.476 | 0.475 | 1948.714 | 0.970 | 0.963 | 0.484 | 0.469 | 0.456 | 0.489 | 0.456 | 0.479 | 0.479 | 0.446 |
| 1954.677 | 0.699 | 0.784 | 0.615 | 0.943 | 0.502 | 0.679 | 0.851 | 0.778 | 0.258 | 0.175 | 1954.677 | 0.636 | 0.634 | 0.952 | 0.971 | 0.588 | 0.563 | 0.813 | 0.862 | 0.200 | 0.200 |
| 1955.697 | 0.735 | 0.793 | 0.987 | 0.829 | 0.973 | 0.997 | 0.706 | 0.696 | 0.113 | 0.120 | 1955.697 | 0.700 | 0.677 | 0.862 | 0.872 | 0.993 | 0.991 | 0.700 | 0.755 | 0.165 | 0.123 |
| 1960.664 | 0.726 | 0.785 | 0.756 | 0.959 | 0.557 | 0.767 | 0.858 | 0.803 | 0.211 | 0.124 | 1960.664 | 0.649 | 0.626 | 0.912 | 0.930 | 0.751 | 0.680 | 0.755 | 0.804 | 0.197 | 0.200 |
| 1976.659 | 0.723 | 0.811 | 0.456 | 0.976 | 0.361 | 0.504 | 0.916 | 0.862 | 0.290 | 0.178 | 1976.659 | 0.595 | 0.611 | 0.967 | 0.977 | 0.338 | 0.351 | 0.868 | 0.901 | 0.237 | 0.252 |
| 1980.729 | 0.574 | 0.494 | 0.966 | 0.350 | 0.907 | 0.935 | 0.393 | 0.400 | 0.337 | 0.418 | 1980.729 | 0.697 | 0.659 | 0.434 | 0.390 | 0.875 | 0.882 | 0.446 | 0.462 | 0.413 | 0.381 |
| 1987.644 | 0.517 | 0.558 | 0.948 | 0.676 | 0.893 | 0.982 | 0.530 | 0.562 | 0.281 | 0.257 | 1987.644 | 0.580 | 0.588 | 0.579 | 0.584 | 0.933 | 0.895 | 0.505 | 0.525 | 0.358 | 0.331 |
| 1995.703 | 0.776 | 0.808 | 0.916 | 0.764 | 0.794 | 0.918 | 0.568 | 0.606 | 0.254 | 0.212 | 1995.703 | 0.845 | 0.843 | 0.710 | 0.754 | 0.861 | 0.853 | 0.555 | 0.619 | 0.294 | 0.233 |
| 1996.724 | 0.888 | 0.863 | 0.991 | 0.524 | 0.955 | 0.993 | 0.353 | 0.481 | 0.283 | 0.281 | 1996.724 | 0.954 | 0.950 | 0.478 | 0.526 | 0.983 | 0.980 | 0.401 | 0.461 | 0.373 | 0.288 |
| 2012.719 | 0.708 | 0.774 | 0.952 | 0.926 | 0.860 | 0.935 | 0.767 | 0.774 | 0.129 | 0.085 | 2012.719 | 0.672 | 0.661 | 0.903 | 0.934 | 0.870 | 0.831 | 0.751 | 0.819 | 0.164 | 0.135 |
| 2026.617 | 0.509 | 0.531 | 0.600 | 0.927 | 0.537 | 0.664 | 0.782 | 0.790 | 0.319 | 0.206 | 2026.617 | 0.508 | 0.513 | 0.778 | 0.796 | 0.698 | 0.681 | 0.607 | 0.638 | 0.342 | 0.328 |
| 2028.714 | 0.720 | 0.806 | 0.536 | 0.952 | 0.448 | 0.579 | 0.830 | 0.801 | 0.296 | 0.193 | 2028.714 | 0.570 | 0.573 | 0.933 | 0.949 | 0.403 | 0.410 | 0.818 | 0.861 | 0.259 | 0.264 |
| 2037.750 | 0.875 | 0.859 | 0.945 | 0.505 | 0.892 | 0.960 | 0.407 | 0.490 | 0.285 | 0.290 | 2037.750 | 0.940 | 0.932 | 0.475 | 0.510 | 0.837 | 0.835 | 0.453 | 0.487 | 0.382 | 0.326 |
| 2048.599 | 0.521 | 0.570 | 0.525 | 0.977 | 0.442 | 0.616 | 0.855 | 0.836 | 0.321 | 0.184 | 2048.599 | 0.502 | 0.490 | 0.902 | 0.910 | 0.533 | 0.570 | 0.742 | 0.763 | 0.286 | 0.284 |
| 2053.745 | 0.949 | 0.928 | 0.733 | 0.505 | 0.556 | 0.664 | 0.405 | 0.486 | 0.420 | 0.394 | 2053.745 | 0.996 | 0.994 | 0.460 | 0.487 | 0.641 | 0.613 | 0.438 | 0.482 | 0.439 | 0.395 |
| 2057.636 | 0.750 | 0.766 | 0.977 | 0.508 | 0.952 | 0.985 | 0.403 | 0.480 | 0.283 | 0.299 | 2057.636 | 0.767 | 0.758 | 0.485 | 0.500 | 0.979 | 0.973 | 0.456 | 0.491 | 0.362 | 0.307 |
| 2084.740 | 0.541 | 0.577 | 0.987 | 0.746 | 0.953 | 0.993 | 0.636 | 0.626 | 0.193 | 0.202 | 2084.740 | 0.581 | 0.580 | 0.751 | 0.776 | 0.985 | 0.979 | 0.601 | 0.637 | 0.261 | 0.220 |
| 2094.772 | 0.899 | 0.870 | 0.451 | 0.501 | 0.399 | 0.354 | 0.454 | 0.501 | 0.496 | 0.507 | 2094.772 | 0.984 | 0.972 | 0.507 | 0.517 | 0.422 | 0.442 | 0.470 | 0.496 | 0.475 | 0.445 |
| 2098.663 | 0.872 | 0.836 | 0.934 | 0.552 | 0.818 | 0.907 | 0.378 | 0.484 | 0.328 | 0.312 | 2098.663 | 0.947 | 0.945 | 0.522 | 0.559 | 0.876 | 0.858 | 0.443 | 0.495 | 0.371 | 0.307 |
| 2100.735 | 0.422 | 0.480 | 0.599 | 0.980 | 0.446 | 0.664 | 0.939 | 0.880 | 0.282 | 0.149 | 2100.735 | 0.426 | 0.447 | 0.996 | 0.997 | 0.542 | 0.532 | 0.935 | 0.922 | 0.156 | 0.203 |
| 2101.755 | 0.560 | 0.640 | 0.993 | 0.868 | 0.972 | 0.997 | 0.792 | 0.753 | 0.097 | 0.098 | 2101.755 | 0.571 | 0.571 | 0.908 | 0.917 | 0.997 | 0.996 | 0.796 | 0.797 | 0.113 | 0.109 |
| 2106.722 | 0.525 | 0.570 | 0.950 | 0.822 | 0.883 | 0.986 | 0.683 | 0.659 | 0.200 | 0.176 | 2106.722 | 0.513 | 0.506 | 0.761 | 0.788 | 0.974 | 0.939 | 0.613 | 0.629 | 0.264 | 0.249 |
| 2122.717 | 0.420 | 0.486 | 0.409 | 0.992 | 0.285 | 0.488 | 0.973 | 0.928 | 0.347 | 0.181 | 2122.717 | 0.357 | 0.395 | 0.999 | 0.999 | 0.320 | 0.323 | 0.958 | 0.945 | 0.211 | 0.267 |
| 2123.670 | 0.544 | 0.605 | 0.624 | 0.875 | 0.586 | 0.298 | 0.427 | 0.781 | 0.484 | 0.343 | 2123.670 | 0.423 | 0.541 | 0.914 | 0.955 | 0.414 | 0.369 | 0.780 | 0.786 | 0.303 | 0.325 |
| 2126.787 | 0.491 | 0.511 | 0.966 | 0.372 | 0.904 | 0.871 | 0.424 | 0.389 | 0.337 | 0.445 | 2126.787 | 0.554 | 0.520 | 0.422 | 0.392 | 0.801 | 0.835 | 0.442 | 0.440 | 0.457 | 0.431 |
| 2130.652 | 0.576 | 0.577 | 0.551 | 0.906 | 0.518 | 0.673 | 0.702 | 0.725 | 0.364 | 0.241 | 2130.652 | 0.528 | 0.520 | 0.807 | 0.837 | 0.661 | 0.634 | 0.651 | 0.676 | 0.318 | 0.315 |
| 2141.761 | 0.745 | 0.779 | 0.979 | 0.723 | 0.925 | 0.986 | 0.568 | 0.599 | 0.204 | 0.199 | 2141.761 | 0.796 | 0.779 | 0.689 | 0.723 | 0.963 | 0.959 | 0.550 | 0.600 | 0.278 | 0.222 |
| 2142.782 | 0.797 | 0.808 | 0.982 | 0.569 | 0.973 | 0.994 | 0.421 | 0.521 | 0.257 | 0.257 | 2142.782 | 0.837 | 0.825 | 0.514 | 0.562 | 0.987 | 0.987 | 0.446 | 0.501 | 0.355 | 0.279 |
| 2145.652 | 0.579 | 0.626 | 0.529 | 0.843 | 0.471 | 0.620 | 0.825 | 0.712 | 0.316 | 0.269 | 2145.652 | 0.546 | 0.537 | 0.955 | 0.964 | 0.411 | 0.414 | 0.847 | 0.860 | 0.238 | 0.267 |
| 2149.697 | 0.547 | 0.528 | 0.576 | 0.672 | 0.639 | 0.979 | 0.586 | 0.553 | 0.386 | 0.268 | 2149.697 | 0.539 | 0.521 | 0.589 | 0.619 | 0.948 | 0.847 | 0.511 | 0.537 | 0.354 | 0.347 |
| 2155.660 | 0.812 | 0.809 | 0.627 | 0.656 | 0.515 | 0.607 | 0.537 | 0.555 | 0.402 | 0.370 | 2155.660 | 0.880 | 0.881 | 0.593 | 0.609 | 0.600 | 0.583 | 0.513 | 0.547 | 0.402 | 0.374 |
| 2155.684 | 0.825 | 0.803 | 0.658 | 0.609 | 0.526 | 0.630 | 0.512 | 0.543 | 0.405 | 0.375 | 2155.684 | 0.897 | 0.889 | 0.535 | 0.572 | 0.616 | 0.595 | 0.502 | 0.533 | 0.409 | 0.380 |
| 2156.680 | 0.806 | 0.778 | 0.666 | 0.494 | 0.546 | 0.587 | 0.460 | 0.491 | 0.428 | 0.439 | 2156.680 | 0.891 | 0.881 | 0.537 | 0.520 | 0.620 | 0.605 | 0.479 | 0.508 | 0.423 | 0.398 |
| 2157.756 | 0.692 | 0.766 | 0.783 | 0.926 | 0.668 | 0.791 | 0.711 | 0.750 | 0.253 | 0.155 | 2157.756 | 0.700 | 0.689 | 0.889 | 0.927 | 0.706 | 0.679 | 0.728 | 0.785 | 0.222 | 0.202 |
| 2158.777 | 0.852 | 0.858 | 0.994 | 0.689 | 0.964 | 0.995 | 0.511 | 0.585 | 0.200 | 0.198 | 2158.777 | 0.863 | 0.855 | 0.669 | 0.712 | 0.992 | 0.987 | 0.510 | 0.586 | 0.290 | 0.209 |
| 2163.743 | 0.747 | 0.800 | 0.919 | 0.774 | 0.800 | 0.941 | 0.628 | 0.640 | 0.224 | 0.181 | 2163.743 | 0.724 | 0.727 | 0.672 | 0.709 | 0.871 | 0.847 | 0.587 | 0.629 | 0.291 | 0.252 |
| 2167.634 | 0.554 | 0.598 | 0.449 | 0.861 | 0.419 | 0.526 | 0.836 | 0.740 | 0.344 | 0.287 | 2167.634 | 0.519 | 0.513 | 0.920 | 0.907 | 0.407 | 0.437 | 0.835 | 0.820 | 0.255 | 0.292 |
| 2172.632 | 0.601 | 0.650 | 0.458 | 0.964 | 0.411 | 0.494 | 0.850 | 0.818 | 0.329 | 0.233 | 2172.632 | 0.638 | 0.645 | 0.892 | 0.907 | 0.571 | 0.555 | 0.719 | 0.747 | 0.274 | 0.274 |
| 2172.675 | 0.562 | 0.527 | 0.684 | 0.404 | 0.600 | 0.836 | 0.715 | 0.491 | 0.312 | 0.387 | 2172.675 | 0.568 | 0.547 | 0.557 | 0.542 | 0.777 | 0.748 | 0.548 | 0.559 | 0.377 | 0.372 |
| 2174.772 | 0.613 | 0.731 | 0.714 | 0.990 | 0.605 | 0.747 | 0.911 | 0.889 | 0.191 | 0.080 | 2174.772 | 0.425 | 0.458 | 0.982 | 0.986 | 0.498 | 0.499 | 0.916 | 0.937 | 0.183 | 0.205 |
| 2183.808 | 0.746 | 0.755 | 0.968 | 0.492 | 0.962 | 0.984 | 0.412 | 0.492 | 0.277 | 0.294 | 2183.808 | 0.788 | 0.761 | 0.439 | 0.473 | 0.948 | 0.944 | 0.440 | 0.484 | 0.384 | 0.323 |
| 2190.724 | 0.586 | 0.595 | 0.986 | 0.661 | 0.934 | 0.988 | 0.671 | 0.611 | 0.173 | 0.219 | 2190.724 | 0.481 | 0.485 | 0.647 | 0.682 | 0.984 | 0.969 | 0.580 | 0.614 | 0.300 | 0.261 |
| 2199.803 | 0.922 | 0.882 | 0.808 | 0.469 | 0.629 | 0.752 | 0.398 | 0.468 | 0.393 | 0.382 | 2199.803 | 0.995 | 0.993 | 0.458 | 0.493 | 0.708 | 0.681 | 0.424 | 0.471 | 0.430 | 0.379 |
| 2202.674 | 0.598 | 0.614 | 0.713 | 0.924 | 0.610 | 0.717 | 0.680 | 0.755 | 0.317 | 0.199 | 2202.674 | 0.601 | 0.617 | 0.830 | 0.853 | 0.708 | 0.687 | 0.672 | 0.703 | 0.279 | 0.265 |
| 2203.694 | 0.704 | 0.720 | 0.982 | 0.756 | 0.966 | 0.990 | 0.545 | 0.614 | 0.209 | 0.192 | 2203.694 | 0.748 | 0.736 | 0.695 | 0.721 | 0.994 | 0.991 | 0.544 | 0.595 | 0.280 | 0.221 |
| 2215.798 | 0.943 | 0.954 | 0.891 | 0.608 | 0.769 | 0.884 | 0.435 | 0.533 | 0.308 | 0.268 | 2215.798 | 0.956 | 0.944 | 0.511 | 0.544 | 0.769 | 0.757 | 0.465 | 0.533 | 0.386 | 0.320 |
| 2244.721 | 0.684 | 0.717 | 0.959 | 0.616 | 0.936 | 0.978 | 0.431 | 0.531 | 0.288 | 0.263 | 2244.721 | 0.701 | 0.705 | 0.583 | 0.601 | 0.914 | 0.906 | 0.487 | 0.530 | 0.358 | 0.304 |
| 2245.772 | 0.674 | 0.714 | 0.880 | 0.810 | 0.817 | 0.872 | 0.704 | 0.654 | 0.195 | 0.205 | 2245.772 | 0.650 | 0.656 | 0.869 | 0.898 | 0.844 | 0.825 | 0.694 | 0.743 | 0.216 | 0.186 |
| 2246.793 | 0.654 | 0.691 | 0.873 | 0.778 | 0.775 | 0.914 | 0.724 | 0.640 | 0.203 | 0.204 | 2246.793 | 0.606 | 0.609 | 0.811 | 0.831 | 0.848 | 0.832 | 0.679 | 0.702 | 0.237 | 0.221 |
| 2256.825 | 0.918 | 0.883 | 0.541 | 0.528 | 0.456 | 0.501 | 0.432 | 0.501 | 0.472 | 0.448 | 2256.825 | 0.985 | 0.982 | 0.469 | 0.483 | 0.466 | 0.488 | 0.451 | 0.477 | 0.479 | 0.442 |
| 2260.715 | 0.808 | 0.806 | 0.962 | 0.478 | 0.876 | 0.955 | 0.383 | 0.474 | 0.313 | 0.311 | 2260.715 | 0.875 | 0.859 | 0.446 | 0.447 | 0.929 | 0.915 | 0.424 | 0.467 | 0.387 | 0.325 |
| 2267.754 | 0.712 | 0.784 | 0.463 | 0.928 | 0.392 | 0.564 | 0.847 | 0.783 | 0.318 | 0.215 | 2267.754 | 0.651 | 0.655 | 0.940 | 0.954 | 0.423 | 0.430 | 0.800 | 0.842 | 0.253 | 0.254 |

**Supplemental Table 3 (2/3): Discriminating N-glycan features of each k-means segment by area under the receiver operating characteristic curve.** Segments are color coded as blue (B), green (G), purple (P), red (R),or yellow (Y) and indicated as N-glycan IMS performed without (-) or with (+) prior multiplexed immunofluorescence (MXIF) or picrosirius red staining (PSR). Values highlighted in green indicate top 10 discriminate features for each segment. Values highlighted in yellow indicate top 30 discriminate features for each segment. K-means segmentation analyses using Manhattan distance (k = 5) and receiver operating characteristic curves were performed in SCiLs v2026a Pro.

| m/z | MxIF- (B) | MxIF+ (B) | MxIF- (G) | MxIF+ (G) | MxIF- (P) | MxIF+ (P) | MxIF- (R) | MxIF+ (R) | MxIF- (Y) | MxIF+ (Y) | m/z | PSR- (B) | PSR+ (B) | PSR- (G) | PSR+ (G) | PSR- (P) | PSR+ (P) | PSR- (R) | PSR+ (R) | PSR- (Y) | PSR+ (Y) |
|---|---|---|---|---|---|---|---|---|---|---|---|---|---|---|---|---|---|---|---|---|---|
| **2268.775** | 0.693 | 0.785 | 0.823 | 0.915 | 0.702 | 0.854 | 0.815 | 0.754 | 0.179 | 0.128 | **2268.775** | 0.614 | 0.622 | 0.913 | 0.912 | 0.729 | 0.709 | 0.772 | 0.807 | 0.196 | 0.191 |
| **2287.819** | 0.658 | 0.704 | 0.988 | 0.616 | 0.964 | 0.995 | 0.495 | 0.545 | 0.244 | 0.250 | **2287.819** | 0.689 | 0.674 | 0.586 | 0.632 | 0.992 | 0.989 | 0.499 | 0.535 | 0.330 | 0.276 |
| **2288.840** | 0.648 | 0.663 | 0.986 | 0.566 | 0.963 | 0.992 | 0.473 | 0.518 | 0.258 | 0.279 | **2288.840** | 0.651 | 0.638 | 0.529 | 0.552 | 0.958 | 0.952 | 0.482 | 0.506 | 0.362 | 0.318 |
| **2289.736** | 0.734 | 0.806 | 0.496 | 0.975 | 0.388 | 0.524 | 0.913 | 0.867 | 0.274 | 0.169 | **2289.736** | 0.658 | 0.665 | 0.963 | 0.968 | 0.397 | 0.408 | 0.858 | 0.891 | 0.219 | 0.230 |
| **2301.718** | 0.882 | 0.832 | 0.681 | 0.537 | 0.521 | 0.606 | 0.440 | 0.502 | 0.432 | 0.414 | **2301.718** | 0.983 | 0.977 | 0.511 | 0.552 | 0.629 | 0.614 | 0.447 | 0.487 | 0.433 | 0.388 |
| **2301.742** | 0.883 | 0.831 | 0.694 | 0.543 | 0.519 | 0.603 | 0.435 | 0.507 | 0.434 | 0.411 | **2301.742** | 0.986 | 0.979 | 0.521 | 0.547 | 0.628 | 0.610 | 0.445 | 0.488 | 0.433 | 0.389 |
| **2302.695** | 0.497 | 0.822 | 0.597 | 0.532 | 0.591 | 0.573 | 0.520 | 0.498 | 0.444 | 0.430 | **2302.695** | 0.871 | 0.920 | 0.534 | 0.559 | 0.679 | 0.624 | 0.460 | 0.504 | 0.423 | 0.383 |
| **2303.814** | 0.579 | 0.617 | 0.914 | 0.893 | 0.812 | 0.927 | 0.689 | 0.717 | 0.217 | 0.148 | **2303.814** | 0.613 | 0.630 | 0.893 | 0.927 | 0.852 | 0.839 | 0.701 | 0.747 | 0.211 | 0.179 |
| **2304.835** | 0.836 | 0.834 | 0.993 | 0.685 | 0.964 | 0.995 | 0.517 | 0.591 | 0.200 | 0.197 | **2304.835** | 0.871 | 0.865 | 0.713 | 0.749 | 0.992 | 0.990 | 0.527 | 0.589 | 0.273 | 0.201 |
| **2310.755** | 0.592 | 0.646 | 0.887 | 0.570 | 0.851 | 0.989 | 0.468 | 0.511 | 0.324 | 0.286 | **2310.755** | 0.573 | 0.592 | 0.534 | 0.553 | 0.963 | 0.905 | 0.483 | 0.510 | 0.370 | 0.339 |
| **2320.829** | 0.725 | 0.781 | 0.968 | 0.746 | 0.956 | 0.988 | 0.581 | 0.626 | 0.191 | 0.178 | **2320.829** | 0.651 | 0.653 | 0.710 | 0.738 | 0.956 | 0.946 | 0.585 | 0.643 | 0.275 | 0.220 |
| **2325.796** | 0.631 | 0.703 | 0.782 | 0.927 | 0.674 | 0.841 | 0.759 | 0.756 | 0.236 | 0.141 | **2325.796** | 0.568 | 0.549 | 0.846 | 0.874 | 0.703 | 0.698 | 0.692 | 0.745 | 0.269 | 0.246 |
| **2326.749** | 0.651 | 0.725 | 0.590 | 0.865 | 0.549 | 0.595 | 0.512 | 0.703 | 0.437 | 0.269 | **2326.749** | 0.577 | 0.600 | 0.810 | 0.866 | 0.662 | 0.607 | 0.638 | 0.705 | 0.319 | 0.292 |
| **2333.732** | 0.480 | 0.512 | 0.519 | 0.969 | 0.444 | 0.596 | 0.862 | 0.828 | 0.325 | 0.205 | **2333.732** | 0.476 | 0.468 | 0.949 | 0.956 | 0.558 | 0.551 | 0.790 | 0.796 | 0.246 | 0.270 |
| **2336.781** | 0.460 | 0.518 | 0.766 | 0.692 | 0.636 | 0.852 | 0.817 | 0.684 | 0.251 | 0.229 | **2336.781** | 0.479 | 0.469 | 0.705 | 0.687 | 0.770 | 0.729 | 0.638 | 0.645 | 0.315 | 0.326 |
| **2339.652** | 0.550 | 0.576 | 0.570 | 0.890 | 0.523 | 0.513 | 0.618 | 0.764 | 0.409 | 0.276 | **2339.652** | 0.562 | 0.566 | 0.779 | 0.784 | 0.610 | 0.591 | 0.617 | 0.629 | 0.353 | 0.355 |
| **2339.695** | 0.549 | 0.560 | 0.543 | 0.915 | 0.471 | 0.551 | 0.804 | 0.797 | 0.331 | 0.241 | **2339.695** | 0.553 | 0.550 | 0.822 | 0.809 | 0.562 | 0.570 | 0.640 | 0.651 | 0.347 | 0.349 |
| **2340.672** | 0.589 | 0.585 | 0.569 | 0.912 | 0.495 | 0.590 | 0.758 | 0.789 | 0.337 | 0.228 | **2340.672** | 0.708 | 0.716 | 0.785 | 0.805 | 0.657 | 0.627 | 0.596 | 0.613 | 0.334 | 0.327 |
| **2340.715** | 0.573 | 0.562 | 0.637 | 0.858 | 0.534 | 0.655 | 0.786 | 0.758 | 0.302 | 0.233 | **2340.715** | 0.597 | 0.594 | 0.733 | 0.727 | 0.622 | 0.637 | 0.593 | 0.600 | 0.366 | 0.358 |
| **2341.791** | 0.710 | 0.786 | 0.502 | 0.951 | 0.435 | 0.569 | 0.817 | 0.798 | 0.313 | 0.202 | **2341.791** | 0.600 | 0.617 | 0.879 | 0.899 | 0.442 | 0.457 | 0.749 | 0.796 | 0.295 | 0.284 |
| **2344.841** | 0.800 | 0.745 | 0.902 | 0.464 | 0.786 | 0.920 | 0.553 | 0.503 | 0.262 | 0.314 | **2344.841** | 0.849 | 0.836 | 0.494 | 0.497 | 0.842 | 0.838 | 0.477 | 0.504 | 0.374 | 0.332 |
| **2345.861** | 0.893 | 0.865 | 0.918 | 0.481 | 0.792 | 0.901 | 0.435 | 0.480 | 0.305 | 0.320 | **2345.861** | 0.972 | 0.961 | 0.443 | 0.481 | 0.824 | 0.811 | 0.424 | 0.465 | 0.403 | 0.345 |
| **2352.776** | 0.547 | 0.628 | 0.985 | 0.646 | 0.964 | 0.996 | 0.482 | 0.547 | 0.272 | 0.255 | **2352.776** | 0.510 | 0.507 | 0.556 | 0.587 | 0.993 | 0.977 | 0.486 | 0.509 | 0.366 | 0.326 |
| **2361.634** | 0.589 | 0.635 | 0.486 | 0.957 | 0.446 | 0.453 | 0.700 | 0.823 | 0.397 | 0.247 | **2361.634** | 0.553 | 0.592 | 0.922 | 0.941 | 0.504 | 0.489 | 0.751 | 0.781 | 0.279 | 0.281 |
| **2361.677** | 0.583 | 0.618 | 0.452 | 0.968 | 0.382 | 0.511 | 0.885 | 0.837 | 0.325 | 0.217 | **2361.677** | 0.568 | 0.571 | 0.935 | 0.939 | 0.506 | 0.499 | 0.773 | 0.790 | 0.260 | 0.276 |
| **2361.856** | 0.941 | 0.923 | 0.943 | 0.501 | 0.838 | 0.925 | 0.365 | 0.475 | 0.315 | 0.305 | **2361.856** | 0.985 | 0.984 | 0.429 | 0.471 | 0.880 | 0.863 | 0.400 | 0.453 | 0.403 | 0.332 |
| **2362.654** | 0.621 | 0.657 | 0.496 | 0.968 | 0.404 | 0.530 | 0.867 | 0.835 | 0.314 | 0.207 | **2362.654** | 0.645 | 0.665 | 0.940 | 0.946 | 0.555 | 0.541 | 0.764 | 0.793 | 0.242 | 0.245 |
| **2367.776** | 0.576 | 0.793 | 0.653 | 0.727 | 0.640 | 0.712 | 0.584 | 0.594 | 0.370 | 0.301 | **2367.776** | 0.772 | 0.796 | 0.677 | 0.701 | 0.768 | 0.728 | 0.538 | 0.565 | 0.344 | 0.319 |
| **2367.819** | 0.825 | 0.810 | 0.882 | 0.672 | 0.766 | 0.890 | 0.547 | 0.555 | 0.271 | 0.264 | **2367.819** | 0.872 | 0.859 | 0.620 | 0.625 | 0.830 | 0.813 | 0.513 | 0.541 | 0.337 | 0.302 |
| **2368.796** | 0.747 | 0.789 | 0.796 | 0.739 | 0.724 | 0.877 | 0.591 | 0.595 | 0.287 | 0.238 | **2368.796** | 0.801 | 0.781 | 0.705 | 0.730 | 0.805 | 0.769 | 0.548 | 0.580 | 0.321 | 0.295 |
| **2377.851** | 0.734 | 0.798 | 0.839 | 0.815 | 0.752 | 0.870 | 0.602 | 0.659 | 0.267 | 0.191 | **2377.851** | 0.666 | 0.638 | 0.692 | 0.741 | 0.675 | 0.674 | 0.601 | 0.668 | 0.341 | 0.298 |
| **2391.830** | 0.503 | 0.534 | 0.665 | 0.852 | 0.589 | 0.917 | 0.820 | 0.697 | 0.272 | 0.179 | **2391.830** | 0.467 | 0.456 | 0.944 | 0.959 | 0.830 | 0.759 | 0.787 | 0.796 | 0.175 | 0.202 |
| **2392.851** | 0.430 | 0.473 | 0.905 | 0.797 | 0.815 | 0.959 | 0.791 | 0.670 | 0.188 | 0.194 | **2392.851** | 0.414 | 0.431 | 0.890 | 0.892 | 0.861 | 0.848 | 0.747 | 0.742 | 0.206 | 0.215 |
| **2393.846** | 0.578 | 0.685 | 0.965 | 0.853 | 0.871 | 0.973 | 0.747 | 0.690 | 0.158 | 0.143 | **2393.846** | 0.465 | 0.457 | 0.906 | 0.920 | 0.866 | 0.827 | 0.755 | 0.787 | 0.191 | 0.189 |
| **2402.883** | 0.916 | 0.883 | 0.542 | 0.509 | 0.471 | 0.527 | 0.436 | 0.488 | 0.465 | 0.449 | **2402.883** | 0.991 | 0.989 | 0.479 | 0.509 | 0.474 | 0.480 | 0.443 | 0.475 | 0.480 | 0.442 |
| **2406.773** | 0.801 | 0.787 | 0.978 | 0.520 | 0.921 | 0.978 | 0.386 | 0.489 | 0.294 | 0.291 | **2406.773** | 0.873 | 0.861 | 0.488 | 0.513 | 0.960 | 0.953 | 0.422 | 0.474 | 0.375 | 0.305 |
| **2413.812** | 0.477 | 0.531 | 0.549 | 0.950 | 0.431 | 0.638 | 0.901 | 0.822 | 0.305 | 0.193 | **2413.812** | 0.453 | 0.462 | 0.987 | 0.989 | 0.510 | 0.506 | 0.877 | 0.875 | 0.201 | 0.237 |
| **2414.833** | 0.400 | 0.464 | 0.921 | 0.906 | 0.812 | 0.944 | 0.851 | 0.749 | 0.160 | 0.139 | **2414.833** | 0.393 | 0.397 | 0.953 | 0.950 | 0.863 | 0.842 | 0.819 | 0.803 | 0.153 | 0.181 |
| **2418.877** | 0.886 | 0.889 | 0.846 | 0.479 | 0.748 | 0.822 | 0.431 | 0.456 | 0.334 | 0.362 | **2418.877** | 0.904 | 0.902 | 0.449 | 0.455 | 0.662 | 0.684 | 0.443 | 0.470 | 0.444 | 0.397 |
| **2423.844** | 0.627 | 0.669 | 0.649 | 0.966 | 0.575 | 0.632 | 0.751 | 0.841 | 0.295 | 0.164 | **2423.844** | 0.561 | 0.588 | 0.951 | 0.970 | 0.500 | 0.491 | 0.829 | 0.862 | 0.225 | 0.232 |
| **2433.877** | 0.561 | 0.604 | 0.981 | 0.521 | 0.969 | 0.994 | 0.439 | 0.482 | 0.291 | 0.313 | **2433.877** | 0.554 | 0.548 | 0.469 | 0.474 | 0.983 | 0.981 | 0.453 | 0.474 | 0.395 | 0.350 |
| **2434.872** | 0.603 | 0.644 | 0.980 | 0.518 | 0.969 | 0.993 | 0.430 | 0.480 | 0.288 | 0.310 | **2434.872** | 0.573 | 0.550 | 0.465 | 0.476 | 0.980 | 0.978 | 0.448 | 0.474 | 0.396 | 0.350 |
| **2435.794** | 0.448 | 0.507 | 0.356 | 0.979 | 0.277 | 0.460 | 0.954 | 0.893 | 0.362 | 0.213 | **2435.794** | 0.411 | 0.445 | 0.996 | 0.996 | 0.386 | 0.388 | 0.935 | 0.930 | 0.201 | 0.247 |
| **2448.852** | 0.700 | 0.706 | 0.927 | 0.702 | 0.866 | 0.982 | 0.539 | 0.582 | 0.256 | 0.222 | **2448.852** | 0.700 | 0.699 | 0.646 | 0.687 | 0.961 | 0.931 | 0.522 | 0.586 | 0.315 | 0.256 |
| **2449.872** | 0.680 | 0.705 | 0.980 | 0.648 | 0.940 | 0.990 | 0.519 | 0.564 | 0.237 | 0.237 | **2449.872** | 0.698 | 0.676 | 0.591 | 0.617 | 0.970 | 0.959 | 0.502 | 0.548 | 0.332 | 0.279 |
| **2450.892** | 0.740 | 0.747 | 0.982 | 0.549 | 0.948 | 0.989 | 0.443 | 0.512 | 0.264 | 0.275 | **2450.892** | 0.751 | 0.740 | 0.483 | 0.519 | 0.957 | 0.944 | 0.443 | 0.482 | 0.380 | 0.322 |
| **2463.771** | 0.856 | 0.841 | 0.777 | 0.550 | 0.641 | 0.776 | 0.425 | 0.497 | 0.390 | 0.351 | **2463.771** | 0.945 | 0.947 | 0.483 | 0.514 | 0.742 | 0.710 | 0.435 | 0.485 | 0.418 | 0.366 |
| **2463.795** | 0.866 | 0.842 | 0.825 | 0.554 | 0.671 | 0.789 | 0.417 | 0.500 | 0.375 | 0.344 | **2463.795** | 0.944 | 0.941 | 0.483 | 0.518 | 0.739 | 0.712 | 0.439 | 0.486 | 0.416 | 0.366 |
| **2465.867** | 0.572 | 0.560 | 0.748 | 0.924 | 0.633 | 0.782 | 0.808 | 0.773 | 0.239 | 0.169 | **2465.867** | 0.514 | 0.516 | 0.934 | 0.946 | 0.630 | 0.632 | 0.801 | 0.823 | 0.215 | 0.221 |
| **2466.887** | 0.698 | 0.755 | 0.983 | 0.778 | 0.972 | 0.996 | 0.667 | 0.673 | 0.142 | 0.145 | **2466.887** | 0.652 | 0.641 | 0.781 | 0.798 | 0.989 | 0.984 | 0.654 | 0.687 | 0.212 | 0.177 |
| **2487.849** | 0.566 | 0.642 | 0.512 | 0.990 | 0.407 | 0.584 | 0.925 | 0.904 | 0.290 | 0.142 | **2487.849** | 0.464 | 0.491 | 0.982 | 0.985 | 0.395 | 0.406 | 0.905 | 0.926 | 0.213 | 0.237 |
| **2490.899** | 0.626 | 0.580 | 0.928 | 0.391 | 0.803 | 0.951 | 0.753 | 0.515 | 0.175 | 0.323 | **2490.899** | 0.704 | 0.697 | 0.479 | 0.416 | 0.921 | 0.912 | 0.543 | 0.529 | 0.331 | 0.323 |
| **2498.834** | 0.517 | 0.570 | 0.987 | 0.543 | 0.973 | 0.997 | 0.458 | 0.492 | 0.286 | 0.307 | **2498.834** | 0.482 | 0.486 | 0.486 | 0.460 | 0.994 | 0.993 | 0.460 | 0.482 | 0.394 | 0.352 |
| **2502.725** | 0.640 | 0.729 | 0.543 | 0.892 | 0.495 | 0.562 | 0.679 | 0.727 | 0.374 | 0.263 | **2502.725** | 0.640 | 0.642 | 0.829 | 0.859 | 0.646 | 0.615 | 0.652 | 0.694 | 0.304 | 0.290 |
| **2504.821** | 0.849 | 0.803 | 0.640 | 0.578 | 0.538 | 0.667 | 0.516 | 0.520 | 0.397 | 0.379 | **2504.821** | 0.934 | 0.922 | 0.556 | 0.580 | 0.599 | 0.591 | 0.493 | 0.528 | 0.412 | 0.378 |
| **2507.914** | 0.914 | 0.884 | 0.962 | 0.496 | 0.874 | 0.950 | 0.368 | 0.475 | 0.303 | 0.301 | **2507.914** | 0.978 | 0.976 | 0.452 | 0.479 | 0.931 | 0.917 | 0.394 | 0.449 | 0.392 | 0.317 |
| **2516.771** | 0.539 | 0.588 | 0.778 | 0.913 | 0.687 | 0.815 | 0.751 | 0.746 | 0.253 | 0.172 | **2516.771** | 0.525 | 0.534 | 0.849 | 0.859 | 0.752 | 0.730 | 0.693 | 0.712 | 0.261 | 0.258 |
| **2523.909** | 0.860 | 0.887 | 0.950 | 0.659 | 0.891 | 0.960 | 0.477 | 0.559 | 0.249 | 0.227 | **2523.909** | 0.846 | 0.828 | 0.577 | 0.625 | 0.889 | 0.879 | 0.476 | 0.549 | 0.353 | 0.280 |
| **2539.904** | 0.654 | 0.759 | 0.797 | 0.931 | 0.728 | 0.862 | 0.754 | 0.764 | 0.214 | 0.120 | **2539.904** | 0.471 | 0.469 | 0.877 | 0.906 | 0.550 | 0.553 | 0.794 | 0.836 | 0.254 | 0.252 |
| **2548.940** | 0.885 | 0.856 | 0.761 | 0.503 | 0.666 | 0.762 | 0.423 | 0.459 | 0.379 | 0.384 | **2548.940** | 0.952 | 0.940 | 0.479 | 0.485 | 0.589 | 0.616 | 0.428 | 0.467 | 0.464 | 0.412 |
| **2564.935** | 0.931 | 0.914 | 0.756 | 0.491 | 0.640 | 0.722 | 0.393 | 0.463 | 0.397 | 0.390 | **2564.935** | 0.962 | 0.956 | 0.425 | 0.439 | 0.583 | 0.599 | 0.423 | 0.458 | 0.474 | 0.425 |
| **2568.826** | 0.687 | 0.722 | 0.982 | 0.690 | 0.955 | 0.990 | 0.493 | 0.570 | 0.244 | 0.226 | **2568.826** | 0.710 | 0.704 | 0.556 | 0.583 | 0.976 | 0.969 | 0.491 | 0.534 | 0.340 | 0.283 |
| **2570.855** | 0.707 | 0.740 | 0.979 | 0.618 | 0.951 | 0.981 | 0.455 | 0.522 | 0.263 | 0.266 | **2570.855** | 0.729 | 0.712 | 0.518 | 0.535 | 0.911 | 0.919 | 0.480 | 0.502 | 0.367 | 0.322 |
| **2573.726** | 0.569 | 0.579 | 0.536 | 0.857 | 0.465 | 0.624 | 0.821 | 0.736 | 0.321 | 0.256 | **2573.726** | 0.581 | 0.575 | 0.764 | 0.767 | 0.609 | 0.614 | 0.654 | 0.658 | 0.328 | 0.331 |
| **2609.829** | 0.881 | 0.841 | 0.874 | 0.506 | 0.732 | 0.856 | 0.388 | 0.474 | 0.360 | 0.341 | **2609.829** | 0.968 | 0.966 | 0.460 | 0.500 | 0.828 | 0.802 | 0.413 | 0.460 | 0.408 | 0.349 |
| **2609.853** | 0.884 | 0.842 | 0.887 | 0.508 | 0.750 | 0.858 | 0.386 | 0.475 | 0.352 | 0.339 | **2609.853** | 0.969 | 0.967 | 0.461 | 0.490 | 0.827 | 0.806 | 0.413 | 0.456 | 0.408 | 0.350 |
| **2610.806** | 0.739 | 0.835 | 0.685 | 0.494 | 0.664 | 0.809 | 0.445 | 0.472 | 0.404 | 0.362 | **2610.806** | 0.916 | 0.944 | 0.478 | 0.495 | 0.847 | 0.765 | 0.415 | 0.471 | 0.407 | 0.359 |
| **2610.904** | 0.847 | 0.582 | 0.904 | 0.578 | 0.797 | 0.897 | 0.441 | 0.515 | 0.310 | 0.325 | **2610.904** | 0.641 | 0.570 | 0.531 | 0.602 | 0.771 | 0.824 | 0.496 | 0.511 | 0.406 | 0.364 |
| **2612.945** | 0.711 | 0.715 | 0.989 | 0.511 | 0.974 | 0.997 | 0.433 | 0.491 | 0.265 | 0.293 | **2612.945** | 0.687 | 0.681 | 0.442 | 0.471 | 0.995 | 0.995 | 0.434 | 0.470 | 0.389 | 0.327 |
| **2646.809** | 0.474 | 0.508 | 0.530 | 0.944 | 0.446 | 0.603 | 0.854 | 0.801 | 0.328 | 0.223 | **2646.809** | 0.486 | 0.482 | 0.944 | 0.942 | 0.571 | 0.565 | 0.770 | 0.768 | 0.255 | 0.281 |
| **2648.783** | 0.518 | 0.533 | 0.563 | 0.923 | 0.511 | 0.597 | 0.733 | 0.765 | 0.358 | 0.247 | **2648.783** | 0.596 | 0.602 | 0.894 | 0.913 | 0.643 | 0.617 | 0.699 | 0.713 | 0.272 | 0.279 |
| **2653.972** | 0.838 | 0.833 | 0.969 | 0.495 | 0.941 | 0.985 | 0.373 | 0.471 | 0.289 | 0.297 | **2653.972** | 0.884 | 0.861 | 0.435 | 0.452 | 0.936 | 0.929 | 0.407 | 0.442 | 0.397 | 0.338 |
| **2654.868** | 0.724 | 0.774 | 0.569 | 0.907 | 0.497 | 0.539 | 0.767 | 0.753 | 0.307 | 0.248 | **2654.868** | 0.630 | 0.633 | 0.822 | 0.845 | 0.512 | 0.527 | 0.664 | 0.710 | 0.334 | 0.313 |
| **2666.874** | 0.785 | 0.770 | 0.866 | 0.561 | 0.770 | 0.877 | 0.424 | 0.486 | 0.345 | 0.328 | **2666.874** | 0.849 | 0.838 | 0.496 | 0.512 | 0.750 | 0.713 | 0.442 | 0.481 | 0.423 | 0.385 |
| **2669.967** | 0.876 | 0.878 | 0.980 | 0.657 | 0.932 | 0.984 | 0.488 | 0.576 | 0.221 | 0.208 | **2669.967** | 0.883 | 0.872 | 0.625 | 0.659 | 0.961 | 0.946 | 0.486 | 0.562 | 0.317 | 0.240 |
| **2685.962** | 0.699 | 0.754 | 0.899 | 0.655 | 0.862 | 0.926 | 0.509 | 0.559 | 0.277 | 0.256 | **2685.962** | 0.573 | 0.573 | 0.551 | 0.581 | 0.773 | 0.777 | 0.501 | 0.551 | 0.408 | 0.358 |
| **2710.993** | 0.919 | 0.893 | 0.774 | 0.489 | 0.659 | 0.755 | 0.404 | 0.463 | 0.384 | 0.380 | **2710.993** | 0.965 | 0.965 | 0.414 | 0.448 | 0.613 | 0.632 | 0.414 | 0.437 | 0.472 | 0.424 |
| **2711.767** | 0.797 | 0.813 | 0.596 | 0.750 | 0.497 | 0.607 | 0.620 | 0.615 | 0.370 | 0.322 | **2711.767** | 0.828 | 0.828 | 0.685 | 0.696 | 0.630 | 0.618 | 0.545 | 0.588 | 0.370 | 0.337 |
| **2711.791** | 0.804 | 0.808 | 0.605 | 0.737 | 0.505 | 0.623 | 0.619 | 0.608 | 0.365 | 0.323 | **2711.791** | 0.839 | 0.824 | 0.650 | 0.664 | 0.653 | 0.645 | 0.535 | 0.573 | 0.372 | 0.341 |
| **2712.843** | 0.722 | 0.553 | 0.760 | 0.476 | 0.645 | 0.849 | 0.559 | 0.499 | 0.342 | 0.367 | **2712.843** | 0.649 | 0.642 | 0.514 | 0.488 | 0.754 | 0.758 | 0.490 | 0.496 | 0.415 | 0.395 |
| **2714.884** | 0.714 | 0.733 | 0.987 | 0.528 | 0.968 | 0.993 | 0.398 | 0.480 | 0.285 | 0.298 | **2714.884** | 0.719 | 0.726 | 0.474 | 0.485 | 0.993 | 0.991 | 0.427 | 0.469 | 0.386 | 0.320 |
| **2716.913** | 0.720 | 0.736 | 0.986 | 0.543 | 0.967 | 0.993 | 0.417 | 0.487 | 0.274 | 0.291 | **2716.913** | 0.695 | 0.678 | 0.481 | 0.498 | 0.963 | 0.955 | 0.439 | 0.472 | 0.389 | 0.337 |
| **2717.934** | 0.677 | 0.690 | 0.984 | 0.561 | 0.964 | 0.972 | 0.437 | 0.494 | 0.273 | 0.299 | **2717.934** | 0.624 | 0.617 | 0.480 | 0.498 | 0.859 | 0.908 | 0.453 | 0.474 | 0.418 | 0.360 |
| **2758.978** | 0.678 | 0.720 | 0.948 | 0.722 | 0.865 | 0.937 | 0.612 | 0.595 | 0.218 | 0.227 | **2758.978** | 0.567 | 0.564 | 0.686 | 0.720 | 0.825 | 0.812 | 0.571 | 0.613 | 0.334 | 0.298 |
| **2771.906** | 0.811 | 0.806 | 0.960 | 0.535 | 0.890 | 0.964 | 0.397 | 0.494 | 0.300 | 0.289 | **2771.906** | 0.844 | 0.845 | 0.434 | 0.473 | 0.917 | 0.890 | 0.407 | 0.457 | 0.407 | 0.343 |
| **2772.859** | 0.743 | 0.824 | 0.728 | 0.540 | 0.684 | 0.874 | 0.448 | 0.502 | 0.389 | 0.315 | **2772.859** | 0.828 | 0.862 | 0.460 | 0.489 | 0.873 | 0.798 | 0.430 | 0.476 | 0.404 | 0.358 |
| **2779.965** | 0.626 | 0.665 | 0.867 | 0.794 | 0.765 | 0.912 | 0.722 | 0.670 | 0.214 | 0.187 | **2779.965** | 0.551 | 0.552 | 0.792 | 0.795 | 0.769 | 0.737 | 0.650 | 0.674 | 0.288 | 0.282 |
| **2800.926** | 0.618 | 0.667 | 0.515 | 0.966 | 0.430 | 0.613 | 0.885 | 0.847 | 0.293 | 0.167 | **2800.926** | 0.543 | 0.558 | 0.954 | 0.945 | 0.539 | 0.524 | 0.821 | 0.845 | 0.221 | 0.238 |
| **2812.932** | 0.803 | 0.778 | 0.925 | 0.533 | 0.854 | 0.937 | 0.393 | 0.465 | 0.321 | 0.322 | **2812.932** | 0.891 | 0.883 | 0.437 | 0.467 | 0.842 | 0.819 | 0.418 | 0.462 | 0.414 | 0.358 |
| **2813.885** | 0.700 | 0.788 | 0.673 | 0.529 | 0.659 | 0.817 | 0.452 | 0.475 | 0.411 | 0.359 | **2813.885** | 0.860 | 0.887 | 0.454 | 0.486 | 0.798 | 0.735 | 0.416 | 0.453 | 0.430 | 0.389 |
| **2813.984** | 0.794 | 0.613 | 0.921 | 0.556 | 0.851 | 0.928 | 0.410 | 0.487 | 0.315 | 0.330 | **2813.984** | 0.641 | 0.598 | 0.473 | 0.515 | 0.815 | 0.842 | 0.458 | 0.482 | 0.425 | 0.379 |
| **2815.004** | 0.683 | 0.579 | 0.905 | 0.561 | 0.845 | 0.893 | 0.444 | 0.501 | 0.321 | 0.338 | **2815.004** | 0.591 | 0.565 | 0.488 | 0.509 | 0.815 | 0.823 | 0.465 | 0.488 | 0.425 | 0.388 |
| **2816.025** | 0.863 | 0.849 | 0.985 | 0.478 | 0.935 | 0.985 | 0.361 | 0.472 | 0.291 | 0.296 | **2816.025** | 0.888 | 0.864 | 0.392 | 0.443 | 0.973 | 0.966 | 0.381 | 0.436 | 0.408 | 0.329 |
| **2832.020** | 0.711 | 0.766 | 0.951 | 0.658 | 0.922 | 0.976 | 0.526 | 0.564 | 0.238 | 0.233 | **2832.020** | 0.626 | 0.612 | 0.539 | 0.562 | 0.891 | 0.877 | 0.506 | 0.548 | 0.367 | 0.323 |
| **2852.981** | 0.545 | 0.610 | 0.692 | 0.928 | 0.599 | 0.768 | 0.763 | 0.767 | 0.288 | 0.172 | **2852.981** | 0.434 | 0.459 | 0.807 | 0.811 | 0.528 | 0.538 | 0.710 | 0.735 | 0.328 | 0.326 |
| **2855.010** | 0.569 | 0.623 | 0.856 | 0.832 | 0.772 | 0.712 | 0.707 | 0.715 | 0.231 | 0.234 | **2855.010** | 0.520 | 0.519 | 0.665 | 0.682 | 0.624 | 0.688 | 0.625 | 0.637 | 0.362 | 0.336 |
| **2873.046** | 0.860 | 0.871 | 0.884 | 0.492 | 0.797 | 0.891 | 0.395 | 0.461 | 0.336 | 0.334 | **2873.046** | 0.798 | 0.804 | 0.380 | 0.409 | 0.649 | 0.650 | 0.406 | 0.431 | 0.494 | 0.451 |
| **2905.036** | 0.634 | 0.707 | 0.920 | 0.818 | 0.800 | 0.931 | 0.630 | 0.632 | 0.243 | 0.198 | **2905.036** | 0.453 | 0.450 | 0.642 | 0.696 | 0.759 | 0.722 | 0.582 | 0.625 | 0.365 | 0.342 |
| **2917.963** | 0.839 | 0.813 | 0.946 | 0.486 | 0.847 | 0.941 | 0.375 | 0.468 | 0.324 | 0.319 | **2917.963** | 0.880 | 0.877 | 0.389 | 0.432 | 0.895 | 0.884 | 0.389 | 0.430 | 0.425 | 0.349 |
| **2919.993** | 0.835 | 0.791 | 0.950 | 0.475 | 0.856 | 0.948 | 0.384 | 0.460 | 0.316 | 0.326 | **2919.993** | 0.819 | 0.801 | 0.411 | 0.425 | 0.855 | 0.856 | 0.404 | 0.429 | 0.432 | 0.382 |
| **2967.946** | 0.691 | 0.720 | 0.754 | 0.849 | 0.675 | 0.796 | 0.708 | 0.684 | 0.257 | 0.209 | **2967.946** | 0.583 | 0.597 | 0.750 | 0.753 | 0.777 | 0.747 | 0.588 | 0.622 | 0.326 | 0.306 |
| **2967.977** | 0.673 | 0.628 | 0.837 | 0.673 | 0.768 | 0.955 | 0.682 | 0.600 | 0.230 | 0.233 | **2967.977** | 0.538 | 0.537 | 0.611 | 0.611 | 0.859 | 0.815 | 0.532 | 0.553 | 0.362 | 0.347 |
| **2968.998** | 0.638 | 0.506 | 0.912 | 0.467 | 0.830 | 0.980 | 0.662 | 0.510 | 0.215 | 0.318 | **2968.998** | 0.496 | 0.509 | 0.538 | 0.515 | 0.900 | 0.871 | 0.502 | 0.513 | 0.385 | 0.366 |
| **2978.077** | 0.745 | 0.765 | 0.976 | 0.473 | 0.955 | 0.990 | 0.398 | 0.463 | 0.286 | 0.311 | **2978.077** | 0.686 | 0.676 | 0.396 | 0.394 | 0.967 | 0.958 | 0.407 | 0.443 | 0.419 | 0.364 |
| **3019.104** | 0.836 | 0.826 | 0.908 | 0.471 | 0.814 | 0.909 | 0.384 | 0.451 | 0.336 | 0.341 | **3019.104** | 0.793 | 0.787 | 0.425 | 0.428 | 0.710 | 0.688 | 0.422 | 0.439 | 0.462 | 0.435 |
| **3114.004** | 0.602 | 0.622 | 0.909 | 0.864 | 0.792 | 0.928 | 0.756 | 0.714 | 0.185 | 0.151 | **3114.004** | 0.534 | 0.529 | 0.833 | 0.841 | 0.826 | 0.788 | 0.644 | 0.673 | 0.274 | 0.264 |
| **3114.035** | 0.592 | 0.576 | 0.911 | 0.727 | 0.795 | 0.898 | 0.746 | 0.657 | 0.190 | 0.219 | **3114.035** | 0.515 | 0.501 | 0.673 | 0.663 | 0.709 | 0.743 | 0.582 | 0.590 | 0.367 | 0.350 |
| **3115.056** | 0.603 | 0.497 | 0.818 | 0.504 | 0.696 | 0.790 | 0.764 | 0.565 | 0.226 | 0.350 | **3115.056** | 0.496 | 0.474 | 0.518 | 0.473 | 0.598 | 0.633 | 0.535 | 0.532 | 0.449 | 0.444 |
| **3124.110** | 0.750 | 0.735 | 0.979 | 0.519 | 0.945 | 0.991 | 0.427 | 0.466 | 0.273 | 0.308 | **3124.110** | 0.649 | 0.645 | 0.396 | 0.437 | 0.965 | 0.949 | 0.408 | 0.437 | 0.424 | 0.370 |
| **3124.135** | 0.694 | 0.684 | 0.980 | 0.477 | 0.954 | 0.991 | 0.430 | 0.453 | 0.277 | 0.327 | **3124.135** | 0.615 | 0.593 | 0.390 | 0.404 | 0.963 | 0.952 | 0.414 | 0.429 | 0.426 | 0.385 |
| **3132.152** | 0.503 | 0.441 | 0.747 | 0.496 | 0.656 | 0.680 | 0.459 | 0.507 | 0.433 | 0.436 | **3132.152** | 0.470 | 0.482 | 0.420 | 0.464 | 0.702 | 0.678 | 0.460 | 0.460 | 0.484 | 0.470 |
| **3154.134** | 0.512 | 0.492 | 0.720 | 0.649 | 0.645 | 0.687 | 0.533 | 0.560 | 0.399 | 0.378 | **3154.134** | 0.457 | 0.465 | 0.549 | 0.581 | 0.641 | 0.603 | 0.527 | 0.540 | 0.444 | 0.440 |
| **3293.184** | 0.580 | 0.515 | 0.705 | 0.463 | 0.649 | 0.669 | 0.466 | 0.478 | 0.424 | 0.453 | **3293.184** | 0.494 | 0.507 | 0.429 | 0.411 | 0.746 | 0.670 | 0.443 | 0.457 | 0.479 | 0.476 |
| **3294.205** | 0.515 | 0.455 | 0.642 | 0.419 | 0.627 | 0.599 | 0.462 | 0.480 | 0.454 | 0.490 | **3294.205** | 0.477 | 0.477 | 0.435 | 0.428 | 0.755 | 0.695 | 0.455 | 0.464 | 0.470 | 0.467 |

**Supplemental Table 3 (3/3): Discriminating N-glycan features of each k-means segment by area under the receiver operating characteristic curve.** Segments are color coded as blue (B), green (G), purple (P), red (R),or yellow (Y) and indicated as N-glycan IMS performed without (-) or with (+) prior multiplexed immunofluorescence (MXIF) or picrosirius red staining (PSR). Values highlighted in green indicate top 10 discriminate features for each segment. Values highlighted in yellow indicate top 30 discriminate features for each segment.   K-means segmentation analyses using Manhattan distance (k = 5) and receiver operating characteristic curves were performed in SCiLs v2026a Pro.

| m/z | MxIF- (B) | MxIF+ (B) | MxIF- (G) | MxIF+ (G) | MxIF- (P) | MxIF+ (P) | MxIF- (R) | MxIF+ (R) | MxIF- (Y) | MxIF+ (Y) | m/z | PSR- (B) | PSR+ (B) | PSR- (G) | PSR+ (G) | PSR- (P) | PSR+ (P) | PSR- (R) | PSR+ (R) | PSR- (Y) | PSR+ (Y) |
|---|---|---|---|---|---|---|---|---|---|---|---|---|---|---|---|---|---|---|---|---|---|
| **3316.187** | 0.493 | 0.456 | 0.600 | 0.437 | 0.599 | 0.567 | 0.478 | 0.481 | 0.464 | 0.499 | **3316.187** | 0.466 | 0.459 | 0.442 | 0.420 | 0.704 | 0.640 | 0.460 | 0.469 | 0.481 | 0.486 |
| **3343.210** | 0.507 | 0.501 | 0.837 | 0.482 | 0.806 | 0.834 | 0.425 | 0.497 | 0.386 | 0.380 | **3343.210** | 0.470 | 0.478 | 0.433 | 0.412 | 0.758 | 0.759 | 0.453 | 0.465 | 0.472 | 0.447 |
| **3439.242** | 0.506 | 0.472 | 0.704 | 0.485 | 0.633 | 0.615 | 0.485 | 0.488 | 0.432 | 0.470 | **3439.242** | 0.467 | 0.483 | 0.449 | 0.470 | 0.670 | 0.582 | 0.461 | 0.461 | 0.489 | 0.501 |
| **3488.247** | 0.518 | 0.446 | 0.587 | 0.512 | 0.570 | 0.567 | 0.477 | 0.536 | 0.472 | 0.457 | **3488.247** | 0.461 | 0.462 | 0.428 | 0.422 | 0.698 | 0.648 | 0.461 | 0.468 | 0.485 | 0.483 |
| **3489.242** | 0.531 | 0.482 | 0.700 | 0.538 | 0.654 | 0.754 | 0.471 | 0.519 | 0.429 | 0.392 | **3489.242** | 0.471 | 0.475 | 0.433 | 0.428 | 0.825 | 0.776 | 0.448 | 0.460 | 0.457 | 0.443 |
| **3489.268** | 0.491 | 0.439 | 0.785 | 0.505 | 0.749 | 0.762 | 0.438 | 0.521 | 0.408 | 0.396 | **3489.268** | 0.467 | 0.472 | 0.443 | 0.434 | 0.827 | 0.791 | 0.466 | 0.468 | 0.444 | 0.433 |
| **3510.229** | 0.509 | 0.464 | 0.603 | 0.520 | 0.575 | 0.583 | 0.481 | 0.530 | 0.468 | 0.453 | **3510.229** | 0.459 | 0.457 | 0.446 | 0.444 | 0.666 | 0.613 | 0.468 | 0.471 | 0.487 | 0.492 |
| **3634.305** | 0.521 | 0.463 | 0.568 | 0.529 | 0.550 | 0.527 | 0.486 | 0.542 | 0.477 | 0.465 | **3634.305** | 0.477 | 0.489 | 0.428 | 0.450 | 0.635 | 0.602 | 0.461 | 0.473 | 0.499 | 0.489 |
| **3635.300** | 0.536 | 0.513 | 0.717 | 0.549 | 0.659 | 0.743 | 0.478 | 0.523 | 0.420 | 0.389 | **3635.300** | 0.480 | 0.475 | 0.435 | 0.453 | 0.778 | 0.757 | 0.457 | 0.472 | 0.462 | 0.440 |
| **3656.287** | 0.525 | 0.480 | 0.548 | 0.534 | 0.536 | 0.524 | 0.485 | 0.536 | 0.485 | 0.467 | **3656.287** | 0.474 | 0.473 | 0.440 | 0.418 | 0.667 | 0.605 | 0.458 | 0.463 | 0.492 | 0.499 |
| **3699.262** | 0.499 | 0.449 | 0.602 | 0.517 | 0.580 | 0.603 | 0.473 | 0.523 | 0.473 | 0.452 | **3699.262** | 0.463 | 0.467 | 0.424 | 0.427 | 0.703 | 0.643 | 0.455 | 0.465 | 0.487 | 0.486 |
