## Supplemental Table 4 for "Multimodal Imaging of the Cellular and Extracellular Microenvironment on the Same Formalin-Fixed Paraffin-Embedded Tissue Section"

**Supplemental Table 4 (1/3): Discriminating extracellular matrix imaging mass spectrometry features of each k-means segment by area under the receiver operating characteristic curve.** Segments are color coded as blue (B), green (G), purple (P), red (R),or yellow (Y) and indicated as N-glycan IMS performed without (-) or with (+) prior multiplexed immunofluorescence (MXIF) or picrosirius red staining (PSR). Values highlighted in green indicate top 10 discriminate features for each segment. Values highlighted in yellow indicate top 30 discriminate features for each segment. K-means segmentation analyses using Manhattan distance (k = 5) and receiver operating characteristic curves were performed in SCiLs v2026a Pro.

| m/z | MxIF- (B) | MxIF+ (B) | MxIF- (G) | MxIF+ (G) | MxIF- (P) | MxIF+ (P) | MxIF- (R) | MxIF+ (R) | MxIF- (Y) | MxIF+ (Y) | m/z | PSR- (B) | PSR+ (B) | PSR- (G) | PSR+ (G) | PSR- (P) | PSR+ (P) | PSR- (R) | PSR+ (R) | PSR- (Y) | PSR+ (Y) |
|---|---|---|---|---|---|---|---|---|---|---|---|---|---|---|---|---|---|---|---|---|---|
| 758.464 | 0.823 | 0.614 | 0.695 | 0.716 | 0.561 | 0.589 | 0.739 | 0.749 | 0.232 | 0.094 | 758.464 | 0.592 | 0.641 | 0.637 | 0.655 | 0.577 | 0.585 | 0.636 | 0.657 | 0.256 | 0.228 |
| 774.432 | 0.624 | 0.590 | 0.836 | 0.754 | 0.593 | 0.542 | 0.927 | 0.883 | 0.116 | 0.065 | 774.432 | 0.425 | 0.509 | 0.765 | 0.755 | 0.589 | 0.573 | 0.735 | 0.817 | 0.134 | 0.144 |
| 781.441 | 0.959 | 0.836 | 0.581 | 0.604 | 0.458 | 0.531 | 0.676 | 0.733 | 0.356 | 0.174 | 781.441 | 0.877 | 0.888 | 0.712 | 0.713 | 0.478 | 0.504 | 0.852 | 0.813 | 0.192 | 0.171 |
| 785.401 | 0.937 | 0.753 | 0.830 | 0.747 | 0.540 | 0.551 | 0.916 | 0.809 | 0.089 | 0.036 | 785.401 | 0.661 | 0.729 | 0.821 | 0.812 | 0.572 | 0.574 | 0.859 | 0.879 | 0.035 | 0.038 |
| 789.346 | 0.285 | 0.403 | 0.426 | 0.497 | 0.586 | 0.559 | 0.363 | 0.446 | 0.532 | 0.498 | 789.346 | 0.360 | 0.384 | 0.458 | 0.481 | 0.580 | 0.587 | 0.368 | 0.402 | 0.511 | 0.480 |
| 790.428 | 0.518 | 0.563 | 0.865 | 0.796 | 0.620 | 0.545 | 0.875 | 0.786 | 0.092 | 0.079 | 790.428 | 0.408 | 0.556 | 0.790 | 0.772 | 0.592 | 0.593 | 0.697 | 0.726 | 0.122 | 0.126 |
| 795.336 | 0.759 | 0.582 | 0.719 | 0.704 | 0.534 | 0.564 | 0.789 | 0.709 | 0.242 | 0.157 | 795.336 | 0.495 | 0.535 | 0.575 | 0.579 | 0.533 | 0.538 | 0.581 | 0.587 | 0.382 | 0.372 |
| 797.435 | 0.880 | 0.808 | 0.880 | 0.777 | 0.559 | 0.498 | 0.951 | 0.886 | 0.061 | 0.017 | 797.435 | 0.613 | 0.735 | 0.901 | 0.882 | 0.516 | 0.521 | 0.917 | 0.945 | 0.015 | 0.015 |
| 801.392 | 0.672 | 0.536 | 0.839 | 0.820 | 0.621 | 0.552 | 0.905 | 0.851 | 0.117 | 0.035 | 801.392 | 0.396 | 0.480 | 0.757 | 0.757 | 0.629 | 0.627 | 0.681 | 0.745 | 0.119 | 0.116 |
| 804.443 | 0.647 | 0.587 | 0.781 | 0.777 | 0.590 | 0.540 | 0.815 | 0.786 | 0.181 | 0.091 | 804.443 | 0.362 | 0.469 | 0.677 | 0.675 | 0.609 | 0.606 | 0.569 | 0.623 | 0.242 | 0.234 |
| 807.38 | 0.920 | 0.665 | 0.729 | 0.679 | 0.487 | 0.547 | 0.844 | 0.723 | 0.219 | 0.160 | 807.38 | 0.525 | 0.540 | 0.564 | 0.566 | 0.556 | 0.561 | 0.571 | 0.569 | 0.367 | 0.364 |
| 809.47 | 0.964 | 0.925 | 0.779 | 0.679 | 0.499 | 0.526 | 0.879 | 0.807 | 0.155 | 0.058 | 809.47 | 0.776 | 0.841 | 0.875 | 0.860 | 0.502 | 0.517 | 0.960 | 0.956 | 0.015 | 0.013 |
| 812.41 | 0.802 | 0.761 | 0.907 | 0.815 | 0.583 | 0.465 | 0.977 | 0.945 | 0.043 | 0.012 | 812.41 | 0.517 | 0.656 | 0.887 | 0.875 | 0.538 | 0.533 | 0.898 | 0.942 | 0.024 | 0.025 |
| 815.376 | 0.623 | 0.530 | 0.865 | 0.836 | 0.620 | 0.514 | 0.928 | 0.910 | 0.103 | 0.039 | 815.376 | 0.417 | 0.496 | 0.666 | 0.643 | 0.556 | 0.547 | 0.641 | 0.667 | 0.278 | 0.299 |
| 819.41 | 0.898 | 0.697 | 0.786 | 0.720 | 0.503 | 0.495 | 0.898 | 0.805 | 0.170 | 0.137 | 819.41 | 0.524 | 0.548 | 0.659 | 0.638 | 0.561 | 0.556 | 0.646 | 0.646 | 0.261 | 0.291 |
| 822.394 | 0.732 | 0.565 | 0.837 | 0.806 | 0.601 | 0.568 | 0.893 | 0.806 | 0.108 | 0.037 | 822.394 | 0.487 | 0.562 | 0.715 | 0.723 | 0.611 | 0.605 | 0.682 | 0.735 | 0.159 | 0.150 |
| 827.41 | 0.833 | 0.708 | 0.892 | 0.818 | 0.593 | 0.506 | 0.960 | 0.876 | 0.043 | 0.014 | 827.41 | 0.502 | 0.626 | 0.853 | 0.850 | 0.588 | 0.575 | 0.835 | 0.894 | 0.023 | 0.023 |
| 827.447 | 0.952 | 0.840 | 0.818 | 0.743 | 0.513 | 0.503 | 0.927 | 0.886 | 0.113 | 0.027 | 827.447 | 0.796 | 0.852 | 0.870 | 0.854 | 0.501 | 0.519 | 0.948 | 0.946 | 0.021 | 0.017 |
| 831.453 | 0.925 | 0.857 | 0.673 | 0.630 | 0.512 | 0.526 | 0.755 | 0.713 | 0.254 | 0.159 | 831.453 | 0.495 | 0.573 | 0.599 | 0.580 | 0.569 | 0.558 | 0.618 | 0.604 | 0.317 | 0.341 |
| 833.401 | 0.670 | 0.580 | 0.875 | 0.841 | 0.602 | 0.498 | 0.932 | 0.909 | 0.098 | 0.034 | 833.401 | 0.494 | 0.592 | 0.760 | 0.745 | 0.558 | 0.560 | 0.736 | 0.773 | 0.159 | 0.160 |
| 835.383 | 0.875 | 0.741 | 0.823 | 0.757 | 0.526 | 0.512 | 0.933 | 0.867 | 0.130 | 0.048 | 835.383 | 0.669 | 0.767 | 0.835 | 0.816 | 0.525 | 0.526 | 0.891 | 0.906 | 0.060 | 0.067 |
| 841.423 | 0.768 | 0.770 | 0.869 | 0.793 | 0.600 | 0.474 | 0.961 | 0.945 | 0.082 | 0.018 | 841.423 | 0.446 | 0.561 | 0.851 | 0.846 | 0.561 | 0.556 | 0.849 | 0.902 | 0.056 | 0.055 |
| 843.403 | 0.871 | 0.731 | 0.920 | 0.837 | 0.566 | 0.456 | 0.991 | 0.967 | 0.025 | 0.005 | 843.403 | 0.666 | 0.759 | 0.894 | 0.879 | 0.514 | 0.521 | 0.942 | 0.960 | 0.008 | 0.008 |
| 843.437 | 0.789 | 0.637 | 0.911 | 0.837 | 0.587 | 0.483 | 0.976 | 0.937 | 0.038 | 0.022 | 843.437 | 0.577 | 0.657 | 0.886 | 0.866 | 0.536 | 0.542 | 0.899 | 0.925 | 0.018 | 0.028 |
| 849.395 | 0.834 | 0.698 | 0.861 | 0.807 | 0.553 | 0.484 | 0.955 | 0.899 | 0.100 | 0.039 | 849.395 | 0.446 | 0.547 | 0.732 | 0.767 | 0.609 | 0.581 | 0.698 | 0.793 | 0.148 | 0.126 |
| 855.405 | 0.607 | 0.557 | 0.827 | 0.804 | 0.588 | 0.488 | 0.915 | 0.894 | 0.151 | 0.088 | 855.405 | 0.497 | 0.581 | 0.764 | 0.761 | 0.556 | 0.556 | 0.785 | 0.812 | 0.144 | 0.142 |
| 858.412 | 0.713 | 0.620 | 0.842 | 0.794 | 0.589 | 0.529 | 0.938 | 0.876 | 0.110 | 0.039 | 858.412 | 0.434 | 0.501 | 0.757 | 0.758 | 0.610 | 0.601 | 0.736 | 0.778 | 0.118 | 0.127 |
| 861.431 | 0.902 | 0.872 | 0.742 | 0.713 | 0.537 | 0.521 | 0.835 | 0.822 | 0.182 | 0.049 | 861.431 | 0.661 | 0.796 | 0.845 | 0.824 | 0.502 | 0.506 | 0.910 | 0.912 | 0.072 | 0.072 |
| 863.401 | 0.763 | 0.651 | 0.825 | 0.749 | 0.556 | 0.513 | 0.935 | 0.858 | 0.126 | 0.089 | 863.401 | 0.530 | 0.581 | 0.653 | 0.648 | 0.573 | 0.576 | 0.652 | 0.660 | 0.252 | 0.253 |
| 865.389 | 0.930 | 0.756 | 0.851 | 0.784 | 0.503 | 0.466 | 0.962 | 0.911 | 0.104 | 0.052 | 865.389 | 0.723 | 0.780 | 0.757 | 0.748 | 0.515 | 0.517 | 0.845 | 0.860 | 0.141 | 0.138 |
| 869.458 | 0.894 | 0.814 | 0.811 | 0.758 | 0.549 | 0.499 | 0.913 | 0.892 | 0.115 | 0.027 | 869.458 | 0.740 | 0.820 | 0.861 | 0.846 | 0.511 | 0.522 | 0.924 | 0.924 | 0.033 | 0.032 |
| 870.419 | 0.916 | 0.912 | 0.857 | 0.727 | 0.547 | 0.512 | 0.934 | 0.836 | 0.073 | 0.028 | 870.419 | 0.598 | 0.718 | 0.903 | 0.881 | 0.511 | 0.517 | 0.943 | 0.963 | 0.013 | 0.018 |
| 872.428 | 0.711 | 0.675 | 0.879 | 0.795 | 0.587 | 0.482 | 0.964 | 0.930 | 0.087 | 0.046 | 872.428 | 0.441 | 0.548 | 0.849 | 0.835 | 0.552 | 0.554 | 0.841 | 0.904 | 0.069 | 0.068 |
| 872.467 | 0.792 | 0.716 | 0.874 | 0.820 | 0.567 | 0.471 | 0.961 | 0.940 | 0.094 | 0.019 | 872.467 | 0.521 | 0.682 | 0.866 | 0.860 | 0.530 | 0.527 | 0.870 | 0.903 | 0.057 | 0.049 |
| 875.394 | 0.842 | 0.693 | 0.917 | 0.841 | 0.578 | 0.477 | 0.977 | 0.930 | 0.029 | 0.009 | 875.394 | 0.564 | 0.681 | 0.891 | 0.882 | 0.543 | 0.541 | 0.877 | 0.917 | 0.014 | 0.014 |
| 881.35 | 0.845 | 0.708 | 0.805 | 0.801 | 0.509 | 0.485 | 0.926 | 0.909 | 0.172 | 0.036 | 881.35 | 0.641 | 0.746 | 0.829 | 0.827 | 0.544 | 0.533 | 0.874 | 0.914 | 0.056 | 0.053 |
| 884.431 | 0.958 | 0.942 | 0.808 | 0.698 | 0.510 | 0.509 | 0.920 | 0.847 | 0.121 | 0.038 | 884.431 | 0.793 | 0.874 | 0.884 | 0.864 | 0.491 | 0.504 | 0.970 | 0.971 | 0.013 | 0.012 |
| 886.454 | 0.707 | 0.665 | 0.932 | 0.851 | 0.615 | 0.455 | 0.988 | 0.983 | 0.044 | 0.012 | 886.454 | 0.540 | 0.648 | 0.889 | 0.867 | 0.532 | 0.542 | 0.917 | 0.934 | 0.019 | 0.026 |
| 887.39 | 0.765 | 0.627 | 0.925 | 0.860 | 0.605 | 0.482 | 0.973 | 0.931 | 0.031 | 0.011 | 887.39 | 0.512 | 0.626 | 0.896 | 0.887 | 0.550 | 0.548 | 0.861 | 0.913 | 0.015 | 0.015 |
| 887.461 | 0.697 | 0.660 | 0.920 | 0.846 | 0.607 | 0.458 | 0.982 | 0.978 | 0.063 | 0.016 | 887.461 | 0.484 | 0.615 | 0.842 | 0.832 | 0.573 | 0.569 | 0.832 | 0.871 | 0.050 | 0.052 |
| 890.403 | 0.720 | 0.573 | 0.899 | 0.830 | 0.626 | 0.536 | 0.953 | 0.875 | 0.056 | 0.020 | 890.403 | 0.382 | 0.503 | 0.832 | 0.831 | 0.629 | 0.613 | 0.765 | 0.855 | 0.036 | 0.037 |
| 891.441 | 0.817 | 0.736 | 0.925 | 0.834 | 0.582 | 0.450 | 0.990 | 0.977 | 0.032 | 0.007 | 891.441 | 0.572 | 0.716 | 0.901 | 0.882 | 0.517 | 0.518 | 0.931 | 0.966 | 0.016 | 0.017 |
| 894.41 | 0.685 | 0.602 | 0.787 | 0.727 | 0.560 | 0.531 | 0.900 | 0.814 | 0.172 | 0.123 | 894.41 | 0.448 | 0.459 | 0.565 | 0.566 | 0.561 | 0.570 | 0.536 | 0.534 | 0.382 | 0.381 |
| 894.446 | 0.785 | 0.682 | 0.745 | 0.762 | 0.526 | 0.478 | 0.864 | 0.861 | 0.224 | 0.103 | 894.446 | 0.400 | 0.515 | 0.637 | 0.632 | 0.590 | 0.583 | 0.562 | 0.612 | 0.291 | 0.285 |
| 897.376 | 0.866 | 0.670 | 0.843 | 0.786 | 0.524 | 0.509 | 0.935 | 0.840 | 0.114 | 0.065 | 897.376 | 0.506 | 0.578 | 0.644 | 0.628 | 0.560 | 0.547 | 0.641 | 0.642 | 0.280 | 0.302 |
| 897.447 | 0.853 | 0.778 | 0.925 | 0.824 | 0.579 | 0.443 | 0.995 | 0.988 | 0.018 | 0.004 | 897.447 | 0.640 | 0.766 | 0.905 | 0.885 | 0.504 | 0.512 | 0.965 | 0.981 | 0.006 | 0.006 |
| 897.488 | 0.552 | 0.525 | 0.730 | 0.730 | 0.580 | 0.499 | 0.735 | 0.797 | 0.255 | 0.187 | 897.488 | 0.429 | 0.455 | 0.593 | 0.610 | 0.524 | 0.527 | 0.560 | 0.609 | 0.392 | 0.368 |
| 900.423 | 0.790 | 0.730 | 0.932 | 0.838 | 0.602 | 0.447 | 0.992 | 0.984 | 0.022 | 0.006 | 900.423 | 0.559 | 0.699 | 0.907 | 0.887 | 0.519 | 0.523 | 0.935 | 0.967 | 0.010 | 0.010 |
| 903.331 | 0.858 | 0.660 | 0.700 | 0.692 | 0.472 | 0.511 | 0.824 | 0.755 | 0.270 | 0.176 | 903.331 | 0.504 | 0.534 | 0.548 | 0.534 | 0.508 | 0.501 | 0.546 | 0.571 | 0.438 | 0.446 |
| 905.402 | 0.705 | 0.534 | 0.894 | 0.839 | 0.629 | 0.554 | 0.932 | 0.843 | 0.059 | 0.022 | 905.402 | 0.394 | 0.509 | 0.835 | 0.840 | 0.632 | 0.615 | 0.727 | 0.822 | 0.038 | 0.035 |
| 908.441 | 0.718 | 0.645 | 0.889 | 0.828 | 0.579 | 0.464 | 0.969 | 0.934 | 0.094 | 0.047 | 908.441 | 0.431 | 0.461 | 0.642 | 0.634 | 0.584 | 0.589 | 0.584 | 0.613 | 0.283 | 0.288 |
| 909.377 | 0.822 | 0.679 | 0.872 | 0.802 | 0.544 | 0.500 | 0.954 | 0.861 | 0.097 | 0.050 | 909.377 | 0.524 | 0.622 | 0.739 | 0.708 | 0.537 | 0.541 | 0.770 | 0.769 | 0.185 | 0.202 |
| 912.387 | 0.715 | 0.547 | 0.811 | 0.790 | 0.573 | 0.554 | 0.900 | 0.799 | 0.159 | 0.075 | 912.387 | 0.394 | 0.410 | 0.519 | 0.510 | 0.541 | 0.530 | 0.494 | 0.506 | 0.460 | 0.479 |
| 912.424 | 0.858 | 0.739 | 0.800 | 0.720 | 0.518 | 0.525 | 0.918 | 0.798 | 0.135 | 0.094 | 912.424 | 0.880 | 0.902 | 0.760 | 0.759 | 0.480 | 0.508 | 0.890 | 0.867 | 0.139 | 0.113 |
| 912.465 | 0.888 | 0.867 | 0.735 | 0.733 | 0.520 | 0.475 | 0.802 | 0.878 | 0.214 | 0.058 | 912.465 | 0.779 | 0.824 | 0.759 | 0.741 | 0.502 | 0.512 | 0.853 | 0.857 | 0.142 | 0.141 |
| 913.346 | 0.810 | 0.669 | 0.842 | 0.783 | 0.535 | 0.528 | 0.923 | 0.839 | 0.129 | 0.049 | 913.346 | 0.546 | 0.669 | 0.810 | 0.809 | 0.562 | 0.550 | 0.801 | 0.853 | 0.087 | 0.083 |
| 913.414 | 0.904 | 0.726 | 0.855 | 0.795 | 0.518 | 0.469 | 0.958 | 0.894 | 0.100 | 0.058 | 913.414 | 0.724 | 0.780 | 0.738 | 0.736 | 0.538 | 0.551 | 0.798 | 0.783 | 0.146 | 0.137 |
| 914.488 | 0.802 | 0.738 | 0.926 | 0.837 | 0.582 | 0.448 | 0.992 | 0.979 | 0.039 | 0.006 | 914.488 | 0.538 | 0.706 | 0.897 | 0.881 | 0.518 | 0.515 | 0.928 | 0.962 | 0.024 | 0.023 |
| 919.435 | 0.756 | 0.634 | 0.832 | 0.774 | 0.599 | 0.528 | 0.906 | 0.876 | 0.119 | 0.052 | 919.435 | 0.306 | 0.397 | 0.688 | 0.684 | 0.731 | 0.727 | 0.553 | 0.597 | 0.123 | 0.134 |
| 922.405 | 0.876 | 0.765 | 0.880 | 0.804 | 0.524 | 0.453 | 0.976 | 0.942 | 0.089 | 0.034 | 922.405 | 0.474 | 0.446 | 0.570 | 0.548 | 0.561 | 0.569 | 0.526 | 0.483 | 0.377 | 0.412 |
| 925.348 | 0.736 | 0.610 | 0.852 | 0.804 | 0.535 | 0.519 | 0.930 | 0.855 | 0.146 | 0.054 | 925.348 | 0.490 | 0.603 | 0.793 | 0.785 | 0.567 | 0.552 | 0.769 | 0.829 | 0.113 | 0.118 |
| 927.39 | 0.656 | 0.483 | 0.809 | 0.776 | 0.592 | 0.588 | 0.851 | 0.729 | 0.152 | 0.101 | 927.39 | 0.397 | 0.420 | 0.543 | 0.536 | 0.575 | 0.572 | 0.470 | 0.478 | 0.412 | 0.424 |
| 928.467 | 0.903 | 0.838 | 0.851 | 0.789 | 0.545 | 0.467 | 0.927 | 0.918 | 0.086 | 0.016 | 928.467 | 0.677 | 0.799 | 0.897 | 0.877 | 0.502 | 0.510 | 0.947 | 0.963 | 0.014 | 0.013 |
| 929.386 | 0.835 | 0.712 | 0.854 | 0.794 | 0.531 | 0.488 | 0.954 | 0.911 | 0.118 | 0.036 | 929.386 | 0.645 | 0.735 | 0.802 | 0.788 | 0.534 | 0.534 | 0.854 | 0.888 | 0.093 | 0.092 |
| 935.4 | 0.886 | 0.771 | 0.839 | 0.784 | 0.509 | 0.460 | 0.959 | 0.932 | 0.129 | 0.044 | 935.4 | 0.643 | 0.773 | 0.842 | 0.829 | 0.520 | 0.514 | 0.907 | 0.934 | 0.059 | 0.059 |
| 936.472 | 0.813 | 0.712 | 0.828 | 0.797 | 0.528 | 0.465 | 0.944 | 0.911 | 0.151 | 0.057 | 936.472 | 0.493 | 0.550 | 0.609 | 0.590 | 0.557 | 0.560 | 0.578 | 0.579 | 0.330 | 0.341 |
| 938.375 | 0.801 | 0.726 | 0.862 | 0.819 | 0.527 | 0.461 | 0.964 | 0.944 | 0.132 | 0.026 | 938.375 | 0.644 | 0.761 | 0.824 | 0.811 | 0.526 | 0.520 | 0.883 | 0.912 | 0.075 | 0.076 |
| 938.487 | 0.884 | 0.887 | 0.737 | 0.740 | 0.510 | 0.456 | 0.900 | 0.933 | 0.209 | 0.043 | 938.487 | 0.849 | 0.892 | 0.797 | 0.780 | 0.484 | 0.493 | 0.942 | 0.923 | 0.094 | 0.099 |
| 941.453 | 0.965 | 0.907 | 0.743 | 0.684 | 0.469 | 0.484 | 0.917 | 0.868 | 0.197 | 0.078 | 941.453 | 0.915 | 0.914 | 0.847 | 0.837 | 0.478 | 0.498 | 0.960 | 0.957 | 0.042 | 0.035 |
| 943.512 | 0.592 | 0.603 | 0.848 | 0.800 | 0.586 | 0.494 | 0.912 | 0.872 | 0.137 | 0.078 | 943.512 | 0.487 | 0.581 | 0.732 | 0.712 | 0.535 | 0.543 | 0.709 | 0.728 | 0.214 | 0.216 |
| 944.499 | 0.793 | 0.754 | 0.924 | 0.831 | 0.598 | 0.446 | 0.987 | 0.981 | 0.032 | 0.006 | 944.499 | 0.594 | 0.741 | 0.904 | 0.882 | 0.513 | 0.516 | 0.930 | 0.965 | 0.015 | 0.013 |
| 952.433 | 0.736 | 0.664 | 0.791 | 0.783 | 0.560 | 0.494 | 0.898 | 0.894 | 0.178 | 0.062 | 952.433 | 0.480 | 0.609 | 0.747 | 0.760 | 0.554 | 0.544 | 0.768 | 0.838 | 0.168 | 0.142 |
| 954.511 | 0.882 | 0.793 | 0.846 | 0.787 | 0.522 | 0.456 | 0.971 | 0.944 | 0.107 | 0.033 | 954.511 | 0.808 | 0.831 | 0.851 | 0.831 | 0.490 | 0.497 | 0.949 | 0.946 | 0.046 | 0.060 |
| 957.484 | 0.461 | 0.503 | 0.751 | 0.688 | 0.604 | 0.554 | 0.883 | 0.755 | 0.202 | 0.188 | 957.484 | 0.368 | 0.410 | 0.662 | 0.666 | 0.601 | 0.610 | 0.635 | 0.656 | 0.246 | 0.240 |
| 960.361 | 0.797 | 0.680 | 0.739 | 0.735 | 0.493 | 0.485 | 0.876 | 0.830 | 0.247 | 0.130 | 960.361 | 0.466 | 0.497 | 0.541 | 0.521 | 0.514 | 0.511 | 0.545 | 0.526 | 0.444 | 0.467 |
| 961.452 | 0.551 | 0.486 | 0.742 | 0.771 | 0.614 | 0.542 | 0.800 | 0.813 | 0.207 | 0.115 | 961.452 | 0.415 | 0.451 | 0.605 | 0.624 | 0.597 | 0.605 | 0.535 | 0.583 | 0.316 | 0.291 |
| 962.486 | 0.686 | 0.692 | 0.857 | 0.812 | 0.565 | 0.443 | 0.951 | 0.950 | 0.142 | 0.059 | 962.486 | 0.528 | 0.620 | 0.782 | 0.714 | 0.518 | 0.523 | 0.851 | 0.829 | 0.145 | 0.199 |
| 963.434 | 0.866 | 0.754 | 0.597 | 0.576 | 0.490 | 0.523 | 0.733 | 0.697 | 0.325 | 0.247 | 963.434 | 0.573 | 0.536 | 0.529 | 0.542 | 0.545 | 0.565 | 0.575 | 0.523 | 0.400 | 0.392 |
| 966.416 | 0.751 | 0.677 | 0.679 | 0.668 | 0.537 | 0.547 | 0.750 | 0.712 | 0.274 | 0.170 | 966.416 | 0.563 | 0.708 | 0.776 | 0.777 | 0.530 | 0.526 | 0.815 | 0.854 | 0.142 | 0.122 |
| 966.484 | 0.798 | 0.730 | 0.823 | 0.791 | 0.549 | 0.462 | 0.930 | 0.917 | 0.148 | 0.056 | 966.484 | 0.483 | 0.537 | 0.635 | 0.627 | 0.573 | 0.580 | 0.606 | 0.618 | 0.286 | 0.286 |
| 970.517 | 0.709 | 0.671 | 0.921 | 0.842 | 0.622 | 0.459 | 0.983 | 0.982 | 0.047 | 0.013 | 970.517 | 0.406 | 0.574 | 0.845 | 0.834 | 0.577 | 0.564 | 0.865 | 0.923 | 0.046 | 0.049 |
| 981.443 | 0.617 | 0.549 | 0.688 | 0.720 | 0.577 | 0.538 | 0.727 | 0.776 | 0.271 | 0.155 | 981.443 | 0.477 | 0.515 | 0.593 | 0.604 | 0.542 | 0.545 | 0.581 | 0.606 | 0.361 | 0.344 |
| 982.445 | 0.754 | 0.693 | 0.780 | 0.791 | 0.560 | 0.475 | 0.881 | 0.925 | 0.188 | 0.052 | 982.445 | 0.506 | 0.615 | 0.745 | 0.753 | 0.543 | 0.540 | 0.751 | 0.808 | 0.181 | 0.158 |
| 984.53 | 0.790 | 0.777 | 0.611 | 0.630 | 0.548 | 0.531 | 0.655 | 0.743 | 0.325 | 0.169 | 984.53 | 0.902 | 0.895 | 0.645 | 0.660 | 0.460 | 0.469 | 0.817 | 0.821 | 0.273 | 0.242 |
| 985.525 | 0.634 | 0.655 | 0.716 | 0.721 | 0.603 | 0.532 | 0.752 | 0.770 | 0.235 | 0.126 | 985.525 | 0.741 | 0.786 | 0.671 | 0.680 | 0.536 | 0.538 | 0.744 | 0.751 | 0.218 | 0.201 |
| 992.497 | 0.708 | 0.653 | 0.832 | 0.811 | 0.574 | 0.463 | 0.944 | 0.940 | 0.150 | 0.056 | 992.497 | 0.303 | 0.346 | 0.534 | 0.540 | 0.655 | 0.644 | 0.384 | 0.415 | 0.376 | 0.384 |
| 1001.53 | 0.879 | 0.840 | 0.895 | 0.792 | 0.534 | 0.446 | 0.990 | 0.967 | 0.063 | 0.014 | 1001.53 | 0.763 | 0.810 | 0.881 | 0.857 | 0.493 | 0.501 | 0.966 | 0.971 | 0.019 | 0.033 |
| 1008.46 | 0.702 | 0.642 | 0.818 | 0.817 | 0.560 | 0.468 | 0.932 | 0.949 | 0.172 | 0.046 | 1008.46 | 0.428 | 0.539 | 0.716 | 0.707 | 0.576 | 0.566 | 0.721 | 0.777 | 0.192 | 0.193 |
| 1013.62 | 0.237 | 0.432 | 0.556 | 0.526 | 0.607 | 0.566 | 0.393 | 0.467 | 0.437 | 0.448 | 1013.62 | 0.255 | 0.367 | 0.560 | 0.589 | 0.593 | 0.620 | 0.395 | 0.467 | 0.420 | 0.350 |
| 1017.48 | 0.718 | 0.625 | 0.917 | 0.849 | 0.611 | 0.476 | 0.976 | 0.957 | 0.047 | 0.016 | 1017.48 | 0.455 | 0.575 | 0.869 | 0.863 | 0.560 | 0.554 | 0.851 | 0.912 | 0.038 | 0.038 |
| 1019.51 | 0.749 | 0.766 | 0.924 | 0.819 | 0.606 | 0.450 | 0.986 | 0.974 | 0.044 | 0.011 | 1019.51 | 0.581 | 0.737 | 0.897 | 0.872 | 0.513 | 0.519 | 0.951 | 0.972 | 0.017 | 0.017 |
| 1032.48 | 0.649 | 0.576 | 0.912 | 0.852 | 0.627 | 0.488 | 0.963 | 0.947 | 0.058 | 0.022 | 1032.48 | 0.390 | 0.502 | 0.845 | 0.839 | 0.575 | 0.568 | 0.797 | 0.871 | 0.069 | 0.069 |
| 1034.58 | 0.983 | 0.921 | 0.605 | 0.582 | 0.422 | 0.504 | 0.731 | 0.743 | 0.357 | 0.185 | 1034.58 | 0.965 | 0.963 | 0.739 | 0.738 | 0.395 | 0.424 | 0.907 | 0.883 | 0.224 | 0.194 |
| 1035.59 | 0.964 | 0.907 | 0.540 | 0.573 | 0.422 | 0.510 | 0.628 | 0.730 | 0.425 | 0.196 | 1035.59 | 0.962 | 0.960 | 0.725 | 0.726 | 0.395 | 0.425 | 0.898 | 0.874 | 0.239 | 0.205 |
| 1039.47 | 0.757 | 0.662 | 0.841 | 0.805 | 0.554 | 0.474 | 0.956 | 0.924 | 0.128 | 0.054 | 1039.47 | 0.554 | 0.583 | 0.718 | 0.701 | 0.546 | 0.550 | 0.785 | 0.775 | 0.185 | 0.204 |
| 1040.49 | 0.918 | 0.884 | 0.887 | 0.775 | 0.533 | 0.456 | 0.975 | 0.947 | 0.058 | 0.010 | 1040.49 | 0.783 | 0.850 | 0.892 | 0.872 | 0.488 | 0.503 | 0.982 | 0.984 | 0.008 | 0.008 |
| 1041.55 | 0.964 | 0.863 | 0.576 | 0.561 | 0.446 | 0.532 | 0.723 | 0.743 | 0.356 | 0.193 | 1041.55 | 0.947 | 0.939 | 0.779 | 0.779 | 0.447 | 0.479 | 0.930 | 0.922 | 0.135 | 0.103 |
| 1043.49 | 0.714 | 0.685 | 0.865 | 0.828 | 0.581 | 0.455 | 0.960 | 0.969 | 0.120 | 0.028 | 1043.49 | 0.448 | 0.541 | 0.789 | 0.780 | 0.562 | 0.558 | 0.787 | 0.842 | 0.124 | 0.125 |
| 1054.47 | 0.620 | 0.536 | 0.828 | 0.811 | 0.574 | 0.497 | 0.922 | 0.895 | 0.160 | 0.080 | 1054.47 | 0.404 | 0.457 | 0.576 | 0.590 | 0.561 | 0.559 | 0.548 | 0.580 | 0.376 | 0.360 |
| 1057.46 | 0.743 | 0.726 | 0.829 | 0.805 | 0.539 | 0.448 | 0.945 | 0.952 | 0.171 | 0.046 | 1057.46 | 0.462 | 0.588 | 0.778 | 0.746 | 0.541 | 0.539 | 0.833 | 0.839 | 0.140 | 0.162 |
| 1058.53 | 0.641 | 0.579 | 0.863 | 0.826 | 0.616 | 0.495 | 0.943 | 0.937 | 0.104 | 0.037 | 1058.53 | 0.564 | 0.677 | 0.785 | 0.771 | 0.577 | 0.566 | 0.805 | 0.833 | 0.090 | 0.101 |
| 1062.47 | 0.916 | 0.859 | 0.775 | 0.729 | 0.500 | 0.474 | 0.902 | 0.873 | 0.176 | 0.068 | 1062.47 | 0.533 | 0.532 | 0.626 | 0.621 | 0.544 | 0.555 | 0.632 | 0.616 | 0.309 | 0.316 |
| 1068.52 | 0.860 | 0.782 | 0.583 | 0.606 | 0.532 | 0.543 | 0.644 | 0.739 | 0.321 | 0.174 | 1068.52 | 0.890 | 0.869 | 0.634 | 0.640 | 0.455 | 0.480 | 0.803 | 0.753 | 0.292 | 0.270 |
| 1072.53 | 0.838 | 0.736 | 0.812 | 0.796 | 0.548 | 0.478 | 0.927 | 0.924 | 0.137 | 0.030 | 1072.53 | 0.891 | 0.911 | 0.768 | 0.772 | 0.483 | 0.490 | 0.916 | 0.913 | 0.120 | 0.106 |
| 1076.56 | 0.707 | 0.707 | 0.705 | 0.770 | 0.546 | 0.450 | 0.841 | 0.937 | 0.278 | 0.085 | 1076.56 | 0.440 | 0.483 | 0.563 | 0.539 | 0.567 | 0.562 | 0.573 | 0.592 | 0.369 | 0.389 |
| 1078.44 | 0.865 | 0.827 | 0.740 | 0.721 | 0.492 | 0.479 | 0.876 | 0.875 | 0.233 | 0.078 | 1078.44 | 0.752 | 0.838 | 0.827 | 0.807 | 0.481 | 0.478 | 0.931 | 0.936 | 0.090 | 0.098 |
| 1080.52 | 0.735 | 0.615 | 0.686 | 0.735 | 0.543 | 0.493 | 0.799 | 0.858 | 0.260 | 0.133 | 1080.52 | 0.628 | 0.589 | 0.562 | 0.560 | 0.533 | 0.543 | 0.614 | 0.569 | 0.365 | 0.377 |
| 1082.64 | 0.914 | 0.783 | 0.831 | 0.767 | 0.525 | 0.498 | 0.939 | 0.888 | 0.107 | 0.033 | 1082.64 | 0.825 | 0.846 | 0.840 | 0.834 | 0.502 | 0.531 | 0.925 | 0.917 | 0.047 | 0.029 |
| 1087.52 | 0.614 | 0.552 | 0.747 | 0.758 | 0.599 | 0.551 | 0.830 | 0.818 | 0.209 | 0.093 | 1087.52 | 0.389 | 0.467 | 0.702 | 0.700 | 0.612 | 0.604 | 0.625 | 0.683 | 0.199 | 0.202 |
| 1096.48 | 0.597 | 0.513 | 0.778 | 0.751 | 0.563 | 0.536 | 0.862 | 0.833 | 0.198 | 0.120 | 1096.48 | 0.471 | 0.523 | 0.670 | 0.687 | 0.590 | 0.592 | 0.635 | 0.674 | 0.234 | 0.214 |
| 1098.53 | 0.675 | 0.688 | 0.735 | 0.774 | 0.600 | 0.497 | 0.689 | 0.797 | 0.237 | 0.099 | 1098.53 | 0.527 | 0.632 | 0.714 | 0.691 | 0.535 | 0.532 | 0.662 | 0.640 | 0.235 | 0.256 |
| 1098.6 | 0.774 | 0.704 | 0.878 | 0.802 | 0.603 | 0.494 | 0.951 | 0.908 | 0.067 | 0.027 | 1098.6 | 0.580 | 0.709 | 0.869 | 0.850 | 0.537 | 0.546 | 0.903 | 0.917 | 0.030 | 0.030 |
| 1102.58 | 0.783 | 0.838 | 0.807 | 0.772 | 0.541 | 0.454 | 0.932 | 0.937 | 0.178 | 0.035 | 1102.58 | 0.570 | 0.702 | 0.857 | 0.826 | 0.490 | 0.483 | 0.928 | 0.945 | 0.082 | 0.101 |
| 1104.62 | 0.927 | 0.759 | 0.724 | 0.695 | 0.460 | 0.463 | 0.874 | 0.825 | 0.241 | 0.160 | 1104.62 | 0.668 | 0.703 | 0.608 | 0.602 | 0.513 | 0.515 | 0.716 | 0.698 | 0.313 | 0.313 |
| 1106.53 | 0.604 | 0.584 | 0.650 | 0.736 | 0.555 | 0.461 | 0.731 | 0.862 | 0.332 | 0.173 | 1106.53 | 0.444 | 0.502 | 0.651 | 0.644 | 0.559 | 0.564 | 0.620 | 0.656 | 0.289 | 0.283 |

**Supplemental Table 4 (2/3): Discriminating extracellular matrix imaging mass spectrometry features of each k-means segment by area under the receiver operating characteristic curve.** Segments are color coded as blue (B), green (G), purple (P), red (R),or yellow (Y) and indicated as N-glycan IMS performed without (-) or with (+) prior multiplexed immunofluorescence (MXIF) or picrosirius red staining (PSR). Values highlighted in green indicate top 10 discriminate features for each segment. Values highlighted in yellow indicate top 30 discriminate features for each segment. K-means segmentation analyses using Manhattan distance (k = 5) and receiver operating characteristic curves were performed in SCiLs v2026a Pro.

| m/z | MxIF- (B) | MxIF+ (B) | MxIF- (G) | MxIF+ (G) | MxIF- (P) | MxIF+ (P) | MxIF- (R) | MxIF+ (R) | MxIF- (Y) | MxIF+ (Y) | m/z | PSR- (B) | PSR+ (B) | PSR- (G) | PSR+ (G) | PSR- (P) | PSR+ (P) | PSR- (R) | PSR+ (R) | PSR- (Y) | PSR+ (Y) |
|---|---|---|---|---|---|---|---|---|---|---|---|---|---|---|---|---|---|---|---|---|---|
| 1111.48 | 0.815 | 0.652 | 0.676 | 0.685 | 0.514 | 0.534 | 0.787 | 0.772 | 0.274 | 0.153 | 1111.48 | 0.547 | 0.557 | 0.572 | 0.583 | 0.544 | 0.553 | 0.582 | 0.610 | 0.366 | 0.345 |
| 1113.53 | 0.692 | 0.615 | 0.801 | 0.761 | 0.568 | 0.518 | 0.909 | 0.866 | 0.157 | 0.082 | 1113.53 | 0.514 | 0.575 | 0.784 | 0.797 | 0.567 | 0.557 | 0.771 | 0.835 | 0.117 | 0.108 |
| 1114.53 | 0.604 | 0.572 | 0.846 | 0.800 | 0.604 | 0.520 | 0.929 | 0.876 | 0.120 | 0.060 | 1114.53 | 0.434 | 0.526 | 0.788 | 0.800 | 0.592 | 0.574 | 0.744 | 0.816 | 0.108 | 0.105 |
| 1115.55 | 0.895 | 0.821 | 0.890 | 0.794 | 0.545 | 0.456 | 0.984 | 0.962 | 0.048 | 0.010 | 1115.55 | 0.740 | 0.821 | 0.894 | 0.874 | 0.497 | 0.509 | 0.970 | 0.980 | 0.007 | 0.006 |
| 1116.51 | 0.504 | 0.478 | 0.765 | 0.770 | 0.593 | 0.483 | 0.783 | 0.841 | 0.221 | 0.168 | 1116.51 | 0.419 | 0.462 | 0.597 | 0.580 | 0.578 | 0.577 | 0.472 | 0.451 | 0.359 | 0.385 |
| 1118.48 | 0.907 | 0.800 | 0.849 | 0.766 | 0.534 | 0.491 | 0.945 | 0.894 | 0.086 | 0.032 | 1118.48 | 0.713 | 0.800 | 0.889 | 0.872 | 0.506 | 0.516 | 0.931 | 0.945 | 0.016 | 0.015 |
| 1120.59 | 0.887 | 0.755 | 0.787 | 0.747 | 0.500 | 0.487 | 0.918 | 0.871 | 0.175 | 0.076 | 1120.59 | 0.813 | 0.856 | 0.803 | 0.799 | 0.492 | 0.508 | 0.912 | 0.901 | 0.094 | 0.081 |
| 1122.56 | 0.942 | 0.928 | 0.842 | 0.743 | 0.504 | 0.461 | 0.960 | 0.926 | 0.105 | 0.024 | 1122.56 | 0.866 | 0.926 | 0.877 | 0.855 | 0.480 | 0.499 | 0.986 | 0.976 | 0.015 | 0.012 |
| 1125.54 | 0.964 | 0.931 | 0.823 | 0.732 | 0.496 | 0.463 | 0.957 | 0.930 | 0.114 | 0.027 | 1125.54 | 0.952 | 0.963 | 0.853 | 0.835 | 0.470 | 0.496 | 0.977 | 0.963 | 0.035 | 0.028 |
| 1128.54 | 0.809 | 0.749 | 0.922 | 0.830 | 0.581 | 0.446 | 0.990 | 0.987 | 0.038 | 0.007 | 1128.54 | 0.694 | 0.790 | 0.894 | 0.875 | 0.498 | 0.505 | 0.967 | 0.981 | 0.014 | 0.015 |
| 1131.54 | 0.737 | 0.685 | 0.898 | 0.827 | 0.595 | 0.472 | 0.977 | 0.948 | 0.062 | 0.020 | 1131.54 | 0.485 | 0.639 | 0.896 | 0.880 | 0.536 | 0.537 | 0.885 | 0.926 | 0.026 | 0.025 |
| 1133.53 | 0.707 | 0.677 | 0.824 | 0.786 | 0.575 | 0.492 | 0.901 | 0.888 | 0.145 | 0.059 | 1133.53 | 0.588 | 0.682 | 0.801 | 0.785 | 0.549 | 0.546 | 0.850 | 0.867 | 0.088 | 0.099 |
| 1137.55 | 0.983 | 0.914 | 0.737 | 0.667 | 0.453 | 0.492 | 0.890 | 0.835 | 0.213 | 0.094 | 1137.55 | 0.974 | 0.964 | 0.750 | 0.722 | 0.408 | 0.446 | 0.923 | 0.879 | 0.195 | 0.186 |
| 1139.6 | 0.950 | 0.949 | 0.851 | 0.733 | 0.527 | 0.470 | 0.954 | 0.919 | 0.074 | 0.019 | 1139.6 | 0.808 | 0.894 | 0.885 | 0.863 | 0.486 | 0.502 | 0.990 | 0.981 | 0.010 | 0.008 |
| 1139.66 | 0.762 | 0.579 | 0.697 | 0.738 | 0.560 | 0.586 | 0.730 | 0.740 | 0.245 | 0.096 | 1139.66 | 0.637 | 0.638 | 0.586 | 0.637 | 0.577 | 0.597 | 0.538 | 0.601 | 0.319 | 0.247 |
| 1140.46 | 0.895 | 0.738 | 0.710 | 0.687 | 0.464 | 0.509 | 0.853 | 0.788 | 0.251 | 0.141 | 1140.46 | 0.568 | 0.585 | 0.571 | 0.551 | 0.502 | 0.495 | 0.609 | 0.604 | 0.398 | 0.420 |
| 1140.53 | 0.662 | 0.661 | 0.701 | 0.690 | 0.596 | 0.524 | 0.779 | 0.811 | 0.239 | 0.140 | 1140.53 | 0.557 | 0.602 | 0.675 | 0.661 | 0.556 | 0.563 | 0.703 | 0.726 | 0.233 | 0.234 |
| 1142.52 | 0.786 | 0.799 | 0.892 | 0.804 | 0.581 | 0.447 | 0.980 | 0.976 | 0.075 | 0.015 | 1142.52 | 0.611 | 0.727 | 0.881 | 0.861 | 0.513 | 0.516 | 0.936 | 0.956 | 0.030 | 0.035 |
| 1146.53 | 0.551 | 0.573 | 0.889 | 0.812 | 0.622 | 0.516 | 0.948 | 0.881 | 0.081 | 0.052 | 1146.53 | 0.341 | 0.474 | 0.808 | 0.800 | 0.618 | 0.599 | 0.743 | 0.829 | 0.080 | 0.086 |
| 1147.52 | 0.859 | 0.831 | 0.859 | 0.763 | 0.550 | 0.471 | 0.964 | 0.925 | 0.087 | 0.032 | 1147.52 | 0.482 | 0.567 | 0.787 | 0.784 | 0.585 | 0.581 | 0.800 | 0.859 | 0.093 | 0.091 |
| 1150.53 | 0.834 | 0.751 | 0.836 | 0.794 | 0.533 | 0.452 | 0.961 | 0.948 | 0.130 | 0.044 | 1150.53 | 0.536 | 0.578 | 0.656 | 0.650 | 0.547 | 0.550 | 0.686 | 0.696 | 0.266 | 0.267 |
| 1151.61 | 0.346 | 0.403 | 0.607 | 0.549 | 0.709 | 0.677 | 0.509 | 0.451 | 0.277 | 0.331 | 1151.61 | 0.406 | 0.481 | 0.609 | 0.629 | 0.620 | 0.641 | 0.539 | 0.529 | 0.291 | 0.261 |
| 1154.63 | 0.869 | 0.778 | 0.686 | 0.697 | 0.508 | 0.519 | 0.795 | 0.794 | 0.258 | 0.107 | 1154.63 | 0.774 | 0.795 | 0.684 | 0.661 | 0.463 | 0.489 | 0.712 | 0.648 | 0.282 | 0.287 |
| 1156.44 | 0.860 | 0.732 | 0.703 | 0.639 | 0.475 | 0.529 | 0.830 | 0.742 | 0.258 | 0.180 | 1156.44 | 0.695 | 0.784 | 0.800 | 0.780 | 0.490 | 0.485 | 0.870 | 0.884 | 0.127 | 0.136 |
| 1157.53 | 0.872 | 0.780 | 0.814 | 0.776 | 0.554 | 0.498 | 0.906 | 0.862 | 0.117 | 0.037 | 1157.53 | 0.836 | 0.892 | 0.831 | 0.821 | 0.519 | 0.532 | 0.908 | 0.900 | 0.040 | 0.035 |
| 1161.58 | 0.929 | 0.921 | 0.757 | 0.734 | 0.489 | 0.456 | 0.912 | 0.913 | 0.190 | 0.045 | 1161.58 | 0.567 | 0.613 | 0.668 | 0.583 | 0.522 | 0.519 | 0.764 | 0.665 | 0.255 | 0.350 |
| 1163.58 | 0.838 | 0.812 | 0.898 | 0.800 | 0.551 | 0.453 | 0.974 | 0.956 | 0.069 | 0.014 | 1163.58 | 0.799 | 0.860 | 0.879 | 0.838 | 0.478 | 0.484 | 0.968 | 0.961 | 0.031 | 0.057 |
| 1164.5 | 0.881 | 0.842 | 0.752 | 0.733 | 0.508 | 0.455 | 0.899 | 0.922 | 0.201 | 0.069 | 1164.5 | 0.760 | 0.738 | 0.690 | 0.651 | 0.485 | 0.505 | 0.822 | 0.755 | 0.230 | 0.263 |
| 1171.56 | 0.820 | 0.784 | 0.867 | 0.805 | 0.555 | 0.451 | 0.974 | 0.966 | 0.096 | 0.018 | 1171.56 | 0.658 | 0.735 | 0.838 | 0.829 | 0.513 | 0.516 | 0.914 | 0.924 | 0.066 | 0.068 |
| 1172.52 | 0.947 | 0.846 | 0.687 | 0.680 | 0.504 | 0.514 | 0.735 | 0.770 | 0.240 | 0.113 | 1172.52 | 0.887 | 0.907 | 0.800 | 0.797 | 0.486 | 0.509 | 0.935 | 0.929 | 0.085 | 0.065 |
| 1173.52 | 0.871 | 0.743 | 0.885 | 0.787 | 0.560 | 0.498 | 0.971 | 0.910 | 0.056 | 0.020 | 1173.52 | 0.691 | 0.766 | 0.844 | 0.837 | 0.547 | 0.549 | 0.913 | 0.938 | 0.022 | 0.020 |
| 1177.55 | 0.878 | 0.907 | 0.759 | 0.725 | 0.518 | 0.463 | 0.910 | 0.921 | 0.188 | 0.045 | 1177.55 | 0.767 | 0.867 | 0.856 | 0.837 | 0.475 | 0.476 | 0.971 | 0.963 | 0.057 | 0.063 |
| 1179.57 | 0.888 | 0.907 | 0.847 | 0.756 | 0.517 | 0.454 | 0.968 | 0.939 | 0.115 | 0.022 | 1179.57 | 0.877 | 0.914 | 0.860 | 0.852 | 0.471 | 0.491 | 0.966 | 0.969 | 0.042 | 0.026 |
| 1184.62 | 0.637 | 0.665 | 0.833 | 0.788 | 0.560 | 0.430 | 0.943 | 0.947 | 0.164 | 0.101 | 1184.62 | 0.833 | 0.873 | 0.858 | 0.841 | 0.474 | 0.478 | 0.966 | 0.961 | 0.047 | 0.057 |
| 1185.56 | 0.805 | 0.781 | 0.772 | 0.765 | 0.554 | 0.465 | 0.892 | 0.909 | 0.175 | 0.060 | 1185.56 | 0.537 | 0.534 | 0.605 | 0.604 | 0.560 | 0.583 | 0.593 | 0.584 | 0.320 | 0.311 |
| 1188.57 | 0.687 | 0.665 | 0.912 | 0.832 | 0.597 | 0.459 | 0.980 | 0.960 | 0.064 | 0.031 | 1188.57 | 0.447 | 0.574 | 0.892 | 0.879 | 0.537 | 0.536 | 0.878 | 0.928 | 0.037 | 0.038 |
| 1193.54 | 0.766 | 0.708 | 0.696 | 0.705 | 0.532 | 0.476 | 0.841 | 0.875 | 0.254 | 0.134 | 1193.54 | 0.489 | 0.515 | 0.563 | 0.594 | 0.532 | 0.534 | 0.577 | 0.632 | 0.396 | 0.355 |
| 1195.5 | 0.886 | 0.729 | 0.771 | 0.740 | 0.505 | 0.508 | 0.894 | 0.839 | 0.184 | 0.082 | 1195.5 | 0.547 | 0.530 | 0.636 | 0.622 | 0.525 | 0.531 | 0.651 | 0.619 | 0.312 | 0.337 |
| 1197.64 | 0.777 | 0.808 | 0.762 | 0.772 | 0.514 | 0.427 | 0.900 | 0.942 | 0.237 | 0.071 | 1197.64 | 0.890 | 0.906 | 0.840 | 0.825 | 0.448 | 0.462 | 0.965 | 0.958 | 0.081 | 0.079 |
| 1201.55 | 0.844 | 0.853 | 0.754 | 0.725 | 0.537 | 0.462 | 0.885 | 0.899 | 0.192 | 0.074 | 1201.55 | 0.511 | 0.488 | 0.528 | 0.567 | 0.581 | 0.589 | 0.480 | 0.496 | 0.398 | 0.366 |
| 1203.59 | 0.812 | 0.834 | 0.790 | 0.760 | 0.520 | 0.428 | 0.932 | 0.947 | 0.192 | 0.068 | 1203.59 | 0.764 | 0.821 | 0.840 | 0.808 | 0.458 | 0.465 | 0.950 | 0.949 | 0.093 | 0.109 |
| 1205.58 | 0.734 | 0.756 | 0.859 | 0.801 | 0.558 | 0.438 | 0.961 | 0.959 | 0.137 | 0.047 | 1205.58 | 0.544 | 0.669 | 0.810 | 0.774 | 0.508 | 0.503 | 0.870 | 0.899 | 0.123 | 0.142 |
| 1210.55 | 0.629 | 0.602 | 0.776 | 0.733 | 0.556 | 0.500 | 0.898 | 0.817 | 0.197 | 0.148 | 1210.55 | 0.405 | 0.465 | 0.591 | 0.582 | 0.560 | 0.565 | 0.523 | 0.523 | 0.371 | 0.374 |
| 1211.47 | 0.796 | 0.672 | 0.709 | 0.702 | 0.504 | 0.528 | 0.826 | 0.801 | 0.266 | 0.126 | 1211.47 | 0.614 | 0.698 | 0.726 | 0.728 | 0.555 | 0.543 | 0.801 | 0.840 | 0.156 | 0.151 |
| 1211.62 | 0.909 | 0.902 | 0.899 | 0.775 | 0.554 | 0.451 | 0.984 | 0.950 | 0.033 | 0.008 | 1211.62 | 0.811 | 0.892 | 0.887 | 0.861 | 0.489 | 0.504 | 0.994 | 0.988 | 0.004 | 0.006 |
| 1217.52 | 0.818 | 0.804 | 0.664 | 0.663 | 0.514 | 0.482 | 0.789 | 0.837 | 0.294 | 0.145 | 1217.52 | 0.674 | 0.780 | 0.656 | 0.700 | 0.489 | 0.483 | 0.779 | 0.844 | 0.279 | 0.213 |
| 1218.56 | 0.562 | 0.548 | 0.835 | 0.822 | 0.615 | 0.499 | 0.915 | 0.899 | 0.144 | 0.064 | 1218.56 | 0.324 | 0.356 | 0.628 | 0.602 | 0.621 | 0.619 | 0.526 | 0.535 | 0.290 | 0.325 |
| 1224.58 | 0.794 | 0.823 | 0.694 | 0.723 | 0.549 | 0.470 | 0.789 | 0.883 | 0.261 | 0.084 | 1224.58 | 0.629 | 0.713 | 0.720 | 0.709 | 0.529 | 0.537 | 0.800 | 0.794 | 0.185 | 0.181 |
| 1226.62 | 0.687 | 0.706 | 0.862 | 0.788 | 0.542 | 0.428 | 0.978 | 0.957 | 0.136 | 0.086 | 1226.62 | 0.604 | 0.676 | 0.875 | 0.852 | 0.496 | 0.485 | 0.940 | 0.954 | 0.053 | 0.081 |
| 1233.61 | 0.924 | 0.899 | 0.851 | 0.743 | 0.511 | 0.458 | 0.969 | 0.907 | 0.100 | 0.045 | 1233.61 | 0.709 | 0.769 | 0.837 | 0.808 | 0.497 | 0.498 | 0.941 | 0.924 | 0.068 | 0.094 |
| 1241.54 | 0.627 | 0.577 | 0.690 | 0.720 | 0.560 | 0.498 | 0.795 | 0.827 | 0.276 | 0.165 | 1241.54 | 0.501 | 0.520 | 0.587 | 0.614 | 0.527 | 0.544 | 0.607 | 0.634 | 0.370 | 0.329 |
| 1242.59 | 0.970 | 0.943 | 0.839 | 0.734 | 0.506 | 0.477 | 0.938 | 0.895 | 0.095 | 0.022 | 1242.59 | 0.873 | 0.918 | 0.881 | 0.860 | 0.481 | 0.501 | 0.989 | 0.982 | 0.009 | 0.006 |
| 1249.57 | 0.883 | 0.894 | 0.822 | 0.750 | 0.529 | 0.470 | 0.950 | 0.922 | 0.127 | 0.022 | 1249.57 | 0.809 | 0.893 | 0.867 | 0.845 | 0.490 | 0.498 | 0.977 | 0.972 | 0.025 | 0.028 |
| 1257.59 | 0.936 | 0.944 | 0.831 | 0.727 | 0.517 | 0.486 | 0.935 | 0.872 | 0.109 | 0.028 | 1257.59 | 0.745 | 0.839 | 0.896 | 0.875 | 0.490 | 0.503 | 0.983 | 0.984 | 0.008 | 0.007 |
| 1264.57 | 0.970 | 0.925 | 0.760 | 0.693 | 0.469 | 0.476 | 0.913 | 0.856 | 0.187 | 0.078 | 1264.57 | 0.740 | 0.772 | 0.756 | 0.729 | 0.495 | 0.501 | 0.899 | 0.876 | 0.146 | 0.167 |
| 1271.56 | 0.874 | 0.833 | 0.676 | 0.660 | 0.515 | 0.503 | 0.810 | 0.791 | 0.258 | 0.135 | 1271.56 | 0.554 | 0.554 | 0.611 | 0.611 | 0.539 | 0.554 | 0.657 | 0.625 | 0.318 | 0.318 |
| 1271.63 | 0.731 | 0.779 | 0.611 | 0.698 | 0.553 | 0.475 | 0.639 | 0.820 | 0.361 | 0.141 | 1271.63 | 0.529 | 0.598 | 0.679 | 0.645 | 0.537 | 0.533 | 0.713 | 0.703 | 0.249 | 0.281 |
| 1279.58 | 0.901 | 0.890 | 0.685 | 0.661 | 0.511 | 0.501 | 0.810 | 0.790 | 0.248 | 0.117 | 1279.58 | 0.515 | 0.530 | 0.608 | 0.614 | 0.543 | 0.555 | 0.618 | 0.627 | 0.332 | 0.318 |
| 1280.55 | 0.926 | 0.904 | 0.714 | 0.678 | 0.492 | 0.498 | 0.841 | 0.831 | 0.229 | 0.085 | 1280.55 | 0.860 | 0.897 | 0.831 | 0.782 | 0.467 | 0.488 | 0.960 | 0.922 | 0.076 | 0.101 |
| 1280.7 | 0.701 | 0.732 | 0.792 | 0.791 | 0.576 | 0.456 | 0.930 | 0.958 | 0.185 | 0.045 | 1280.7 | 0.510 | 0.567 | 0.653 | 0.620 | 0.540 | 0.532 | 0.717 | 0.710 | 0.271 | 0.306 |
| 1283.64 | 0.836 | 0.876 | 0.829 | 0.748 | 0.581 | 0.489 | 0.920 | 0.888 | 0.101 | 0.024 | 1283.64 | 0.612 | 0.734 | 0.894 | 0.869 | 0.510 | 0.523 | 0.954 | 0.961 | 0.018 | 0.020 |
| 1286.67 | 0.885 | 0.882 | 0.725 | 0.726 | 0.503 | 0.438 | 0.888 | 0.915 | 0.223 | 0.081 | 1286.67 | 0.805 | 0.745 | 0.751 | 0.678 | 0.428 | 0.435 | 0.900 | 0.842 | 0.206 | 0.283 |
| 1295.55 | 0.845 | 0.829 | 0.625 | 0.622 | 0.497 | 0.509 | 0.721 | 0.744 | 0.335 | 0.179 | 1295.55 | 0.631 | 0.726 | 0.759 | 0.751 | 0.496 | 0.500 | 0.858 | 0.859 | 0.170 | 0.163 |
| 1302.52 | 0.896 | 0.825 | 0.596 | 0.587 | 0.459 | 0.495 | 0.737 | 0.718 | 0.359 | 0.234 | 1302.52 | 0.658 | 0.667 | 0.589 | 0.573 | 0.470 | 0.468 | 0.708 | 0.700 | 0.375 | 0.388 |
| 1302.67 | 0.958 | 0.942 | 0.846 | 0.732 | 0.506 | 0.465 | 0.957 | 0.925 | 0.093 | 0.024 | 1302.67 | 0.955 | 0.969 | 0.852 | 0.834 | 0.463 | 0.493 | 0.975 | 0.959 | 0.043 | 0.030 |
| 1305.63 | 0.808 | 0.839 | 0.739 | 0.714 | 0.552 | 0.475 | 0.846 | 0.867 | 0.211 | 0.087 | 1305.63 | 0.657 | 0.751 | 0.792 | 0.770 | 0.473 | 0.475 | 0.877 | 0.891 | 0.155 | 0.158 |
| 1306.62 | 0.903 | 0.852 | 0.784 | 0.734 | 0.505 | 0.463 | 0.929 | 0.901 | 0.164 | 0.066 | 1306.62 | 0.557 | 0.599 | 0.664 | 0.648 | 0.550 | 0.543 | 0.710 | 0.713 | 0.246 | 0.266 |
| 1321.58 | 0.886 | 0.792 | 0.737 | 0.692 | 0.487 | 0.488 | 0.878 | 0.853 | 0.217 | 0.113 | 1321.58 | 0.589 | 0.674 | 0.763 | 0.741 | 0.495 | 0.500 | 0.861 | 0.867 | 0.172 | 0.178 |
| 1322.58 | 0.899 | 0.835 | 0.794 | 0.734 | 0.502 | 0.469 | 0.936 | 0.914 | 0.157 | 0.059 | 1322.58 | 0.772 | 0.846 | 0.840 | 0.821 | 0.471 | 0.480 | 0.955 | 0.957 | 0.078 | 0.083 |
| 1323.79 | 0.792 | 0.641 | 0.624 | 0.691 | 0.544 | 0.604 | 0.633 | 0.683 | 0.330 | 0.117 | 1323.79 | 0.571 | 0.699 | 0.574 | 0.585 | 0.537 | 0.552 | 0.594 | 0.587 | 0.365 | 0.322 |
| 1324.64 | 0.769 | 0.768 | 0.579 | 0.599 | 0.624 | 0.566 | 0.607 | 0.725 | 0.297 | 0.167 | 1324.64 | 0.295 | 0.270 | 0.435 | 0.475 | 0.589 | 0.609 | 0.311 | 0.334 | 0.547 | 0.505 |
| 1328.63 | 0.758 | 0.735 | 0.765 | 0.765 | 0.567 | 0.491 | 0.875 | 0.894 | 0.195 | 0.055 | 1328.63 | 0.493 | 0.582 | 0.786 | 0.781 | 0.563 | 0.561 | 0.797 | 0.846 | 0.116 | 0.112 |
| 1336.67 | 0.686 | 0.708 | 0.705 | 0.732 | 0.562 | 0.475 | 0.812 | 0.854 | 0.265 | 0.123 | 1336.67 | 0.475 | 0.532 | 0.679 | 0.670 | 0.548 | 0.532 | 0.706 | 0.739 | 0.249 | 0.266 |
| 1340.62 | 0.885 | 0.914 | 0.734 | 0.697 | 0.530 | 0.477 | 0.882 | 0.895 | 0.194 | 0.060 | 1340.62 | 0.941 | 0.961 | 0.809 | 0.789 | 0.447 | 0.469 | 0.961 | 0.942 | 0.101 | 0.095 |
| 1343.67 | 0.977 | 0.960 | 0.764 | 0.688 | 0.472 | 0.487 | 0.895 | 0.870 | 0.181 | 0.053 | 1343.67 | 0.966 | 0.975 | 0.841 | 0.826 | 0.453 | 0.486 | 0.968 | 0.949 | 0.063 | 0.045 |
| 1346.63 | 0.574 | 0.457 | 0.517 | 0.509 | 0.559 | 0.585 | 0.493 | 0.468 | 0.428 | 0.433 | 1346.63 | 0.404 | 0.413 | 0.477 | 0.493 | 0.506 | 0.509 | 0.429 | 0.447 | 0.546 | 0.528 |
| 1350.65 | 0.934 | 0.915 | 0.593 | 0.587 | 0.500 | 0.515 | 0.686 | 0.746 | 0.324 | 0.170 | 1350.65 | 0.980 | 0.974 | 0.721 | 0.719 | 0.442 | 0.462 | 0.919 | 0.893 | 0.187 | 0.167 |
| 1352.66 | 0.805 | 0.815 | 0.697 | 0.699 | 0.546 | 0.489 | 0.799 | 0.840 | 0.249 | 0.105 | 1352.66 | 0.861 | 0.852 | 0.724 | 0.723 | 0.494 | 0.507 | 0.870 | 0.869 | 0.164 | 0.152 |
| 1359.68 | 0.951 | 0.905 | 0.625 | 0.665 | 0.487 | 0.493 | 0.777 | 0.863 | 0.292 | 0.086 | 1359.68 | 0.951 | 0.952 | 0.725 | 0.710 | 0.458 | 0.470 | 0.924 | 0.904 | 0.171 | 0.169 |
| 1361.68 | 0.823 | 0.824 | 0.704 | 0.704 | 0.510 | 0.452 | 0.894 | 0.901 | 0.250 | 0.110 | 1361.68 | 0.784 | 0.815 | 0.656 | 0.614 | 0.501 | 0.495 | 0.824 | 0.777 | 0.238 | 0.281 |
| 1365.66 | 0.963 | 0.926 | 0.634 | 0.631 | 0.458 | 0.478 | 0.775 | 0.805 | 0.308 | 0.147 | 1365.66 | 0.744 | 0.702 | 0.510 | 0.494 | 0.480 | 0.485 | 0.639 | 0.545 | 0.438 | 0.468 |
| 1381.64 | 0.929 | 0.908 | 0.661 | 0.638 | 0.503 | 0.504 | 0.780 | 0.819 | 0.263 | 0.114 | 1381.64 | 0.945 | 0.962 | 0.768 | 0.756 | 0.441 | 0.465 | 0.947 | 0.917 | 0.146 | 0.131 |
| 1384.69 | 0.805 | 0.807 | 0.785 | 0.774 | 0.532 | 0.442 | 0.948 | 0.963 | 0.183 | 0.046 | 1384.69 | 0.720 | 0.828 | 0.821 | 0.801 | 0.472 | 0.480 | 0.947 | 0.943 | 0.104 | 0.100 |
| 1386.69 | 0.882 | 0.903 | 0.798 | 0.730 | 0.536 | 0.483 | 0.930 | 0.902 | 0.141 | 0.030 | 1386.69 | 0.880 | 0.933 | 0.829 | 0.815 | 0.505 | 0.517 | 0.949 | 0.935 | 0.039 | 0.037 |
| 1399.65 | 0.822 | 0.792 | 0.924 | 0.816 | 0.580 | 0.442 | 0.992 | 0.991 | 0.034 | 0.005 | 1399.65 | 0.759 | 0.840 | 0.894 | 0.872 | 0.490 | 0.503 | 0.986 | 0.989 | 0.007 | 0.008 |
| 1408.68 | 0.854 | 0.875 | 0.639 | 0.670 | 0.526 | 0.492 | 0.750 | 0.826 | 0.298 | 0.109 | 1408.68 | 0.560 | 0.592 | 0.613 | 0.599 | 0.528 | 0.517 | 0.654 | 0.665 | 0.326 | 0.344 |
| 1410.68 | 0.895 | 0.911 | 0.768 | 0.722 | 0.512 | 0.461 | 0.931 | 0.921 | 0.176 | 0.048 | 1410.68 | 0.943 | 0.949 | 0.828 | 0.804 | 0.456 | 0.478 | 0.966 | 0.949 | 0.075 | 0.076 |
| 1421.64 | 0.888 | 0.836 | 0.832 | 0.774 | 0.508 | 0.441 | 0.968 | 0.954 | 0.133 | 0.040 | 1421.64 | 0.881 | 0.886 | 0.736 | 0.719 | 0.455 | 0.484 | 0.901 | 0.858 | 0.181 | 0.174 |
| 1421.8 | 0.897 | 0.875 | 0.678 | 0.725 | 0.502 | 0.493 | 0.734 | 0.863 | 0.278 | 0.049 | 1421.8 | 0.885 | 0.939 | 0.685 | 0.704 | 0.463 | 0.480 | 0.786 | 0.817 | 0.246 | 0.189 |
| 1426.68 | 0.976 | 0.941 | 0.677 | 0.652 | 0.484 | 0.494 | 0.825 | 0.847 | 0.240 | 0.090 | 1426.68 | 0.978 | 0.976 | 0.782 | 0.775 | 0.422 | 0.458 | 0.940 | 0.916 | 0.150 | 0.121 |
| 1430.66 | 0.680 | 0.651 | 0.740 | 0.742 | 0.560 | 0.491 | 0.860 | 0.871 | 0.212 | 0.110 | 1430.66 | 0.581 | 0.712 | 0.733 | 0.760 | 0.565 | 0.537 | 0.718 | 0.837 | 0.167 | 0.129 |
| 1437.61 | 0.696 | 0.689 | 0.797 | 0.784 | 0.528 | 0.435 | 0.940 | 0.961 | 0.196 | 0.085 | 1437.61 | 0.522 | 0.641 | 0.755 | 0.726 | 0.500 | 0.488 | 0.803 | 0.842 | 0.199 | 0.215 |
| 1437.65 | 0.866 | 0.868 | 0.554 | 0.486 | 0.527 | 0.557 | 0.557 | 0.516 | 0.360 | 0.320 | 1437.65 | 0.913 | 0.945 | 0.695 | 0.744 | 0.452 | 0.460 | 0.885 | 0.922 | 0.218 | 0.147 |
| 1438.72 | 0.956 | 0.961 | 0.741 | 0.690 | 0.486 | 0.480 | 0.890 | 0.876 | 0.194 | 0.056 | 1438.72 | 0.960 | 0.970 | 0.847 | 0.828 | 0.459 | 0.478 | 0.974 | 0.957 | 0.051 | 0.050 |
| 1441.61 | 0.771 | 0.775 | 0.847 | 0.798 | 0.529 | 0.437 | 0.957 | 0.952 | 0.158 | 0.047 | 1441.61 | 0.559 | 0.677 | 0.851 | 0.827 | 0.484 | 0.493 | 0.901 | 0.918 | 0.101 | 0.104 |
| 1446.69 | 0.909 | 0.924 | 0.617 | 0.622 | 0.506 | 0.501 | 0.715 | 0.771 | 0.311 | 0.144 | 1446.69 | 0.960 | 0.967 | 0.711 | 0.754 | 0.442 | 0.453 | 0.898 | 0.904 | 0.204 | 0.147 |
| 1448.66 | 0.929 | 0.868 | 0.632 | 0.622 | 0.497 | 0.505 | 0.760 | 0.795 | 0.285 | 0.149 | 1448.66 | 0.830 | 0.843 | 0.634 | 0.646 | 0.486 | 0.498 | 0.764 | 0.761 | 0.281 | 0.251 |
| 1458.72 | 0.905 | 0.917 | 0.693 | 0.713 | 0.477 | 0.461 | 0.867 | 0.894 | 0.263 | 0.065 | 1458.72 | 0.882 | 0.922 | 0.808 | 0.817 | 0.451 | 0.464 | 0.945 | 0.947 | 0.112 | 0.084 |
| 1460.7 | 0.917 | 0.923 | 0.744 | 0.720 | 0.513 | 0.477 | 0.900 | 0.901 | 0.184 | 0.038 | 1460.7 | 0.815 | 0.869 | 0.797 | 0.810 | 0.503 | 0.507 | 0.933 | 0.940 | 0.082 | 0.061 |
| 1463.59 | 0.767 | 0.742 | 0.695 | 0.697 | 0.504 | 0.462 | 0.832 | 0.847 | 0.287 | 0.155 | 1463.59 | 0.545 | 0.572 | 0.532 | 0.516 | 0.485 | 0.489 | 0.554 | 0.584 | 0.466 | 0.461 |
| 1472.7 | 0.740 | 0.763 | 0.838 | 0.794 | 0.565 | 0.449 | 0.941 | 0.945 | 0.147 | 0.044 | 1472.7 | 0.647 | 0.757 | 0.841 | 0.819 | 0.500 | 0.509 | 0.905 | 0.917 | 0.080 | 0.080 |
| 1480.76 | 0.841 | 0.893 | 0.844 | 0.746 | 0.555 | 0.477 | 0.958 | 0.914 | 0.106 | 0.022 | 1480.76 | 0.733 | 0.829 | 0.858 | 0.834 | 0.516 | 0.523 | 0.955 | 0.953 | 0.023 | 0.031 |
| 1482.69 | 0.930 | 0.891 | 0.632 | 0.625 | 0.511 | 0.514 | 0.754 | 0.781 | 0.277 | 0.135 | 1482.69 | 0.970 | 0.961 | 0.669 | 0.685 | 0.431 | 0.462 | 0.885 | 0.860 | 0.252 | 0.206 |
| 1483.73 | 0.920 | 0.900 | 0.886 | 0.763 | 0.529 | 0.453 | 0.982 | 0.950 | 0.059 | 0.016 | 1483.73 | 0.912 | 0.950 | 0.865 | 0.842 | 0.473 | 0.498 | 0.980 | 0.965 | 0.027 | 0.022 |
| 1488.68 | 0.714 | 0.717 | 0.707 | 0.732 | 0.567 | 0.495 | 0.780 | 0.831 | 0.253 | 0.109 | 1488.68 | 0.568 | 0.576 | 0.702 | 0.634 | 0.528 | 0.555 | 0.725 | 0.631 | 0.230 | 0.293 |
| 1494.69 | 0.776 | 0.778 | 0.714 | 0.717 | 0.546 | 0.477 | 0.834 | 0.862 | 0.239 | 0.107 | 1494.69 | 0.689 | 0.715 | 0.655 | 0.654 | 0.516 | 0.522 | 0.739 | 0.721 | 0.261 | 0.258 |
| 1495.73 | 0.923 | 0.903 | 0.627 | 0.541 | 0.480 | 0.513 | 0.732 | 0.654 | 0.310 | 0.252 | 1495.73 | 0.968 | 0.971 | 0.736 | 0.716 | 0.416 | 0.431 | 0.896 | 0.870 | 0.207 | 0.206 |
| 1497.76 | 0.899 | 0.905 | 0.744 | 0.719 | 0.477 | 0.442 | 0.914 | 0.921 | 0.225 | 0.072 | 1497.76 | 0.932 | 0.933 | 0.737 | 0.701 | 0.409 | 0.426 | 0.923 | 0.891 | 0.212 | 0.225 |
| 1502.74 | 0.826 | 0.858 | 0.676 | 0.689 | 0.533 | 0.493 | 0.809 | 0.844 | 0.271 | 0.092 | 1502.74 | 0.548 | 0.562 | 0.559 | 0.537 | 0.521 | 0.528 | 0.601 | 0.579 | 0.396 | 0.412 |
| 1504.68 | 0.872 | 0.840 | 0.653 | 0.674 | 0.519 | 0.494 | 0.771 | 0.820 | 0.279 | 0.119 | 1504.68 | 0.708 | 0.737 | 0.666 | 0.659 | 0.507 | 0.518 | 0.780 | 0.753 | 0.247 | 0.245 |
| 1505.71 | 0.929 | 0.893 | 0.794 | 0.724 | 0.476 | 0.453 | 0.943 | 0.897 | 0.169 | 0.071 | 1505.71 | 0.374 | 0.343 | 0.336 | 0.341 | 0.504 | 0.499 | 0.325 | 0.314 | 0.702 | 0.706 |
| 1508.76 | 0.937 | 0.880 | 0.533 | 0.528 | 0.474 | 0.533 | 0.569 | 0.595 | 0.397 | 0.275 | 1508.76 | 0.978 | 0.978 | 0.767 | 0.769 | 0.410 | 0.432 | 0.928 | 0.899 | 0.178 | 0.154 |
| 1518.71 | 0.844 | 0.852 | 0.666 | 0.659 | 0.532 | 0.513 | 0.783 | 0.816 | 0.271 | 0.108 | 1518.71 | 0.717 | 0.785 | 0.727 | 0.705 | 0.512 | 0.516 | 0.864 | 0.860 | 0.166 | 0.173 |
| 1521.69 | 0.883 | 0.890 | 0.781 | 0.721 | 0.512 | 0.458 | 0.934 | 0.920 | 0.170 | 0.060 | 1521.69 | 0.903 | 0.944 | 0.821 | 0.792 | 0.445 | 0.462 | 0.962 | 0.944 | 0.098 | 0.102 |
| 1521.81 | 0.953 | 0.961 | 0.699 | 0.644 | 0.490 | 0.525 | 0.826 | 0.803 | 0.232 | 0.076 | 1521.81 | 0.924 | 0.955 | 0.835 | 0.809 | 0.484 | 0.496 | 0.962 | 0.940 | 0.045 | 0.055 |
| 1528.72 | 0.783 | 0.827 | 0.763 | 0.758 | 0.556 | 0.470 | 0.884 | 0.893 | 0.198 | 0.052 | 1528.72 | 0.546 | 0.628 | 0.757 | 0.726 | 0.536 | 0.538 | 0.829 | 0.832 | 0.151 | 0.172 |

**Supplemental Table 4 (3/3): Discriminating extracellular matrix imaging mass spectrometry features of each k-means segment by area under the receiver operating characteristic curve.** Segments are color coded as blue (B), green (G), purple (P), red (R),or yellow (Y) and indicated as N-glycan IMS performed without (-) or with (+) prior multiplexed immunofluorescence (MXIF) or picrosirius red staining (PSR). Values highlighted in green indicate top 10 discriminate features for each segment. Values highlighted in yellow indicate top 30 discriminate features for each segment. K-means segmentation analyses using Manhattan distance (k = 5) and receiver operating characteristic curves were performed in SCiLs v2026a Pro.

| m/z | MxIF- (B) | MxIF+ (B) | MxIF- (G) | MxIF+ (G) | MxIF- (P) | MxIF+ (P) | MxIF- (R) | MxIF+ (R) | MxIF- (Y) | MxIF+ (Y) | m/z | PSR- (B) | PSR+ (B) | PSR- (G) | PSR+ (G) | PSR- (P) | PSR+ (P) | PSR- (R) | PSR+ (R) | PSR- (Y) | PSR+ (Y) |
|---|---|---|---|---|---|---|---|---|---|---|---|---|---|---|---|---|---|---|---|---|---|
| **1540.81** | 0.813 | 0.845 | 0.883 | 0.789 | 0.552 | 0.448 | 0.974 | 0.961 | 0.095 | 0.014 | **1540.81** | 0.790 | 0.869 | 0.877 | 0.854 | 0.479 | 0.482 | 0.974 | 0.969 | 0.031 | 0.041 |
| **1543.68** | 0.836 | 0.799 | 0.667 | 0.645 | 0.509 | 0.501 | 0.809 | 0.782 | 0.275 | 0.164 | **1543.68** | 0.396 | 0.419 | 0.502 | 0.504 | 0.551 | 0.566 | 0.395 | 0.409 | 0.490 | 0.474 |
| **1562.79** | 0.806 | 0.837 | 0.705 | 0.743 | 0.521 | 0.444 | 0.850 | 0.913 | 0.264 | 0.078 | **1562.79** | 0.515 | 0.512 | 0.531 | 0.512 | 0.515 | 0.522 | 0.551 | 0.516 | 0.443 | 0.463 |
| **1578.76** | 0.774 | 0.799 | 0.683 | 0.722 | 0.548 | 0.459 | 0.803 | 0.902 | 0.273 | 0.097 | **1578.76** | 0.739 | 0.797 | 0.745 | 0.710 | 0.477 | 0.480 | 0.885 | 0.860 | 0.177 | 0.200 |
| **1588.81** | 0.738 | 0.786 | 0.901 | 0.796 | 0.585 | 0.458 | 0.975 | 0.947 | 0.085 | 0.025 | **1588.81** | 0.607 | 0.725 | 0.863 | 0.837 | 0.483 | 0.482 | 0.935 | 0.946 | 0.076 | 0.089 |
| **1609.81** | 0.951 | 0.923 | 0.627 | 0.623 | 0.479 | 0.495 | 0.767 | 0.811 | 0.299 | 0.133 | **1609.81** | 0.977 | 0.979 | 0.774 | 0.735 | 0.429 | 0.452 | 0.950 | 0.909 | 0.147 | 0.159 |
| **1610.79** | 0.893 | 0.870 | 0.766 | 0.744 | 0.502 | 0.454 | 0.929 | 0.927 | 0.186 | 0.049 | **1610.79** | 0.974 | 0.978 | 0.772 | 0.756 | 0.446 | 0.472 | 0.947 | 0.912 | 0.133 | 0.123 |
| **1622.83** | 0.830 | 0.818 | 0.610 | 0.606 | 0.513 | 0.499 | 0.722 | 0.761 | 0.323 | 0.199 | **1622.83** | 0.908 | 0.949 | 0.794 | 0.762 | 0.453 | 0.461 | 0.950 | 0.920 | 0.117 | 0.133 |
| **1622.96** | 0.304 | 0.441 | 0.570 | 0.500 | 0.544 | 0.525 | 0.561 | 0.461 | 0.433 | 0.510 | **1622.96** | 0.336 | 0.421 | 0.545 | 0.513 | 0.533 | 0.534 | 0.494 | 0.474 | 0.455 | 0.479 |
| **1626.75** | 0.739 | 0.769 | 0.751 | 0.751 | 0.557 | 0.451 | 0.896 | 0.927 | 0.215 | 0.082 | **1626.75** | 0.606 | 0.671 | 0.677 | 0.660 | 0.510 | 0.519 | 0.765 | 0.762 | 0.254 | 0.253 |
| **1629.74** | 0.892 | 0.903 | 0.780 | 0.740 | 0.530 | 0.478 | 0.921 | 0.894 | 0.154 | 0.031 | **1629.74** | 0.851 | 0.917 | 0.834 | 0.815 | 0.493 | 0.508 | 0.959 | 0.941 | 0.048 | 0.047 |
| **1639.82** | 0.766 | 0.837 | 0.535 | 0.655 | 0.555 | 0.476 | 0.524 | 0.796 | 0.418 | 0.163 | **1639.82** | 0.547 | 0.558 | 0.530 | 0.531 | 0.516 | 0.520 | 0.552 | 0.535 | 0.438 | 0.437 |
| **1641.74** | 0.675 | 0.664 | 0.696 | 0.686 | 0.560 | 0.506 | 0.811 | 0.806 | 0.257 | 0.162 | **1641.74** | 0.615 | 0.618 | 0.625 | 0.611 | 0.516 | 0.539 | 0.683 | 0.653 | 0.312 | 0.312 |
| **1648.75** | 0.787 | 0.771 | 0.645 | 0.648 | 0.541 | 0.502 | 0.758 | 0.800 | 0.291 | 0.163 | **1648.75** | 0.879 | 0.871 | 0.615 | 0.611 | 0.461 | 0.486 | 0.790 | 0.742 | 0.308 | 0.290 |
| **1651.73** | 0.872 | 0.883 | 0.671 | 0.687 | 0.527 | 0.490 | 0.791 | 0.835 | 0.261 | 0.091 | **1651.73** | 0.603 | 0.609 | 0.553 | 0.546 | 0.512 | 0.525 | 0.602 | 0.557 | 0.401 | 0.404 |
| **1654.98** | 0.375 | 0.414 | 0.507 | 0.520 | 0.609 | 0.623 | 0.401 | 0.448 | 0.451 | 0.408 | **1654.98** | 0.300 | 0.376 | 0.528 | 0.493 | 0.577 | 0.581 | 0.387 | 0.402 | 0.459 | 0.479 |
| **1656.77** | 0.791 | 0.835 | 0.861 | 0.768 | 0.547 | 0.444 | 0.974 | 0.945 | 0.115 | 0.046 | **1656.77** | 0.752 | 0.845 | 0.878 | 0.851 | 0.470 | 0.481 | 0.968 | 0.964 | 0.046 | 0.050 |
| **1667.7** | 0.841 | 0.836 | 0.641 | 0.663 | 0.514 | 0.497 | 0.745 | 0.814 | 0.313 | 0.128 | **1667.7** | 0.770 | 0.818 | 0.684 | 0.666 | 0.482 | 0.491 | 0.834 | 0.797 | 0.233 | 0.238 |
| **1670.79** | 0.765 | 0.780 | 0.712 | 0.680 | 0.545 | 0.481 | 0.852 | 0.851 | 0.234 | 0.134 | **1670.79** | 0.889 | 0.875 | 0.697 | 0.678 | 0.458 | 0.477 | 0.875 | 0.850 | 0.217 | 0.216 |
| **1675.79** | 0.841 | 0.885 | 0.772 | 0.753 | 0.528 | 0.446 | 0.917 | 0.942 | 0.187 | 0.039 | **1675.79** | 0.849 | 0.911 | 0.805 | 0.794 | 0.452 | 0.468 | 0.951 | 0.942 | 0.116 | 0.102 |
| **1681.82** | 0.934 | 0.924 | 0.736 | 0.682 | 0.498 | 0.478 | 0.895 | 0.892 | 0.195 | 0.070 | **1681.82** | 0.937 | 0.943 | 0.853 | 0.834 | 0.456 | 0.479 | 0.977 | 0.969 | 0.051 | 0.046 |
| **1692.82** | 0.843 | 0.906 | 0.745 | 0.710 | 0.521 | 0.467 | 0.882 | 0.883 | 0.216 | 0.069 | **1692.82** | 0.825 | 0.886 | 0.840 | 0.814 | 0.462 | 0.473 | 0.954 | 0.947 | 0.079 | 0.084 |
| **1697.78** | 0.777 | 0.806 | 0.639 | 0.674 | 0.563 | 0.484 | 0.722 | 0.846 | 0.293 | 0.129 | **1697.78** | 0.508 | 0.604 | 0.675 | 0.717 | 0.542 | 0.539 | 0.632 | 0.676 | 0.272 | 0.225 |
| **1697.87** | 0.908 | 0.881 | 0.615 | 0.631 | 0.481 | 0.490 | 0.714 | 0.794 | 0.328 | 0.153 | **1697.87** | 0.795 | 0.716 | 0.423 | 0.427 | 0.463 | 0.471 | 0.552 | 0.544 | 0.545 | 0.532 |
| **1703.81** | 0.876 | 0.848 | 0.627 | 0.619 | 0.511 | 0.499 | 0.763 | 0.781 | 0.294 | 0.170 | **1703.81** | 0.668 | 0.663 | 0.595 | 0.580 | 0.500 | 0.511 | 0.694 | 0.644 | 0.343 | 0.357 |
| **1705.78** | 0.850 | 0.808 | 0.689 | 0.671 | 0.517 | 0.491 | 0.816 | 0.837 | 0.250 | 0.127 | **1705.78** | 0.757 | 0.764 | 0.600 | 0.596 | 0.496 | 0.507 | 0.639 | 0.654 | 0.343 | 0.325 |
| **1708.89** | 0.790 | 0.851 | 0.667 | 0.734 | 0.520 | 0.439 | 0.821 | 0.932 | 0.304 | 0.077 | **1708.89** | 0.512 | 0.511 | 0.575 | 0.557 | 0.508 | 0.505 | 0.680 | 0.647 | 0.379 | 0.409 |
| **1731.78** | 0.838 | 0.845 | 0.724 | 0.717 | 0.532 | 0.471 | 0.857 | 0.894 | 0.223 | 0.076 | **1731.78** | 0.836 | 0.896 | 0.784 | 0.761 | 0.451 | 0.469 | 0.945 | 0.924 | 0.139 | 0.134 |
| **1767.94** | 0.935 | 0.927 | 0.777 | 0.715 | 0.494 | 0.472 | 0.924 | 0.910 | 0.167 | 0.041 | **1767.94** | 0.946 | 0.968 | 0.793 | 0.753 | 0.447 | 0.470 | 0.944 | 0.901 | 0.119 | 0.133 |
| **1772.8** | 0.869 | 0.892 | 0.793 | 0.716 | 0.504 | 0.457 | 0.948 | 0.902 | 0.172 | 0.072 | **1772.8** | 0.901 | 0.922 | 0.848 | 0.830 | 0.451 | 0.465 | 0.954 | 0.946 | 0.072 | 0.074 |
| **1781.92** | 0.939 | 0.947 | 0.594 | 0.618 | 0.487 | 0.518 | 0.684 | 0.752 | 0.336 | 0.130 | **1781.92** | 0.967 | 0.970 | 0.730 | 0.678 | 0.426 | 0.452 | 0.920 | 0.860 | 0.197 | 0.219 |
| **1794.8** | 0.810 | 0.811 | 0.653 | 0.628 | 0.524 | 0.496 | 0.800 | 0.778 | 0.285 | 0.180 | **1794.8** | 0.647 | 0.620 | 0.547 | 0.537 | 0.484 | 0.511 | 0.612 | 0.553 | 0.425 | 0.423 |
| **1822.83** | 0.711 | 0.726 | 0.608 | 0.613 | 0.565 | 0.520 | 0.675 | 0.757 | 0.334 | 0.205 | **1822.83** | 0.674 | 0.683 | 0.595 | 0.578 | 0.490 | 0.517 | 0.668 | 0.630 | 0.359 | 0.353 |
| **1829.91** | 0.855 | 0.908 | 0.841 | 0.734 | 0.533 | 0.481 | 0.939 | 0.879 | 0.126 | 0.037 | **1829.91** | 0.797 | 0.884 | 0.861 | 0.841 | 0.484 | 0.498 | 0.948 | 0.940 | 0.044 | 0.042 |
| **1847.08** | 0.239 | 0.400 | 0.570 | 0.430 | 0.566 | 0.473 | 0.476 | 0.395 | 0.440 | 0.662 | **1847.08** | 0.308 | 0.415 | 0.529 | 0.480 | 0.549 | 0.520 | 0.417 | 0.428 | 0.476 | 0.533 |
| **1851.9** | 0.834 | 0.899 | 0.689 | 0.691 | 0.534 | 0.481 | 0.800 | 0.838 | 0.260 | 0.091 | **1851.9** | 0.648 | 0.662 | 0.584 | 0.569 | 0.513 | 0.516 | 0.651 | 0.610 | 0.353 | 0.370 |
| **1864.11** | 0.287 | 0.402 | 0.543 | 0.433 | 0.535 | 0.468 | 0.491 | 0.407 | 0.472 | 0.658 | **1864.11** | 0.309 | 0.427 | 0.517 | 0.464 | 0.521 | 0.495 | 0.437 | 0.441 | 0.510 | 0.563 |
| **1867.87** | 0.803 | 0.879 | 0.682 | 0.677 | 0.546 | 0.484 | 0.783 | 0.836 | 0.270 | 0.106 | **1867.87** | 0.779 | 0.842 | 0.721 | 0.688 | 0.456 | 0.474 | 0.861 | 0.833 | 0.218 | 0.222 |
| **1873.88** | 0.747 | 0.766 | 0.601 | 0.622 | 0.558 | 0.521 | 0.641 | 0.705 | 0.339 | 0.205 | **1873.88** | 0.678 | 0.671 | 0.556 | 0.534 | 0.486 | 0.504 | 0.612 | 0.559 | 0.410 | 0.421 |
| **1924.91** | 0.874 | 0.864 | 0.679 | 0.695 | 0.520 | 0.474 | 0.808 | 0.866 | 0.256 | 0.095 | **1924.91** | 0.967 | 0.959 | 0.714 | 0.673 | 0.419 | 0.443 | 0.908 | 0.856 | 0.221 | 0.235 |
| **1942.91** | 0.851 | 0.877 | 0.803 | 0.751 | 0.529 | 0.464 | 0.936 | 0.926 | 0.157 | 0.032 | **1942.91** | 0.891 | 0.942 | 0.830 | 0.805 | 0.455 | 0.470 | 0.948 | 0.937 | 0.086 | 0.086 |
| **1950.91** | 0.819 | 0.824 | 0.695 | 0.687 | 0.531 | 0.484 | 0.816 | 0.862 | 0.256 | 0.106 | **1950.91** | 0.833 | 0.851 | 0.707 | 0.688 | 0.453 | 0.466 | 0.848 | 0.828 | 0.229 | 0.229 |
| **1964.9** | 0.845 | 0.873 | 0.683 | 0.706 | 0.523 | 0.464 | 0.821 | 0.885 | 0.265 | 0.086 | **1964.9** | 0.626 | 0.620 | 0.574 | 0.563 | 0.492 | 0.509 | 0.632 | 0.586 | 0.392 | 0.396 |
| **1972.89** | 0.887 | 0.877 | 0.702 | 0.681 | 0.505 | 0.470 | 0.852 | 0.884 | 0.240 | 0.098 | **1972.89** | 0.898 | 0.895 | 0.658 | 0.623 | 0.441 | 0.467 | 0.853 | 0.779 | 0.271 | 0.285 |
| **1976.01** | 0.880 | 0.920 | 0.550 | 0.580 | 0.481 | 0.480 | 0.624 | 0.744 | 0.409 | 0.212 | **1976.01** | 0.926 | 0.911 | 0.646 | 0.608 | 0.412 | 0.423 | 0.849 | 0.801 | 0.308 | 0.329 |
| **1980.87** | 0.829 | 0.841 | 0.657 | 0.681 | 0.522 | 0.472 | 0.784 | 0.875 | 0.294 | 0.112 | **1980.87** | 0.810 | 0.841 | 0.662 | 0.626 | 0.443 | 0.457 | 0.809 | 0.774 | 0.291 | 0.303 |
| **1986.89** | 0.785 | 0.787 | 0.615 | 0.656 | 0.548 | 0.497 | 0.677 | 0.794 | 0.326 | 0.159 | **1986.89** | 0.725 | 0.707 | 0.584 | 0.561 | 0.469 | 0.492 | 0.686 | 0.625 | 0.377 | 0.386 |
| **2002.86** | 0.780 | 0.766 | 0.590 | 0.616 | 0.522 | 0.493 | 0.667 | 0.768 | 0.363 | 0.212 | **2002.86** | 0.682 | 0.695 | 0.548 | 0.533 | 0.466 | 0.478 | 0.611 | 0.583 | 0.437 | 0.435 |
| **2010.96** | 0.793 | 0.845 | 0.589 | 0.602 | 0.545 | 0.504 | 0.648 | 0.745 | 0.351 | 0.194 | **2010.96** | 0.830 | 0.838 | 0.596 | 0.597 | 0.453 | 0.473 | 0.750 | 0.723 | 0.349 | 0.324 |
| **2080** | 0.793 | 0.824 | 0.757 | 0.679 | 0.511 | 0.468 | 0.907 | 0.843 | 0.218 | 0.138 | **2080** | 0.867 | 0.889 | 0.775 | 0.733 | 0.453 | 0.465 | 0.917 | 0.887 | 0.147 | 0.172 |
| **2093.06** | 0.830 | 0.832 | 0.717 | 0.737 | 0.504 | 0.443 | 0.837 | 0.905 | 0.259 | 0.089 | **2093.06** | 0.875 | 0.882 | 0.727 | 0.705 | 0.438 | 0.453 | 0.883 | 0.845 | 0.211 | 0.218 |
| **2095.04** | 0.869 | 0.873 | 0.818 | 0.741 | 0.518 | 0.472 | 0.943 | 0.908 | 0.148 | 0.041 | **2095.04** | 0.918 | 0.949 | 0.810 | 0.784 | 0.477 | 0.496 | 0.945 | 0.917 | 0.079 | 0.082 |
| **2132.99** | 0.801 | 0.813 | 0.684 | 0.669 | 0.531 | 0.493 | 0.793 | 0.818 | 0.271 | 0.134 | **2132.99** | 0.859 | 0.880 | 0.676 | 0.646 | 0.454 | 0.470 | 0.848 | 0.786 | 0.251 | 0.265 |
| **2134** | 0.858 | 0.870 | 0.698 | 0.650 | 0.522 | 0.518 | 0.810 | 0.786 | 0.247 | 0.116 | **2134** | 0.921 | 0.946 | 0.745 | 0.698 | 0.469 | 0.490 | 0.903 | 0.839 | 0.153 | 0.178 |
| **2215.01** | 0.811 | 0.789 | 0.563 | 0.575 | 0.530 | 0.529 | 0.604 | 0.684 | 0.369 | 0.236 | **2215.01** | 0.912 | 0.870 | 0.546 | 0.537 | 0.424 | 0.447 | 0.733 | 0.650 | 0.414 | 0.409 |
| **2437.09** | 0.864 | 0.865 | 0.666 | 0.649 | 0.478 | 0.459 | 0.815 | 0.827 | 0.305 | 0.163 | **2437.09** | 0.878 | 0.859 | 0.689 | 0.642 | 0.428 | 0.450 | 0.853 | 0.771 | 0.261 | 0.295 |
| **2459.08** | 0.842 | 0.826 | 0.598 | 0.597 | 0.480 | 0.474 | 0.718 | 0.740 | 0.366 | 0.239 | **2459.08** | 0.741 | 0.708 | 0.521 | 0.508 | 0.446 | 0.453 | 0.621 | 0.558 | 0.468 | 0.483 |
