## Supplemental Table 5 for "Multimodal Imaging of the Cellular and Extracellular Microenvironment on the Same Formalin-Fixed Paraffin-Embedded Tissue Section"

**Supplemental Table 5: MALDI IMS peptides with decreased maximum peak intensity after multiplexed immunofluorescence.** Calibrated MALDI IMS m/z values and corresponding maximum peak intensity without (MxIF-) or with (MxIF+) prior multiplexed immunofluorescence and corresponding percent change from MxIF- to MxIF+. Colocalization coefficients by Pearson's Correlation Coefficient in SCiLS show how well (max = 1.00) m/z values co-localize to representative m/z value (m/z = 1013.606) shown in main figure. If m/z values were identifiable within 8 ppm by LC-TIMS-MS/MS reference library, calculated protonated peptide mass, mass error, modified peptide sequence, peptide start, peptide end, gene name and protein description are listed.

| Calibrated m/z | MxIF- Max Peak Intensity | MxIF+ Max Peak Intensity | Percent Change | Colocalization coefficient with m/z = 1013.606 | Identified within 8 ppm | Calculated Protonated Peptide Mass | Mass Error | Modified Peptide | Peptide Start | Peptide End | Gene | Protein Description |
|---|---|---|---|---|---|---|---|---|---|---|---|---|
| 789.331 | 74.30 | 24.20 | -67% | 0.37 | Yes | 789.3373 | -7.85 | GEQGPSGAS | 1127 | 1135 | COL1A1 | Collagen alpha-1(I) chain |
| 790.413 | 232.47 | 177.51 | -24% | -0.05 | No | Unidentified | | | | | | |
| 957.469 | 196.65 | 189.00 | -4% | 0.00 | Yes | 957.471 | -1.88 | PVEEAPKGM | 161 | 169 | ARHGDIA | Rho GDP-dissociation inhibitor 1 |
| 1013.606 | 200.14 | 8.39 | -96% | 1.00 | Yes | 1013.599 | 6.61 | LVAASQAALGL | 599 | 609 | ALB | Albumin |
| 1151.592 | 372.71 | 302.18 | -19% | 0.08 | No | Unidentified | | | | | | |
| 1346.612 | 598.32 | 540.70 | -10% | -0.06 | No | Unidentified | | | | | | |
| 1622.946 | 89.90 | 5.46 | -94% | 0.84 | No | Unidentified | | | | | | |
| 1847.066 | 105.21 | 6.03 | -94% | 0.87 | No | Unidentified | | | | | | |
| 1864.091 | 115.51 | 6.00 | -95% | 0.86 | Yes | 1864.0902 | 0.59 | LLSPGSVDPLTRLVLVNA | 149 | 166 | SERPINB6 | Serpin B6 |
